# DyMoTree decodes early cell state transitions and drivers from single-cell transcriptomes using a tree-structured neural network

**DOI:** 10.64898/2026.06.09.731114

**Authors:** Jiayi Wang, Rufeng Li, Chen Guo, Min Qiang, Shuai Wang, Genhui Wang, Kangsheng Tu, Yungang Xu

**Affiliations:** Department of Cell Biology and Genetics, School of Basic Medical Sciences, Xi’an Jiaotong University Health Science Center, Xi’an, Shaanxi 710061, China; Department of Hepatobiliary Surgery, The First Affiliated Hospital of Xi’an Jiaotong University, Xi’an, Shaanxi, 710061, China

**Keywords:** single-cell transcriptomics, cell fate inference, lineage-resolved modeling, cell state transitions, tree-structured neural network, driver gene discovery

## Abstract

Inferring early cell fate from single-cell RNA-sequencing data is essential for identifying cellular origins and fate plasticity in development and disease. However, existing methods often fail to exploit tree-structured lineage trajectories, limiting the accuracy and interpretability of fate mapping. Here we present DyMoTree, a computational framework that models cell fate decisions as nonlinear mappings between progenitor and terminal cell states under explicit lineage constraints. By integrating lineage graphs with a tree-structured neural architecture, DyMoTree learns lineage-resolved cell-state transition maps from single-cell transcriptomes, enabling robust inference of early fate bias and identification of fate-specific progenitor substates and driver genes. Across simulations, lineage-tracing experiments, and in vivo systems, DyMoTree outperformed existing methods in resolving early fate biases. Applications to mouse embryogenesis, lung adenocarcinoma progression, and CAR-T immunotherapy revealed regulatory programs underlying developmental and disease-associated transitions. DyMoTree provides a general framework for modeling lineage-resolved cell-state dynamics underlying development and disease progression.

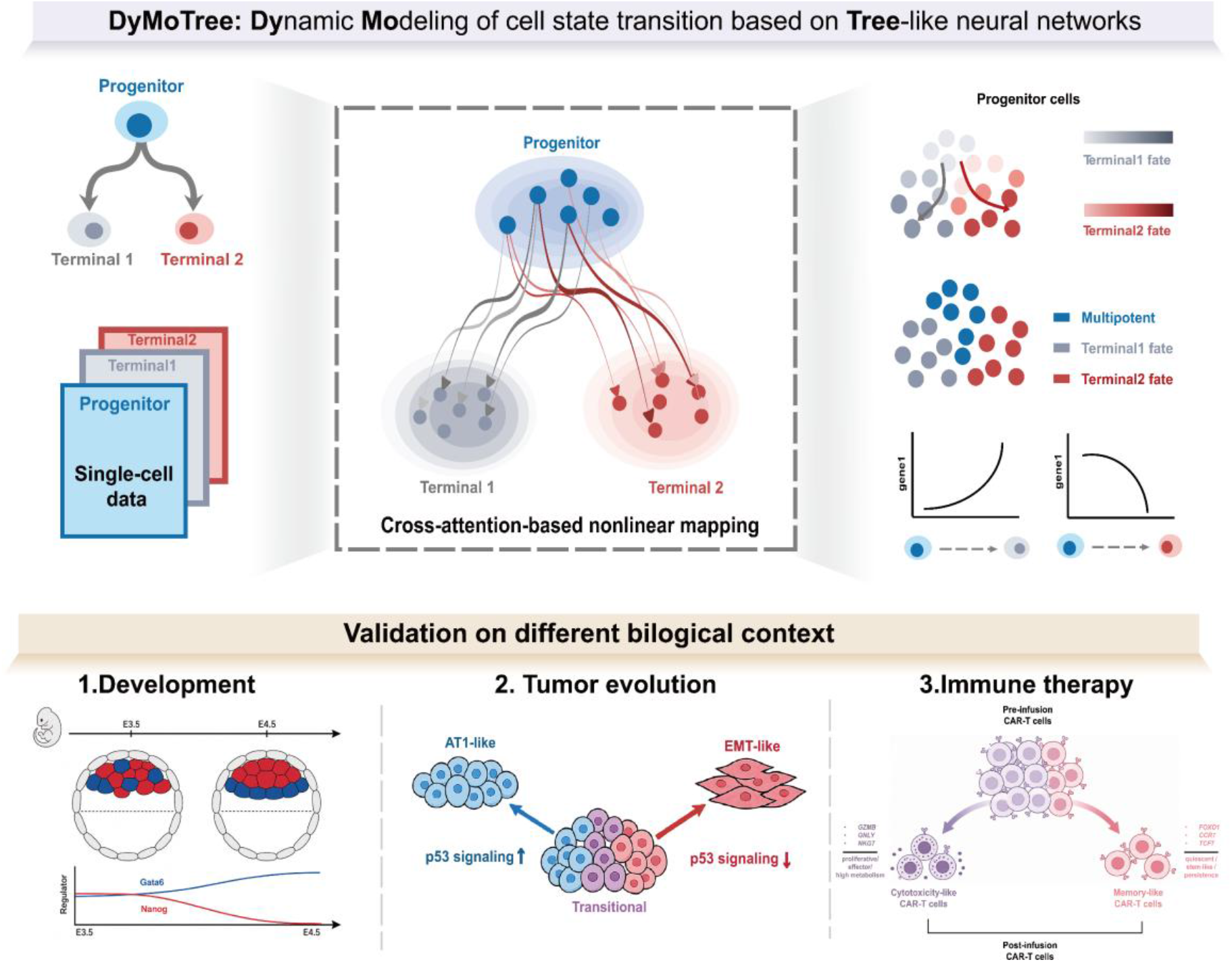

## Introduction

Understanding the fundamental principles that govern cellular origins and fate plasticity is central to deciphering biological processes across diverse contexts, including development, regeneration, cellular reprogramming, and disease progression^1-3^. Cell fate decisions lie at the core of these processes and are essential for shaping biological systems. In many biological systems, cell differentiation follows an organized lineage structure that can be experimentally established or computationally inferred, providing a structural scaffold that links progenitor populations to their terminal states^4^. In many of these settings, particularly during disease progression, fate commitment precedes overt transcriptional state shifts^5, 6^. Resolving such early fate decisions is therefore critical for understanding how initial gene expression programs drive long-term fate outcomes in both physiological and pathogenic settings.

Single-cell RNA sequencing (scRNA-seq) has revolutionized the study of cellular heterogeneity and differentiation across complex biological systems^7, 8^. Despite its transformative impact, the inherently static nature of single-cell measurements precludes direct temporal tracking, rendering the inference of cellular dynamics and early fate biases fundamentally challenging^9^. Single-cell lineage-tracing approaches partially alleviate this limitation by combining scRNA-seq with heritable barcodes to enable the recovery of clonal relationships. However, their applicability is predominantly confined to *in vitro* systems and model organisms, and is further constrained by incomplete labeling, barcode loss, and limited scalability, particularly in disease-relevant and *in vivo* contexts^4, 6, 8^. This disconnect underscores the need for computational frameworks capable of inferring early cell fate bias and its molecular drivers directly from scRNA-seq data within a structured lineage hierarchy, without relying on additional experimental information.

Building on this need, a range of computational approaches has been developed to infer cell fate from single-cell transcriptomic data. Transcriptomic similarity-based approaches link progenitor cells to terminal cell states according to gene expression similarity and infer fate probabilities using, for example, machine learning models trained on well-annotated terminal cell types^10, 11^. However, their performance depends heavily on accurate endpoint annotation, and when cellular states are continuous or lineage boundaries are ambiguous, these approaches may yield unstable estimates of early fate bias. Other efforts have adopted Markov state models to represent differentiation as probabilistic transitions between cells^12-15^. By incorporating additional directional information, such as pseudotime or RNA velocity, these approaches construct more reliable directed transition matrices on the cell state manifold and propagate local transition probabilities to estimate fate probabilities. Although effective in densely sampled and relatively smooth differentiation landscapes, the performance of these methods is highly context dependent and degrades under noisy or sparsely sampled conditions, such as disease progression. More recently, optimal transport (OT)-based frameworks have been introduced to model cell state transitions by solving minimal transport problems in gene expression space^16-19^. But the inferred transport plan is strictly determined by a predefined cost matrix, which is typically defined using simple distance-based metrics in gene expression space. Furthermore, the linear transport formulation underlying OT limits its ability to capture complex, nonlinear relationships that characterize cell state transitions across dynamic biological systems.

To fill this gap, we introduce DyMoTree, a computational framework that models cell state transitions as nonlinear information flow across lineage levels. Unlike traditional approaches that propagate local transitions across cell state manifolds^12-15, 20, 21^, DyMoTree learns conditional mappings from progenitor states to terminal states, constrained by an explicit lineage structure. By integrating lineage graphs, derived from transcriptomic similarity, with a tree-structured neural architecture, DyMoTree captures the complex nonlinear relationships from progenitor state to terminal states. This formulation enables three key tasks, including robust inference of early progenitor fate biases, identification of fate-specific progenitor substates, and discovery of lineage-specific driver genes. In contrast to methods relying on RNA velocity, metabolic labeling, or experimental lineage tracing^12, 14, 15, 19, 22^, DyMoTree requires only single-cell gene expression profiles and an explicitly defined or computationally inferred lineage structure. DyMoTree replaces cost-constrained transport and diffusion-based propagation with data-driven, nonlinear lineage-level coupling, thereby enhancing both the accuracy and interpretability of cell fate inference.

We systematically evaluated DyMoTree across multiple levels of data, including simulated datasets, *in vitro* single-cell lineage-tracing datasets, and *in vivo* scRNA-seq datasets combined with single-cell T cell receptor (TCR) sequencing data. Across all experimental conditions, DyMoTree accurately recovered early fate biases in progenitor states. We further applied DyMoTree to early mouse embryogenesis datasets, where we successfully reconstructed dynamic expression trends of key regulators during the second cell-fate decision. In lung adenocarcinoma (LUAD) tumor progression and B-cell acute lymphoblastic leukemia (B-ALL) CAR-T immunotherapy datasets, DyMoTree inferred biologically interpretable transition patterns and recovered molecular mechanisms underlying different lineage biases. Together, these results establish DyMoTree as a general framework, which reframes cell fate prediction as the learning of conditional nonlinear mappings across hierarchical lineage levels, for modeling lineage-resolved cell state transitions directly from single-cell transcriptomic data. Specifically, its ability to robustly resolve early cell fate biases and uncover lineage-specific regulatory programs holds great promise for deciphering developmental processes and facilitating early detection of pathogenic changes in disease settings such as tumor progression, thereby laying a solid foundation for advancing both basic biological research and clinical translational applications.

## Results

### Problem formulation for modeling lineage-level cell state transitions

Unlike traditional trajectory inference approaches that rely on local continuity assumptions and model gradual state evolution along differentiation paths^20, 21^, lineage-resolved fate inference instead focuses on linking progenitor cells to terminal states^16-19, 22^. Following this paradigm, we formulate lineage-resolved transitions as the direct inference of cell state mapping relationships from progenitor to terminal cells. Most existing cell fate modeling frameworks depend on a core assumption that cell state transitions follow a similarity principle, that within the cell state manifold, a cell tends to transition toward the direction defined by the most similar cells^23^. We adopt the same assumption to model lineage-level cell-cell state mappings.

In practice, many studies quantify lineage-resolved or cross-timepoint cell state similarity by computing Euclidean distances in the cell state space or by measuring shortest-path distances on a KNN graph constructed over the cell state manifold^16, 18, 19^. To more precisely characterize lineage-resolved state transitions, we combine graph-based shortest-path distances with a linear kernel term in gene expression space to quantify progenitor-terminal similarity (**Fig. 1a, Methods**). First, we construct an undirected K-nearest neighbor (KNN) graph within the cell state manifold underlying the known lineage structure, where each node represents the observed transcriptomic profile of a single cell, and edges connect the most similar cells based on gene expression. We then compute shortest-path distances on this graph to represent structural reachability on the manifold. This geometric distance reflects the minimal transition cost required for a progenitor cell to move toward a specified terminal cell along the cell state manifold. Due to the equidistant or near-equidistant conditions, the shortest-path distance alone cannot adequately capture transition directionality. To address this limitation, we incorporate an additional linear kernel term to quantify the geometric alignment between cell state vectors in gene expression space. The resulting composite similarity matrix provides a refined representation between each progenitor cell and potential terminal cells.

**Fig. 1.**
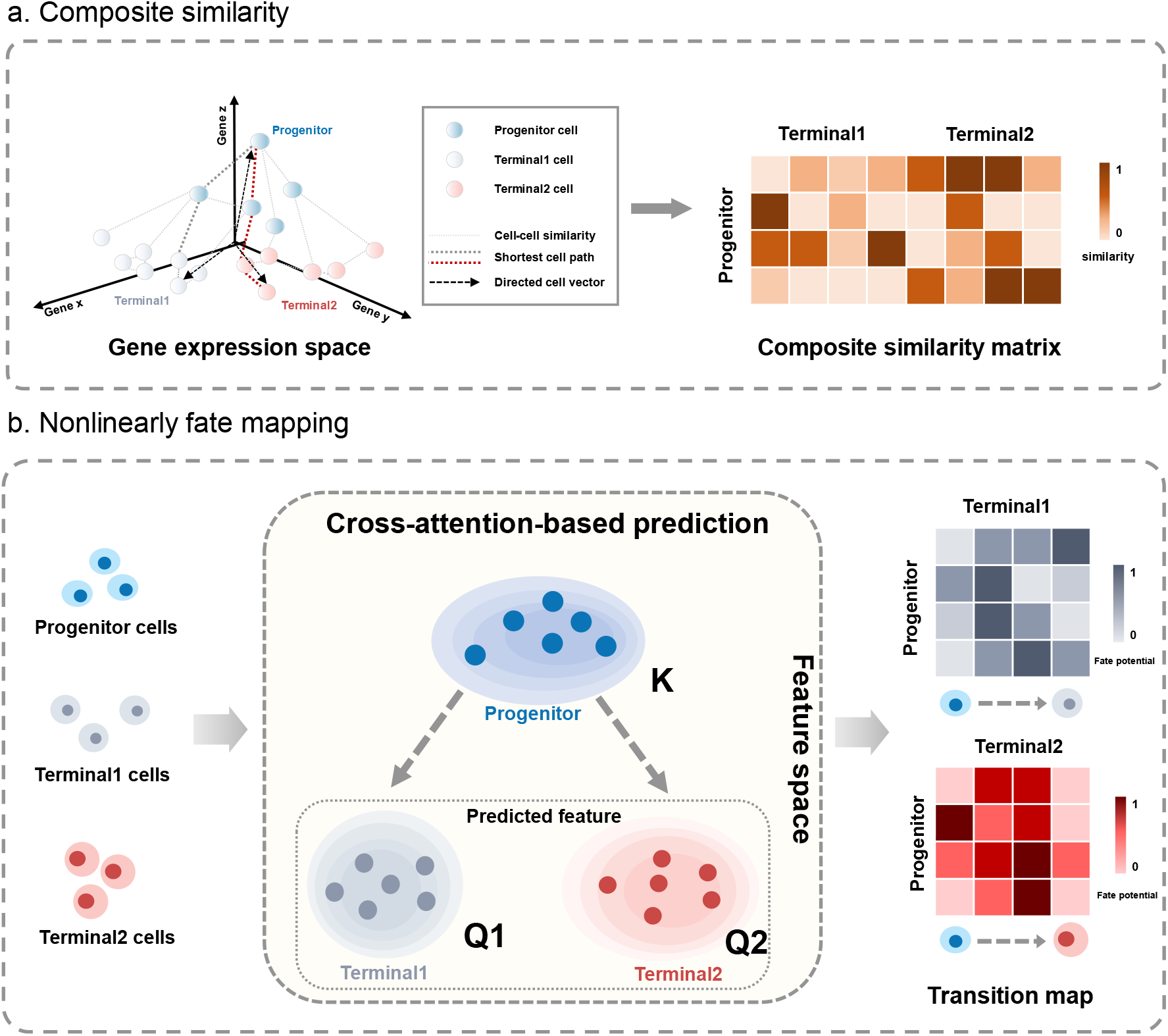
Computational formulation of lineage-level cell state transitions. **(a)** Composite metric for measuring progenitor-terminal similarity. Progenitor–terminal relationships are quantified by integrating two complementary measures: structural reachability, represented by shortest-path distances on a KNN graph in gene expression space that approximate transition cost along the cell state manifold, and geometric alignment, represented by directed cell vectors that capture directional similarity between cell states. Together, these components define a composite similarity metric that refines lineage-level relationships beyond simple Euclidean proximity. **(b)** Nonlinear fate mapping under lineage graph constraints. Progenitor and terminal cell embeddings are connected through a cross-attention-based prediction framework, in which progenitor states provide contextual information to predict terminal representations. The resulting attention weights form a transition map that quantifies lineage-resolved progenitor-to-terminal relationships.

Nevertheless, as a static and linear construct, this similarity matrix remains insufficient for modeling dynamic lineage-level transitions. Rather than directly solving an optimal transport problem to obtain a coupling matrix, we formulate lineage-resolved cell state transitions as a nonlinear mapping problem and learn this mapping through neural networks (**Fig. 1b, Methods**). Intuitively, the mapping from progenitor cells to terminal cells can be viewed as a cross-context prediction task, in which progenitor cell states provide the context for predicting terminal cell features. Inspired by cross-attention mechanisms widely used in deep learning to model interactions between distinct entity sets^24^, we can learn a progenitor-terminal transition map through a cross-attention-based architecture. In this formulation, progenitor cell embeddings serve as the conditioning context for predicting terminal cell representations, and the resulting attention weights quantify the contribution of each progenitor cell to the predicted terminal states.

To regularize this nonlinear mapping and improve the stability of the cross-attention learning, we adopt a graph reconstruction task and introduce a lineage graph to guide the learning process **(Methods, Extended Data Fig. 1)**. This graph consists of two components, including an intra-state graph based on Euclidean distance to capture local relationships within each cell state, and an inter-state graph that models potential state transition relationships between progenitor and terminal states. Notably, the inter-state graph is constructed using composite similarity and provides the primary lineage-level constraints. During training, the predicted terminal embeddings are encouraged to respect the connectivity structure encoded in the lineage graph. This structural guidance helps the cross-attention mechanism learn more robust and biologically meaningful relationships from progenitor to terminal cells. Consequently, the learned attention weights reflect lineage-level coupling, allowing the attention matrix to serve as an interpretable fate map for downstream analysis.

### Overview of the DyMoTree framework

We developed DyMoTree, an integrated computational framework for inferring early cell fate biases that captures lineage-resolved nonlinear cell state transitions by deep neural networks **(Fig. 2)**. DyMoTree takes as input a known lineage differentiation structure and single-cell transcriptomic data **(Fig. 2a)**, without requiring additional biological priors such as lineage tracing, RNA velocity, or temporal information^25-28^. By learning a lineage-resolved cell state transition map from progenitor to terminal cells for each lineage pair **(Fig. 2b)**, the framework quantifies progenitor fate potentials and enables identification of fate-specific progenitor substates and lineage-specific driver genes **(Fig. 2c)**.

**Fig. 2.**
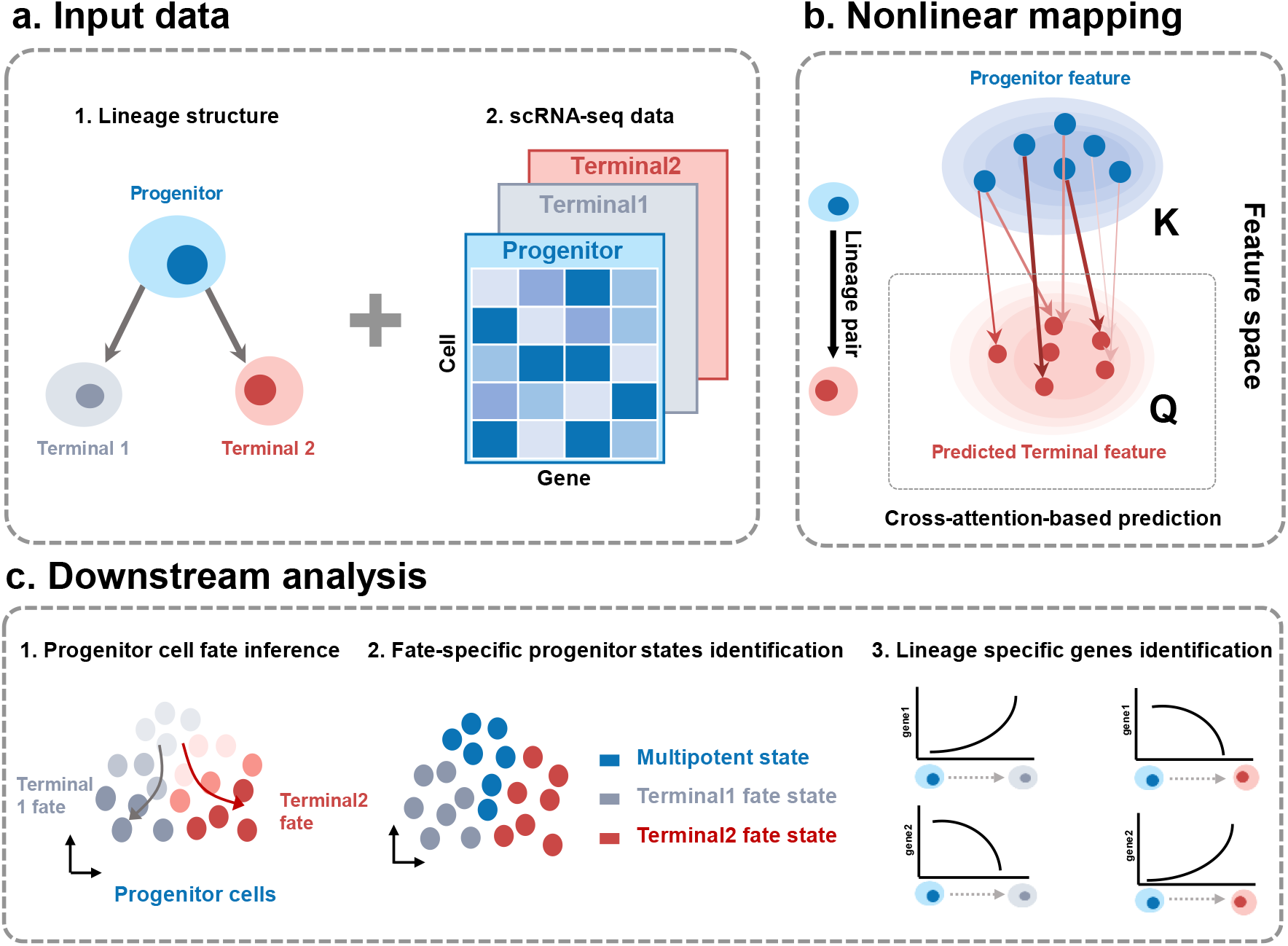
The DyMoTree framework for modeling lineage-resolved cell-state transitions. **(a)** DyMoTree takes a known lineage differentiation structure and gene expression profiles derived from the scRNA-seq data of a lineage structure as inputs. **(b)** Nonlinear fate mapping under a lineage pair. Terminal cell features are predicted from progenitor features through a cross-attention–based prediction framework, in which progenitor states serve as Key and terminal states serve as Query. **(c)** Downstream analyses enabled by DyMoTree include: (1) cell fate inference, which quantifies progenitor cell fate potential toward terminal states; (2) identification of fate-specific progenitor states; and (3) characterization of lineage-specific genes.

At its core, DyMoTree implements a lineage-structured encoder-decoder neural network **(Methods, Extended Data Figs. 2**–**3)**. The encoder comprises CellModules, which embed intra-state transitions using a two-layer attention-based graph neural network^29^, and LineageModules, which model inter-state transitions across the lineage. LineageModules employ a cross-attention mechanism^30^ to quantify each progenitor’s contribution to terminal cells, producing a cell-state transition map that captures nonlinear transition relationships and predicts terminal cell features from progenitor cell features. The training process is supervised through reconstruction of intra- and inter-state transition graphs, which guides the encoder to learn biologically meaningful lineage-resolved mappings under the constraint of the lineage graph.

To stabilize optimization and promote robust nonlinear mappings, DyMoTree adopts a two-stage pretraining strategy **(Methods, Extended Data Fig. 4)**, which is applied specifically to CellModules to establish robust transition representations before full model training. In the first stage, each CellModule is independently pretrained on its corresponding intra-state transition graph, enabling it to capture local transition topologies within each cell state along the differentiation hierarchy. In the second stage, CellModules are further optimized by jointly reconstructing intra- and inter-state transition graphs across lineage pairs, allowing them to incorporate both local and lineage-resolved transition patterns. This staged pretraining equips CellModules with hierarchical transition knowledge and guides the LineageModules to more effectively capture progenitor-to-terminal fate relationships while reducing spurious correlations.

To quantify progenitor fate potentials, we aggregate the nonlinear transition scores from the transition map at the progenitor cell level, calculating the likelihood of each progenitor differentiating toward each terminal state **(Methods)**. Cells in biological systems rarely distribute uniformly in state space; instead, they form structured manifolds shaped by the underlying gene regulatory program^31,32^. Using the fate potentials inferred by DyMoTree, we approximate a fate manifold for progenitor cells based on fate potential–associated genes **(Methods)**, where the manifold’s topology reflects lineage-specific fate patterns, and distinct branches correspond to transitions toward different terminal cell types. We use archetype analysis on this manifold to identify fate-specific progenitor substates^31-33^ **(Methods, Extended Data Fig. 5)**. Finally, by integrating progenitor fate potentials with sparse modeling and network-based smoothing, DyMoTree uncovers lineage-specific driver genes associated with differentiation^34,35^. **(Methods, Extended Data Fig. 6)**.

### DyMoTree outperforms state-of-the-art methods in resolving cell fate choices during cell state transitions

We first assessed the reliability of DyMoTree by using simulated single-cell transcriptomic profiles following a bifurcating lineage in which stem cells progress continuously toward two terminal states (**Methods, Fig. 3a–b**)^36^. DyMoTree successfully identified clear fate biases within the stem cell state **(Fig. 3c)**. The simulation also specified three fate-biased substates within the stem population, including stemness, Child1-biased, and Child2-biased **(Fig. 3d)**. Leveraging these inferred potentials, DyMoTree effectively delineated the three fate-biased stem substates in a manner consistent with the simulation design **(Fig. 3e)**.

**Fig. 3.**
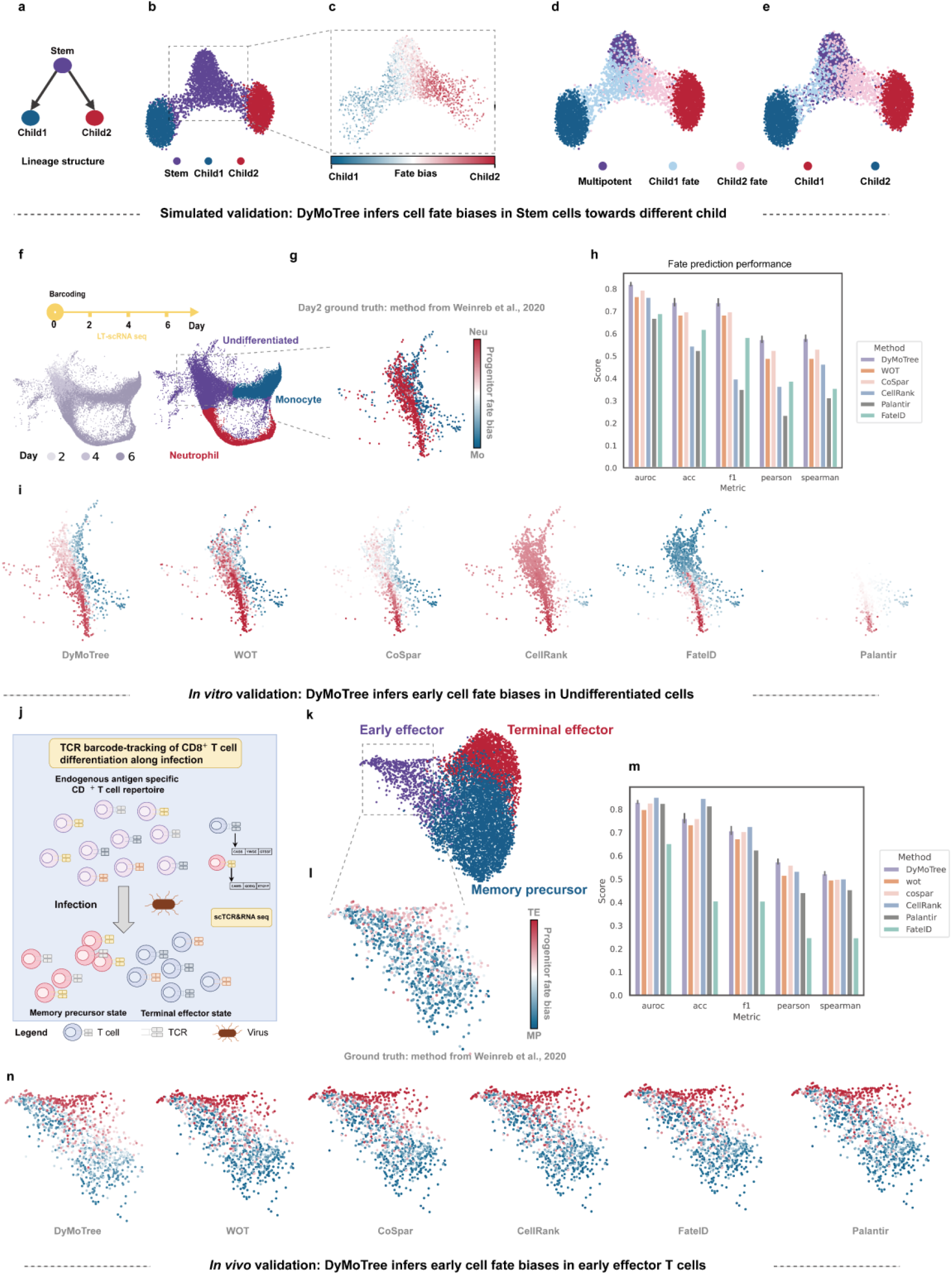
Benchmarking DyMoTree on early cell fate inference. **(a)**Simulated lineage differentiation structure from the Stem state toward Child1 and Child2 states. **(b)**UMAP of the Splatter-simulated single-cell transcriptomic data. Colors correspond to cell state annotation. **(c)** DyMoTree inferred progenitor cell fate bias in the Stem state. **(d-e)** Splatter-simulated (left) and DyMoTree-identified (right) fate-specific cell states. **(f)** Experimental design of single-cell lineage-tracing system (top) and SPRING visualization of the hematopoiesis dataset from Weinreb et al. Colors correspond to sampling time point and cell state annotation from the original study. **(g)** Ground truth cell fate bias toward neutrophil (red) and monocyte (blue) in day 2 lineage-barcoded HSPCs using the original method from Weinreb et al. **(h)** Comparison of DyMoTree’s performance on early cell fate bias inference with five approaches. **(i)** SPRING visualization of day 2 lineage-barcoded HSPCs. Color denotes cell fate bias toward neutrophil (red) and monocyte (blue) in day 2 HSPCs inferred by different approaches. **(j)** Conceptual illustration of CD8^+^ T cell-specific differentiation during infection. **(k)** UMAP of CD8^+^ T cell-specific differentiation process. Colors corresponding to cell state annotation followed by the original study. **(l)** Ground truth cell fate bias toward Terminal effector (TE) state (red) and memory precursor (MP) state (blue) in Early effector (EE) state using the original method from Weinreb et al. **(m)** Comparison of DyMoTree’s performance on early cell fate bias inference with five different approaches. **(n)** UMAP of Early effector state. Colors denote cell fate bias toward TE (red) and MP (blue) in EE state inferred by different approaches.

To assess DyMoTree under realistic biological conditions, we evaluated its performance on a lineage-tracing dataset of mouse hematopoiesis generated using the LARRY barcoding system, which provides experimentally derived fate biases as a stringent ground truth for early cell fate inference^6^ **(Fig. 3f–g)**. On this *in vitro* system, DyMoTree consistently achieved strong performance across multiple evaluation metrics, demonstrating high prediction accuracy and close concordance with experimentally observed fate biases **(Fig. 3h)**. Visualization of inferred fate biases further showed that DyMoTree delineates clear and coherent fate boundaries among early hematopoietic progenitor cells, effectively separating monocyte and neutrophil trajectories before overt transcriptional divergence **(Fig. 3i)**. Compared with existing approaches^10,12,14,18,19^, DyMoTree produces more structured and consistent fate-bias patterns at early progenitor stages. These results demonstrate that DyMoTree accurately captures nonlinear fate bias from early progenitor states and achieves robust performance on experimentally grounded datasets.

To further assess DyMoTree in more complex biological settings, we applied it to a CD8^+^ T-cell differentiation dataset during acute viral infection in mice, which integrates scRNA-seq with single-cell TCR sequencing and captures dynamic *in vivo* fate transitions^38, 39^ **(Fig. 3j)**. This system introduces additional challenges for fate inference due to heterogeneous microenvironmental influences and increased cellular variability. Using TCR clonotypes as endogenous lineage barcodes, this dataset provides an experimentally grounded reference for evaluating fate bias in a physiologically relevant context **(Fig. 3k–l)**. On this dataset, DyMoTree maintained strong and consistent performance across multiple evaluation metrics, demonstrating high agreement with lineage-informed fate biases **(Fig. 3m)**. In particular, DyMoTree effectively resolved fate potentials at early progenitor stages, enabling clear identification of cell fate trajectories toward terminal-effector and memory lineages, even among less differentiated cell populations **(Fig. 3n)**. Together, these results indicate that DyMoTree generalizes well beyond *in vitro* systems and reliably captures nonlinear fate relationships under physiologically relevant conditions.

Notably, OT-based approaches^18,19^ (WOT and CoSpar) showed strong performance in this setting, consistent with our motivation that directly modeling cell fate mapping relationships can improve fate inference **(Fig. 3h–i, m–n)**. We further investigated the performance of directly computing a lineage-resolved composite similarity matrix and leveraging it as a basis for constructing a lineage graph to inform DyMoTree’s nonlinear fate mapping (**Supplementary Figs. 1–4**). The results demonstrate that incorporating neural networks significantly enhances early progenitor fate inference compared with linear computations. DyMoTree is thus able to learn biologically meaningful mappings of cell state transitions. Additionally, ablation experiments indicate that the adoption of composite similarity and the two-stage pretraining strategy both contribute to improved stability and robustness of DyMoTree’s inference (**Supplementary Figs. 5–6**). Overall, these results indicate that DyMoTree provides reliable and robust performance on the inference of early fate bias by learning a nonlinear, lineage-level coupling between progenitor and terminal states.

In addition, we evaluated DyMoTree’s reliability in identifying fate-specific progenitor states using the lineage-tracing datasets. Among day 2 HSPCs, three distinct subpopulations were identified, corresponding to multipotent, monocyte-fate, and neutrophil-fate states (**Extended Data Fig. 7**). Compared with conventional state-identification methods^21, 40^, DyMoTree identified a greater number of fate-biased cells, particularly within the monocyte and neutrophil branches, most of which carried lineage barcodes consistent with the predicted fates, supporting its ability to resolve early differentiation bias (**Extended Data Fig. 7a**). To further validate the identified fate-specific states, we quantified the stemness score using the entropy of DyMoTree-derived fate potential distributions. The resulting entropy values showed a strong positive correlation with CytoTrace-derived stemness score^41^ (**Extended Data Fig. 7b**), supporting the biological relevance of DyMoTree’s fate potential estimates. Consistent with this, the stemness marker gene Cd34 and CytoTrace scores were highest in the multipotent state identified by DyMoTree, whereas classical fate-determining genes displayed lineage-specific expression patterns in the corresponding monocyte- and neutrophil-fate states^6^ (**Extended Data Fig. 7c–e**).

In summary, DyMoTree exhibited robust performance in quantifying early cell fate biases across real datasets under diverse conditions. It also identified biologically consistent, fate-specific progenitor states, providing a reliable computational framework for analyzing the molecular basis of cell fate determination.

### DyMoTree reveals the dynamics of cell fate decisions during ICM bifurcation in early embryogenesis

During the second cell-fate decision in mammalian embryogenesis, the inner cell mass (ICM) of the blastocyst segregates into two distinct lineages, the epiblast (Epi) and the primitive endoderm (PrE)^42, 43^. To demonstrate the application potential of DyMoTree in embryonic developmental contexts, we collected and processed single-cell transcriptomic data of mouse gut endoderm development, extracting all annotated ICM, Epi, and PrE cells from embryonic days E3.5 to E4.5 **(Fig. 4a, Supplementary Fig. 7a**)^44^.

**Fig. 4.**
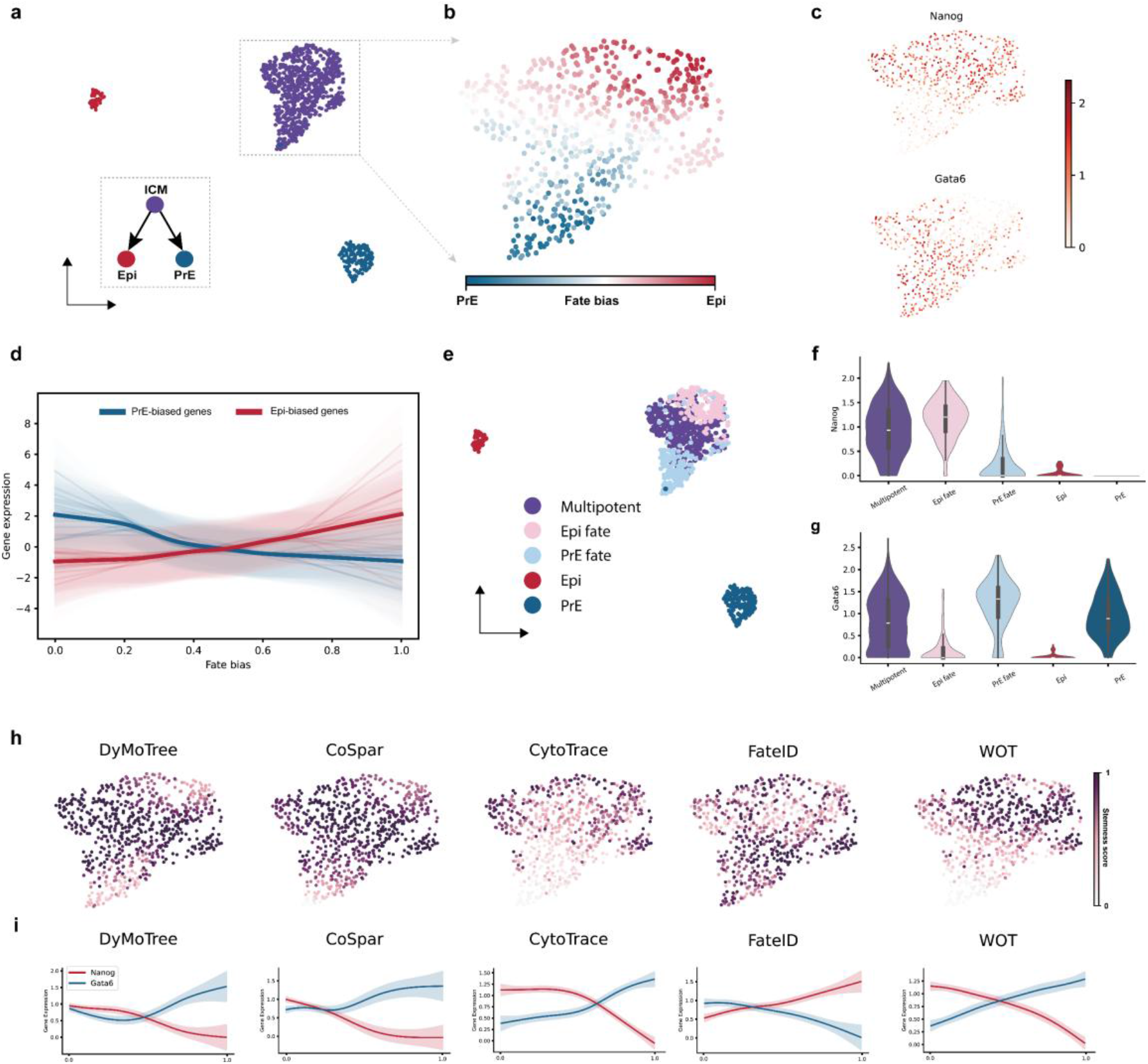
DyMoTree resolves the cell state transition dynamics of ICM bifurcation. **(a)** UMAP illustration of inner cell mass (ICM) differentiating into epiblast (Epi) and primitive endoderm (PrE) state. **(b)** DyMoTree inferred cell fate bias toward Epi (red) and PrE (blue) state in ICM. **(c)** Gene expression of Nanog (top) and Gata6 (bottom) in the ICM state. **(d)** Expression trends of the top 20 contributing genes to Epi-(light red) and PrE-(light blue) fate, respectively, in ICM cells. Cells are sorted along the fate bias inferred by DyMoTree, which corresponds to cell fate potential from PrE-fate (0) to Epi-fate (1). Bold lines represent the mean expression trends of all fate-contributing genes. **(e)** UMAP illustration of Epi, PrE, and fate-specific ICM states identified by DyMoTree. **(f-g)** Violin plots show gene expression patterns of Nanog (top) and Gata6 (bottom) in Epi, PrE, and fate-specific ICM states. **(h)** Comparison of stemness score calculated by cell fate potentials from different approaches. Note that CellRank and Palantir cannot infer cell fate bias in ICM cells **(Supplementary Fig. 9). (i)** Gene expression trends of Nanog and Gata6 in ICM cells. Cells are sorted along differentiation pseudotime defined by the stemness score calculated from different approaches.

Firstly, we applied DyMoTree to infer the fate potentials of ICM cells toward the Epi and PrE lineages and computed the fate bias score for the ICM state. The inferred fate-bias revealed a clear boundary within the ICM cells, suggesting distinct differentiation trajectories toward the two lineages (**Fig. 4b**). This lineage bifurcation is regulated primarily by the transcription factors Nanog and Gata6, which drive the specification of Epi and PrE fates, respectively^42^. In the early ICM, both transcription factors are co-expressed within individual cells; as development progresses, their expression patterns become mutually exclusive, marking the commitment of cells toward either the Epi or PrE lineage^42, 45-47^. In the results, the expression levels of these two regulators exhibited opposing gradients along the corresponding fate potential, confirming that DyMoTree accurately captured the underlying cell-state transition patterns during early embryogenesis (**Fig. 4c, Supplementary Fig. 7b–c**).

Next, we extracted the top 200 genes most correlated with Epi- or PrE-biased fate potentials within ICM cells and then quantified each gene’s contribution to fate potential through our regression-based strategy. We selected the top 20 contributing genes for each lineage direction (**Fig. 4d, Supplementary Fig. 8a–c**). These genes exhibited expression patterns consistent with the inferred fate potentials. For the Epi direction, the highest-contributing genes included canonical Epi differentiation regulators (Nanog, Sox2) and marker genes (Zfp42, Klf2, Klf4)^44, 45^. Interestingly, the PrE-fate related gene Fgf4 showed a positive trend aligning with Epi-fate potential. Previous studies have shown that Fgf4, secreted by Epi cells, can induce neighboring unspecified cells to adopt a PrE identity^48^. This observation suggests that, as a diffusible signaling molecule, Fgf4 promotes PrE differentiation in a non-cell-autonomous manner, highlighting the limitations of current computational approaches that fail to account for cell-cell communication when identifying fate-associated drivers^32, 49-51^. Genes associated with PrE fate potential comprised classical PrE differentiation regulators (Gata6) and marker genes (Gata4, Sox17, Pdgfra, Fgfr2) ^44, 45^. Notably, extracellular matrix (ECM)-related genes, including Lama1, Col4a1, and Serpinh1, were also among the top contributors, suggesting enhanced cell migration and ECM interaction in PrE-biased ICM cells. Functional enrichment analysis further revealed distinct biological programs associated with each fate (**Supplementary Fig. 8d**). Epi-fate-specific genes were enriched in processes such as stem cell population maintenance and chromatin remodeling, consistent with a pluripotent, differentiation-permissive state. In contrast, PrE-fate-associated genes were enriched in the ERK1/2 signaling cascade, in agreement with previous findings that ERK activation drives ICM segregation toward the PrE lineage^52^.

We further identified three fate-specific cell states within ICM cells, corresponding to the multipotent state, PrE-biased state, and Epi-biased state (**Fig. 4e**). These states closely recapitulate the sequential progression of ICM differentiation, from an early mixed state characterized by co-expression of Gata6 and Nanog, to intermediate fate-biased states resembling pre-PrE and pre-Epi states, and ultimately toward fully segregated PrE and Epi lineages. In line with this developmental continuum^42, 46, 47^, Gata6 and Nanog were moderately co-expressed in the multipotent ICM state but exhibited mutually exclusive expression patterns in the two fate-biased populations **(Fig. 4f–g**), confirming the biological accuracy of DyMoTree-inferred fate-specific states.

Having established that DyMoTree accurately identifies fate-specific subpopulations within the ICM, we further examined whether it could also reconstruct the dynamic progression of ICM differentiation. We evaluated the biological significance of the stemness score calculated from cell fate potentials by ordering ICM cells along a differentiation pseudotime defined by the stemness score (**Fig. 4h, Supplementary Fig. 9a–b**). Along this axis, DyMoTree faithfully reconstructed the expression dynamics of Gata6 and Nanog, showing their gradual divergence from an early mixed-expression state to lineage-specific activation patterns. In contrast, other computational methods failed to accurately capture the onset and progression of the fate bifurcation process (**Fig. 4i, Supplementary Fig. 9c–d**). Together with benchmark results from *in vitro* lineage-tracing and *in vivo* CD8^+^ T-cell differentiation datasets, these findings demonstrate that DyMoTree provides a robust and biologically consistent framework for modeling cell state transitions.

### DyMoTree resolves fate potentials of transitional tumor cells during lung adenocarcinoma evolution

Tumor cell-state plasticity plays a key role in disease progression and therapeutic resistance^53^. To demonstrate DyMoTree’s applicability in modeling dynamic cell-state transitions during tumor evolution, we analyzed time-series single-cell transcriptomic data from a genetically engineered mouse model (GEMM) of lung adenocarcinoma (LUAD), spanning the transition from normal alveolar type 2 (AT2) cells to malignant LUAD cells^54^. Previous studies have identified a population of highly plastic transitional tumor cells, characterized by high Trp53 expression, that can differentiate into tumor states with varying degrees of malignancy^55^. The p53 signaling pathway is strongly activated in these cells and has been shown to promote transitions toward low-malignant, AT1-like states.

To investigate whether DyMoTree can resolve cell state transition patterns during tumor progression, we extracted transitional, AT1-like, and EMT-like tumor cell states and constructed a lineage differentiation tree describing transitions from the transitional state toward AT1 and EMT states (**Fig. 5a**). The fate potentials inferred by DyMoTree revealed a clear fate boundary within the transitional tumor cell state, with part of the cells biased toward the low-malignant AT1-like fate and the others toward the high-malignant EMT-like fate (**Fig. 5b**). To verify the reliability of the inferred fate potentials, we measured the activity of gene sets related to p53 signaling, metabolic processes, and cell proliferation (**Fig. 5c, Supplementary Fig. 10a, Supplementary Table 1**)^55^. The p53 signaling activity increased progressively with higher AT1 fate potential, accompanied by reduced metabolic and proliferative activity. Moreover, AT1 cell marker Ager correlated positively with the p53 pathway activity, which rose along the AT1 fate potential, but negatively with the EMT-associated marker Hnf4a, whose expression increased along the EMT fate potential (**Supplementary Fig. 10b–d**). These results indicate that DyMoTree reconstructed biologically coherent fate potentials that align with previously established biological mechanisms during LUAD tumor progression.

**Fig. 5.**
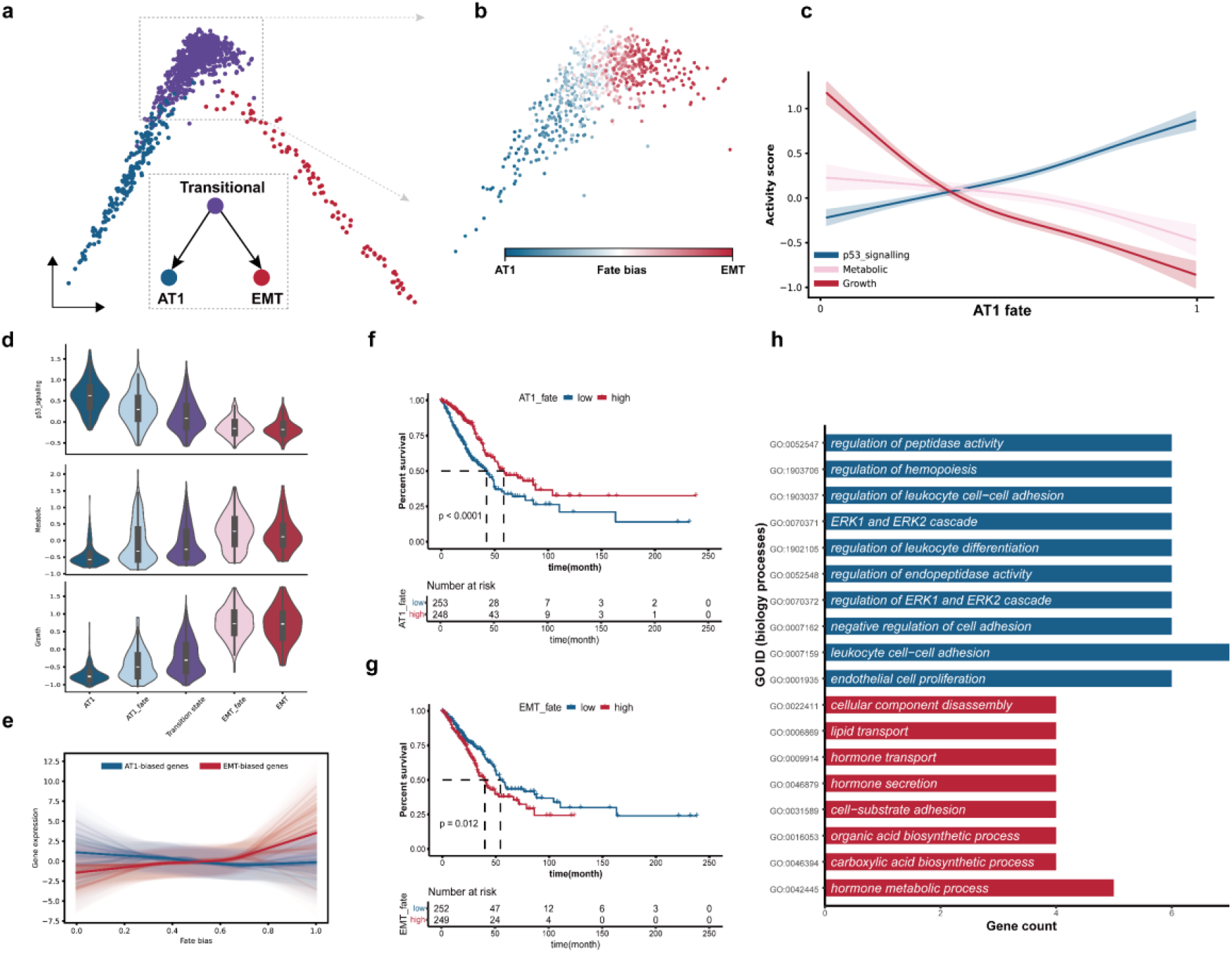
DyMoTree resolves p53-mediated fate decisions and prognostic signatures in lung adenocarcinoma (LUAD) evolution. **(a)** Diffusion map of transitional, AT1-like, and EMT-like tumor cell states. **(b)** DyMoTree inferred cell fate bias toward EMT-(red) and AT1-(blue) state in transitional tumor state. **(c)** Activity trends of p53 signaling pathway (blue), metabolic (pink), and cell growth (red) in the transitional tumor state. Cells are sorted along AT1 fate potential. **(d)** Violin plots show activity of the p53 signaling pathway (top), metabolic (mid), and cell growth (bottom) in AT1, EMT, and fate-specific transitional tumor states. **(e)** Expression trends of the top 30 contributing genes to EMT-(light red) and AT1-(light blue) fate, respectively, in transitional tumor cells. Cells are sorted along the fate bias inferred by DyMoTree, which corresponds to cell fate potential from AT1-fate (0) to EMT-fate (1). Bold lines represent the mean expression trends of all fate-contributing genes. **(f-g)** Kaplan– Meier survival plot of TCGA LUAD cohort stratified by mean expression level of AT1-(top) and EMT-(bottom) fate-specific genes. **(h)** The top enriched GO terms of the AT1-fate specific (blue) and EMT-fate specific (red) genes, respectively.

Building upon the inferred fate potentials, we identified three fate-specific tumor cell states within the transitional state, corresponding to AT1-biased, transition, and EMT-biased states. Trp53 expression was higher in the transition and AT1-biased states than in the EMT-biased state. Ager and Hnf4a were markedly upregulated in the AT1- and EMT-biased states, respectively. (**Supplementary Fig. 11a–d**). Consistent with previous findings, the fate-specific tumor cell states displayed continuous functional activity gradients along the two transition directions (**Fig. 5d**)^55^. As transitional LUAD cells shifted toward the low-malignant AT1-like state, p53 signaling activity progressively increased, accompanied by gradual decreases in metabolic and proliferative activities. In contrast, transitions toward the high-malignant EMT-like state were characterized by enhanced metabolic and proliferative programs. In addition, the activity of the antigen processing and presentation pathway also showed a clear and continuous increase as transitional cells progressed toward the AT1-like state (**Supplementary Fig. 11e–f**).

To identify key signatures related to the malignant transition of the transitional state, we selected the top 200 genes most correlated with each fate potential and quantified each gene’s contribution to AT1- or EMT-biased fate potentials (**Supplementary Fig. 12, Fig. 5e**). The top 30 contributing genes for each fate branch were both significantly correlated with clinical prognosis in LUAD patients. Patients with higher AT1 fate potential showed relatively better survival outcomes, whereas those with higher EMT potential had poorer prognosis (**Fig. 5f-g, Supplementary Fig. 13**). Gene ontology (GO) enrichment analysis^56^ revealed that AT1 lineage-specific drivers were most strongly enriched in immune response-related biological processes (**Fig. 5h**). This suggests that during p53-driven transitions toward the AT1-like state, tumor immune evasion capacity is reduced. In contrast, EMT lineage-specific drivers were enriched in biological processes associated with cellular structural remodeling, metabolic regulation, and cell adhesion processes (**Fig. 5h**). Furthermore, analysis of large-scale LUAD patient RNA-seq cohorts revealed significant differential expression of lineage-specific genes between TP53 wild-type and mutant groups as well as between different survival outcome groups (**Supplementary Fig. 14a–d**). AT1 lineage-specific drivers were highly expressed in the TP53 wild-type group, supporting p53’s role as a key driver promoting transitions toward low-malignant AT1-like states. However, despite the significant associations, lineage-specific gene expression alone lacked predictive power for TP53 mutation status or survival outcomes (**Supplementary Fig. 14e–f**), likely due to additional layers of multi-omics regulation influencing tumor evolution beyond transcriptomic changes.

Collectively, DyMoTree effectively reconstructed biologically meaningful cell fate potentials within transitional LUAD cells, accurately identified fate-specific tumor cell states, and provided mechanistic insight into how p53 activity orchestrates tumor evolution toward tumor states with varying degrees of malignancy.

### DyMoTree dissects fate bias linked to therapeutic potency in pre-infusion CAR-T cells

To further demonstrate the versatility of DyMoTree in characterizing cell state transitions, we reanalyzed a longitudinal single-cell transcriptomic dataset of CD8^+^ CAR-T cells from 26 B-cell acute lymphoblastic leukemia (B-ALL) patients who received CAR-T infusion and exhibited heterogeneous clinical outcomes^57^. The dataset encompassed CD8^+^ CAR-T cells collected from the manufacturing stage (pre-infusion) through post-infusion time points up to 15 months, allowing us to trace the emergence of post-infusion states from their pre-infusion origins. In this setting, pre-infusion CD8^+^ CAR-T cells were treated as the progenitor population, while post-infusion cytotoxic and memory-like cell states were designated as terminal states (**Fig. 6a**).

**Fig. 6.**
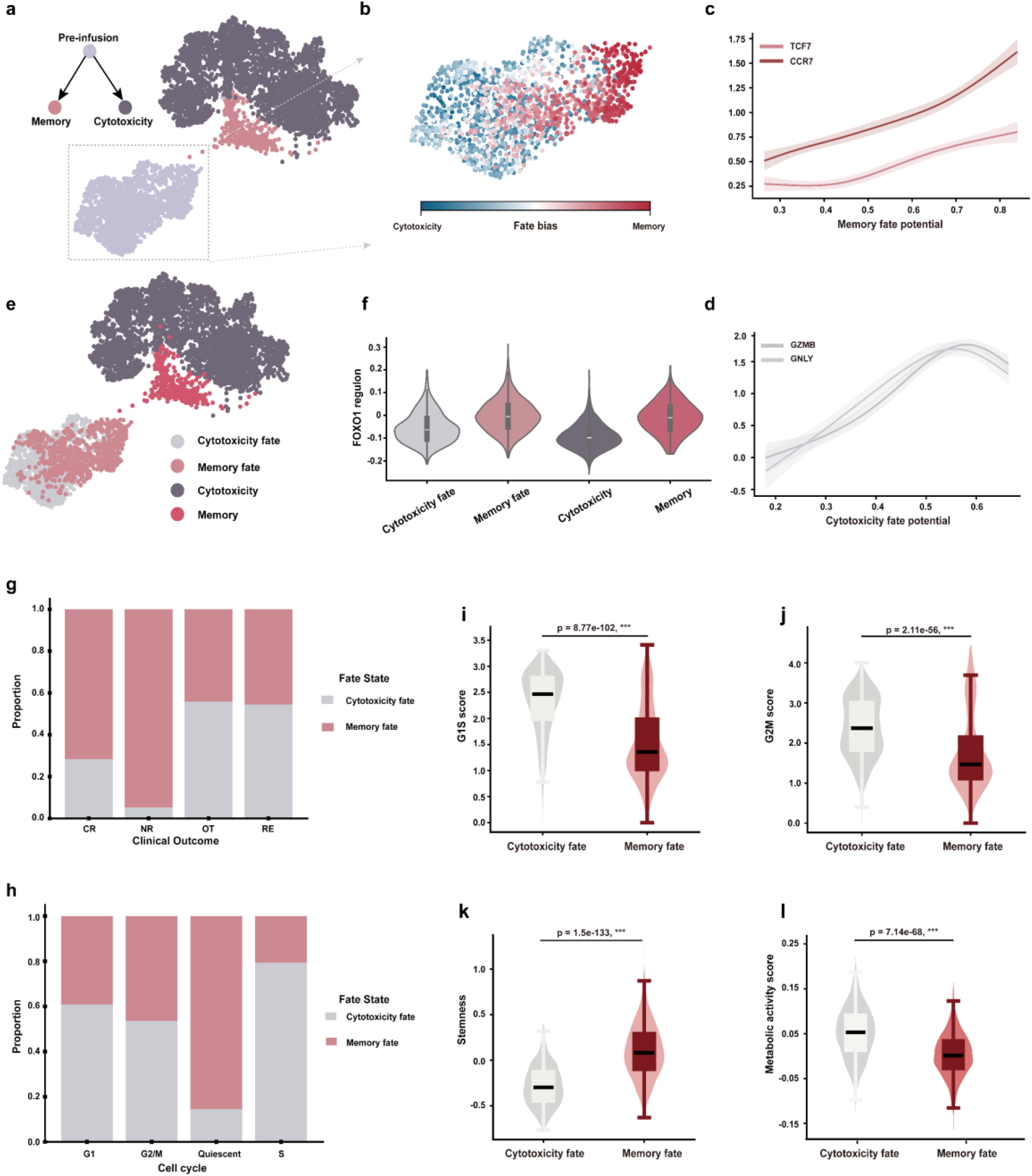
DyMoTree resolves pre-infusion CAR-T cell-state transitions linked to therapeutic potency and metabolic plasticity. **(a**) UMAP illustration of the CAR-T cell-specific differentiation process. Cells are annotated from the original study. **(b)** DyMoTree inferred cell fate bias toward memory-(red) and cytotoxicity-(blue) state in pre-infusion CAR-T cells. **(c-d)** Gene expression trends of marker genes corresponding to memory-state (top) and cytotoxicity-state (bottom) in pre-infusion CAR-T cells. Cells are sorted along cell fate potentials inferred by DyMoTree. **(e)** UMAP illustration of memory, cytotoxicity, and fate-specific states in pre-infusion CAR-T cells. **(f)** Violin plots show activity of FOXO1 regulon in memory, cytotoxicity, and fate-specific pre-infusion CAR-T states. **(g-h)** Fate-specific pre-infusion CAR-T-state composition in different clinical outcomes (top) and cell cycle stages (bottom). Annotations were all taken from the original study. **(i-l)** Violin plots show gene set scores of G1S, G2M, Stemness, and Metabolic in cytotoxicity- and memory-specific pre-infusion CAR-T cell states.

Although DyMoTree did not reveal a clearly demarcated fate boundary within pre-infusion CAR-T cells, it uncovered discernible fate directions toward cytotoxic and memory-like states (**Fig. 6b**). Consistent with the inferred fate potentials, T cell memory-state-related genes exhibited progressive upregulation along the memory-fate potential (**Fig. 6c**)^58, 59^, while genes related to cytotoxic function increased along the cytotoxic-fate potential (**Fig. 6d**)^60, 61^. Furthermore, by identifying genes that most strongly contributed to the cytotoxic-lineage fate potential, we recovered canonical cytotoxicity-related genes such as GZMB, GNLY, and NKG7, which are involved in cytotoxic granule exocytosis, a defining feature of highly active cytotoxic CAR-T cells^60, 61^ (**Supplementary Fig. 15a**). In contrast, genes most strongly correlated with the memory-lineage fate potential included CCR7, TCF7, and IL7R, well-established markers of memory T cells^58, 59^ (**Supplementary Fig. 15b**). In addition, STAT1, which is known to maintain naïve CD8 ^+^ T cell quiescence^62^, showed progressively increased expression with rising memory-fate potential scores (**Supplementary Fig. 15c–d**). Their upregulation suggests enhanced maintenance of the memory-like state in pre-infusion CAR-T cells. Taken together, these results demonstrate that we accurately inferred fate potentials corresponding to distinct post-infusion T cell states.

Based on fate potentials inferred by DyMoTree, we identified two fate-specific pre-infusion CAR-T states corresponding to memory-biased and cytotoxicity-biased lineages (**Fig. 6e**). The memory and cytotoxicity scores defined in the original study were significantly higher in the memory- and cytotoxicity-biased states, respectively (**Supplementary Fig. 16a–b**)^57^. Previous studies have established the transcription factor FOXO1 as a pivotal regulator that drives CD8^+^ T cell differentiation toward a memory state^63, 64^. In agreement with this, FOXO1 exhibited higher regulon activity in the memory-biased state compared to the cytotoxicity-biased state (**Fig. 6f and Supplementary Fig. 16c**). Building on these findings, we next characterized the phenotypic features of the two fate-specific pre-infusion CAR-T states. The previous study had also demonstrated that FOXO1 can enhance CAR-T persistence and therapeutic efficacy by driving the differentiation of CAR-T cells toward a memory state^64^. Interestingly, across patients with distinct therapeutic responses, memory-biased pre-infusion CAR-T cells were enriched in both complete responders (CR) and non-responders (NR) (**Fig. 6g**), underscoring the essential contribution of memory-like states to CAR-T cell persistence and antitumor activity. To further characterize the distinct biological properties of these two states, we conducted a comprehensive analysis of cellular activity and functional programs. The results showed that memory-biased cells were predominantly quiescent, residing in the G_0_ phase of the cell cycle, whereas cytotoxicity-biased cells displayed higher proliferative activity, as reflected by markedly elevated cell-cycle activity scores (**Fig. 6h–j**). In addition, the elevated expression of stemness-associated genes in the memory-biased state, together with the observed quiescent phenotype, indicates that this state possesses stem cell–like characteristics (**Fig. 6k**).

We further applied scMetabolism to quantify pathway activities in the two fate-specific pre-infusion CAR-T cell states and characterize their metabolic programs (**Fig. 6l, Supplementary Fig. 17**)^65^. In the cytotoxicity-biased state, cells showed pronounced activation of energy-producing and biosynthetic pathways, including the TCA cycle, fatty acid elongation, purine, folate, and one-carbon metabolism, as well as multiple amino acid pathways. This pattern reflects a highly metabolic effector phenotype characterized by enhanced mitochondrial respiration and nucleotide biosynthesis. In contrast, the memory-biased state exhibited selective upregulation of lipid- and glycan-related processes, including sphingolipid metabolism, glycosphingolipid biosynthesis, O-glycan and glycosaminoglycan biosynthesis. These changes suggest that metabolic reprogramming occurs toward membrane and signaling remodeling in the memory-biased state. Overall, we speculate that cytotoxicity-biased cells exhibit a relatively high level of energy metabolism to sustain effector functions, whereas memory-biased cells predominantly upregulate lipid- and glycan-related pathways, suggesting that membrane remodeling may play an important role in maintaining stem-like potential within the memory state.

Taken together, all results suggest that DyMoTree can effectively resolve early cell-state transition dynamics and uncover biologically meaningful mechanisms across multiple biological systems.

## Discussion

In this study, we developed DyMoTree, a computational framework for modeling lineage-resolved cell state transitions from single-cell transcriptomic data. Unlike existing approaches that infer cell fate through trajectory reconstruction, Markov diffusion, or optimal transport^10, 12-16, 18-21^, this framework reframes fate inference as the learning of nonlinear transition mappings between progenitor and terminal cell states within a structured lineage hierarchy. By integrating similarity-derived lineage graphs with lineage-structured deep neural networks, DyMoTree enables robust inference of early progenitor fate potentials and facilitates the identification of fate-specific progenitor substates and lineage-associated regulatory programs.

DyMoTree advances lineage-resolved fate inference through three main methodological components. First, the framework formulates cell fate inference as the learning of nonlinear mappings between progenitor and terminal cell states under explicit lineage constraints. By integrating cell-state similarity with lineage hierarchy information, it constructs lineage graphs that capture both local relationships among cells and cross-lineage differentiation patterns, providing a structured representation for modeling lineage-level transitions. Second, these transitions are learned through a lineage-structured deep neural network that combines graph neural networks with cross-attention mechanisms^30, 66^. The graph neural network modules capture local transition patterns within cell states, whereas the cross-attention mechanism models the contributions of progenitor cells to terminal states and produces an interpretable transition map quantifying lineage-level coupling. Third, the inferred transition map enables characterization of the progenitor fate landscape and identification of fate-specific progenitor substates. By integrating fate potentials with sparse modeling and network-based smoothing, the framework further identifies lineage-specific driver genes and regulatory programs underlying cell fate commitment.

Existing trajectory inference frameworks typically model cellular dynamics as continuous flows on a single connected manifold, where transitions occur through local neighborhoods in gene expression space. This formulation is well-suited for modeling within-lineage differentiation processes characterized by gradual transcriptional changes. However, these approaches generally rely on assumptions of local continuity and first-order Markovian dynamics. As a result, they may be less suited for modeling cross-lineage or cross-hierarchical transitions, where cell states may reside in stratified manifold components, and transitions are not necessarily supported by locally continuous paths. In contrast, DyMoTree models differentiation as lineage-level mappings between progenitor and terminal states under explicit lineage constraints, enabling the inference of early fate biases even when transcriptional divergence remains subtle.

Benchmark analyses showed that the proposed framework consistently outperformed existing methods in inferring early cell fate biases across diverse datasets. It accurately recovered progenitor fate potentials in both simulated systems and lineage-tracing datasets and remained robust in complex *in vivo* settings such as CD8^+^ T-cell differentiation during viral infection. The method also exhibited stable performance across a wide range of neighborhood parameters, highlighting its robustness to variations in manifold topology. Beyond predictive performance, DyMoTree provided biologically meaningful insights across multiple biological contexts. In developmental systems, its application to early mouse embryogenesis reconstructed the second cell-fate decision within the inner cell mass, recapitulating the opposing expression dynamics of *Nanog* and *Gata6*. In disease settings, the framework resolved lineage specification of transitional lung adenocarcinoma cells and revealed distinct fate programs associated with tumor progression. Furthermore, in CAR-T immunotherapy for B-ALL, DyMoTree identified pre-infusion T-cell subpopulations biased toward memory-like or cytotoxic states, linking early transcriptional programs to post-infusion persistence and therapeutic efficacy.

Because DyMoTree operates on graphs constructed from low-dimensional embeddings, the framework is inherently modality-agnostic and can, in principle, be extended to other single-cell omics modalities that support neighborhood graph construction. In this study, we focused on scRNA-seq data using PCA-based embeddings, but the framework remains flexible with respect to feature representation and can incorporate alternative embedding strategies suited to different data modalities. Despite its robust performance, several challenges remain. The topology of lineage graphs may vary substantially across biological systems, and differences in graph density or transition patterns can influence model optimization and require moderate parameter tuning. In addition, the lineage graph design increases computational complexity for large-scale datasets, as both intra-state and inter-state graphs must be constructed and reconstructed during training. Modeling highly complex lineage architectures, such as deeply branched or cyclic differentiation processes, also remains challenging. Future work may address these limitations by integrating the framework with multimodal and spatial datasets and by developing more scalable strategies for modeling large and complex lineage structures.

Overall, DyMoTree provides a versatile framework for modeling lineage-resolved cell fate transitions directly from single-cell transcriptomic data. By enabling robust inference of early fate biases and identification of lineage-associated regulatory programs, it offers a useful computational approach for studying cellular differentiation across diverse biological contexts, including development, regeneration, cellular reprogramming, and disease progression.

## Methods

### Description of lineage differentiation structure

In DyMoTree, we require a lineage differentiation structure that specifies the hierarchical relationships between cell states. This hierarchy is represented as a directed tree ***L*** = (***S, E***), where ***S*** denotes the set of cell states, and ***E*** denotes the set of directed edges representing differentiation relationships between cell states. Each edge ***e*** = (***p, t***) ∈ ***E*** indicates a differentiation direction from a progenitor state ***p*** to a terminal state ***t***. For each lineage pair (***p, t***), we define the corresponding cell sets ***C***_***p***_ and ***C***_***t***_, which contain all single cells annotated as belonging to states ***p*** and ***t***, respectively. DyMoTree models cell-state transitions from the progenitor cell population ***C***_***p***_to the terminal cell population ***C***_***t***_ at single-cell resolution.

### Cell fate inference for early progenitor state

In DyMoTree, cell fate inference is implemented within a graph-centric integrative framework that combines lineage graph construction with a tree-structured neural network. The lineage graph consists of an intra-state transition graph and an inter-state transition graph. The inter-state transition relationships within the lineage graph are constructed based on a composite similarity metric at single-cell resolution. The tree-structured neural network performs cell fate inference by cross-attention-based feature mapping constrained by the lineage graph. In addition to learning informative cell embeddings, which is a common objective in neural network-based models, DyMoTree places particular emphasis on the interpretability of the cross-attention mechanism and uses it to model cross-state transition patterns from progenitor to terminal cells on the lineage structure.

#### Composite similarity metric

The similarity metric in DyMoTree is defined by combining the shortest-path distance with a linear kernel term, enabling comprehensive characterization of lineage-aware cell–cell similarity under a given lineage structure.

#### Shortest-path manifold distance

Given a lineage structure *L* = (*S, E*), let:

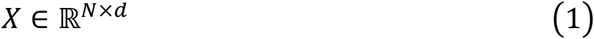

denote the embedding matrix of all cells in *L*, where N is the number of cells in *S* and d is the dimension of cell embedding. By default, we selected the top 20 principal components (PCs). Each row of X corresponds to a cell, and each column corresponds to a principal component.

A K-nearest-neighbor (KNN) graph *G* is constructed in the PCA space using Euclidean distance. Each cell is connected to its K nearest neighbors, with K = 50 by default **(Supplementary Fig. 18)**. To facilitate shortest-path computation, the adjacency matrix of *G* is symmetrized, yielding an undirected graph. The shortest-path distance matrix from progenitor cells *C*_*p*_ to all terminal cells *C*_*t*_ is denoted as:

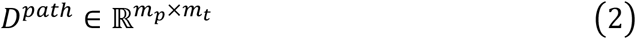

Where *m*_*p*_ and *m*_*t*_ are the numbers of progenitor and terminal cells, respectively.

#### Linear kernel similarity

To capture transcriptional alignment between a progenitor cell *i* ∈ *C*_*p*_ and a terminal cell *j* ∈ *C*_*t*_, we compute a linear kernel between their PCA embeddings:

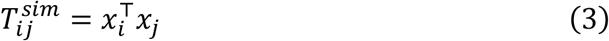

This term complements the manifold-based distance by emphasizing global linear similarity in the feature space.

To integrate both metrics, the shortest-path distances are first converted into a transition preference matrix:

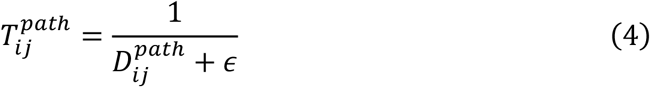

Where *ϵ* > 0 ensures numerical stability. Both *T*^*path*^ and *T*^*sim*^ are then column-wise normalized using min-max scaling for each terminal cell *j*:

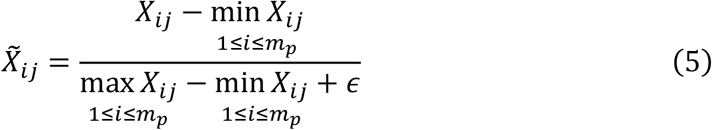

Finally, the overall composite similarity matrix is obtained by summing the normalized matrices:

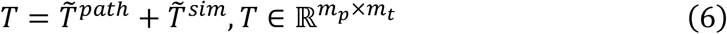

This composite metric effectively integrates local manifold topology and global transcriptional similarity, providing a robust measure for lineage-level cell-cell relationships.

#### Intra-state Transition Graph

To characterize cell-state-specific transition patterns, we construct a K-nearest neighbor (KNN) graph 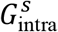 for each cell state *s* in the PCA-reduced gene expression space. Each node represents a single cell, and each edge encodes the transcriptional similarity-based potential transition from a cell to its neighboring cells. The number of neighbors *k* is chosen consistently with that used in the shortest-path distance computation for the composite similarity metric. Through empirical tuning, we set *k* = 50 by default **(Supplementary Fig. 18)**. This intra-state graph captures the local topology and directional relationships among cells within the same state, serving as a foundation for the subsequent graph-based neural network modeling.

#### Inter-state Transition Graph

Lineage-level state transitions are modeled as processes from progenitor states *s*_*p*_ to terminal states *s*_*t*_. For each lineage pair (*s*_*p*_, *s*_*t*_) in the lineage structure, we construct a bipartite graph 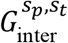, where the two node sets correspond to the progenitor cell population *C*_*p*_ and the terminal cell population *C*_*t*_, respectively. Each edge represents a potential transition from a progenitor cell to a terminal cell.

The bipartite edges are derived from the composite similarity matrix *T*, which quantifies the transition preference between progenitor and terminal cells. To capture the relative propensity of a progenitor cell *i* ∈ *C*_*p*_ toward a specific terminal cell *j* ∈ *C*_*t*_, the similarity matrix is first normalized along the progenitor dimension:

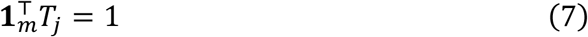

Where *m* = ∣ *C*_*p*_ ∣ is the number of progenitor cells, and *T*_*j*_ is the column corresponding to terminal cell *j*.

For a given terminal state *s*_*t*_, we extract the subset of the similarity matrix corresponding to all terminal cells in *C*_*t*_:

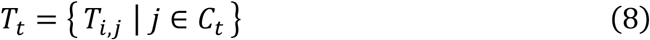

We then compute the expected transition propensity of progenitor cell *i* toward the terminal state *s*_*t*_ as the mean over all terminal cells in that state:

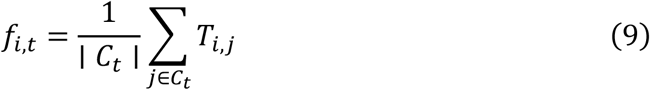

A bipartite edge between progenitor cell *i* and terminal cell *j* is established only if the cell-level transition propensity exceeds the expected propensity toward other terminal states:

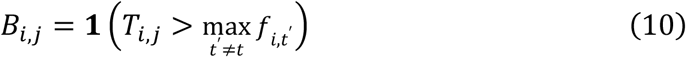

Where **1**(⋅) is the indicator function. Given the noise inherent in transcriptome-derived similarity, negative edges were retained only for progenitor cells showing a high degree of dissimilarity from the corresponding terminal state, defined as having negative connections to more than 80% of terminal cells by default. This ensures that edges capture lineage-specific differentiation preferences while filtering out weaker or non-specific transitions.

#### Lineage Graph

The complete lineage graph is obtained by integrating the intra-state and inter-state graphs. Formally, the lineage graph *G*_lineage_ consists of the union of all intra-state graphs 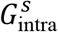 across cell states and all inter-state bipartite graphs 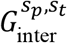 across lineage pairs. This hierarchical graph encodes both local cell–cell relationships within states and global progenitor-to-terminal transitions across states, providing a structured scaffold for the tree-structured neural network to learn nonlinear lineage-level mappings and perform early cell fate inference.

#### The tree-structured neural network

The neural network model in the DyMoTree framework adopts an encoder–decoder architecture. The encoder is organized according to the lineage differentiation structure and consists of CellModules and LineageModules. CellModules correspond to individual cell states and learn transition-related cell embeddings. LineageModules correspond to differentiation relationships between cell states and learn nonlinear transition patterns from progenitor states to terminal states. DyMoTree adopts a dual decoder, which consists of an intra-state decoder and an inter-state decoder. The intra-state component reconstructs intra-state transition graphs, whereas the inter-state component reconstructs inter-state transition graphs.

#### CellModule

Each cell state in the lineage structure is associated with a parameter-independent *CellModule*, which models the intra-state transition topology among cells. We denote the intra-state transition graph as

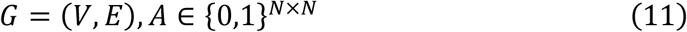

Where *A*_*ij*_ = 1 indicates a potential transition from cell *𝓋*_*i*_ (ancestor) to cell *𝓋*_*j*_ (descendant), and *A*_*ij*_ = 0 otherwise. The node features are given by the gene expression matrix

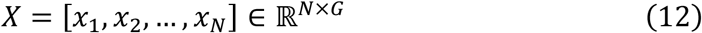

Where *x*_*i*_ ∈ ℝ^*G*^ denotes the expression profile of the cell *𝓋*_*i*_, and *G* is the number of genes.

To mitigate technical noise and sparsity inherent in single-cell RNA sequencing data, we first project the raw gene expression profiles into a latent feature space using a two-layer multilayer perceptron (MLP):

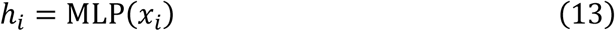

Where *h*_*i*_ ∈ ℝ^*F*^ denotes the latent representation of cell *𝓋*_*i*_, and *F* is the dimensionality of the latent feature space.

#### Ancestor-aggregated message passing

Unlike conventional graph attention networks that aggregate messages from neighboring nodes without explicit biological directionality, we propose an ancestor-aggregated message passing scheme to explicitly model information propagation along developmental transitions. Specifically, for each target (descendant) cell, the model aggregates information from all its candidate ancestor cells, mimicking a one-step transition process within the cell state manifold. This formulation allows the learned representations to reflect local developmental trajectories rather than undirected neighborhood similarity.

Given a descendant cell *𝓋*_*j*_, we estimate the importance of each candidate ancestor cell *𝓋*_*i*_ ∈ 𝒩_*j*_ using an attention mechanism based on their latent representations. To explicitly encode transition directionality, we introduce independent linear transformations for ancestor and descendant cells:

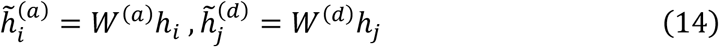

where *W*^(*a*)^ and *W*^(*d*)^ are learnable weight matrices.

The transition score between cells *𝓋*_*i*_ (ancestor) and *𝓋*_*j*_ (descendant) is defined using a cosine attention function:

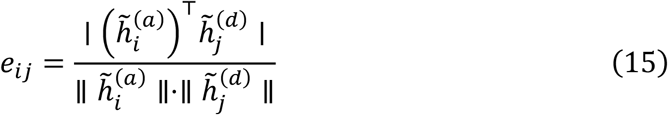

Where ∥⋅∥ denotes the Euclidean norm. The absolute value captures the strength of transition similarity while ignoring sign-specific effects.

The attention coefficients are normalized across all candidate ancestor cells using a temperature-scaled softmax function:

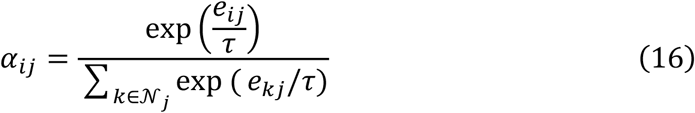

Where 𝒩_*j*_ denotes the set of candidate ancestor cells for *𝓋*_*j*_, and τ is a temperature parameter controlling the sharpness of the distribution. A smaller τ leads to a more concentrated distribution, enabling the model to focus on a small number of high-confidence transitions.

The embedding of each descendant cell is updated by aggregating messages from its ancestor cells according to the learned attention coefficients. We employ a multi-head attention mechanism to capture diverse transition patterns:

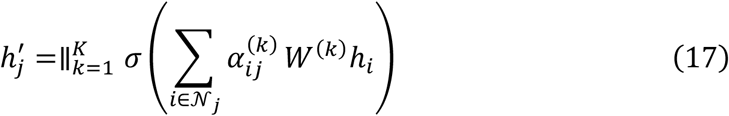

Where *K* denotes the number of attention heads, *W*^(*k*)^ is the learnable transformation matrix for the *k*-th head, *σ* is a nonlinear activation function, and ∥ denotes concatenation.

To capture higher-order transition structures within the cell state manifold, we stack two graph attention layers:

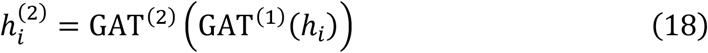

Where 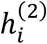 represents the final embedding of the cell *𝓋*_*i*_, encoding local intra-state transition topology.

#### LineageModule

Each LineageModule is designed to model cell state transitions within a lineage pair

(𝒫, 𝒯), where 𝒫 denotes a progenitor state, and 𝒯 denotes a terminal state. The module is parameter-independent, with a distinct module instantiated for each lineage pair. Given a lineage pair, the latent embeddings obtained from the corresponding CellModules are denoted as

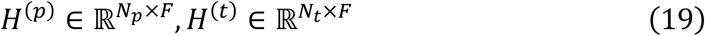

Where *N*_*p*_ and *N*_*t*_ denote the numbers of progenitor and terminal cells, respectively, and *F* is the embedding dimensionality.

To model directional information flow from progenitor to terminal states, the LineageModule explicitly models cross-state relationships by learning a nonlinear mapping from progenitor cells to terminal cells. This is achieved through a cross-attention mechanism, in which terminal cells serve as queries and progenitor cells serve as keys, to selectively aggregate information from progenitor cells, thereby capturing lineage-dependent feature transformations across different stages of cellular differentiation.

#### Cross-attention formulation

The embeddings are first projected into query and key spaces via learnable transformations:

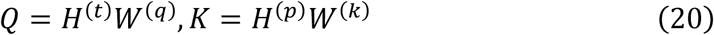

Where *W*^(*q*)^ and *W*^(*k*)^ are learnable weight matrices. The unnormalized cross-attention matrix is computed as:

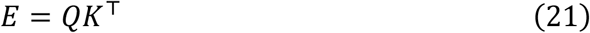

We first apply linear normalization to the attention scores, followed by a sigmoid transformation that maps the scores to bounded, non-negative values while keeping them in the near-linear regime of the sigmoid function to improve training stability:

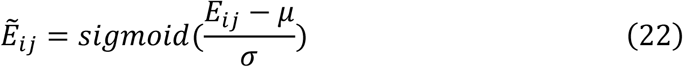

Where *μ* and *σ* denote the global mean and standard deviation of *E*, respectively. The sigmoid-transformed normalized attention scores are then used as the cell-state transition matrix, with each element representing the inferred transition strength from a progenitor cell to a terminal cell.

The attention weights are then defined as:

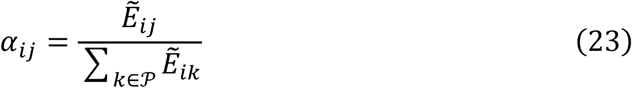

such that 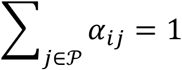 for each terminal cell 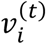.

#### Cross-state feature aggregation

The predicted embeddings of terminal cells are obtained by aggregating progenitor embeddings using the learned attention weights:

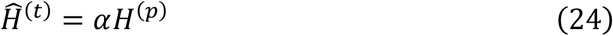

Where 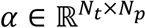 is the attention matrix.

The decoders are designed to match the distinct topological properties of the two reconstruction tasks: a directed within-state transition graph and a bipartite cross-state transition graph.

#### Intra-state decoder

For each cell state, we employ an intra-state decoder to reconstruct the directed intra-state transition graph from the latent cell embeddings learned by the corresponding *CellModule*. To capture asymmetric transition likelihoods between cells, we adopt a gravity-inspired decoding mechanism, which models the probability of a directed edge as a function of both relative position and node-specific importance in the latent space.

Specifically, for each cell embedding *h*_*i*_ ∈ ℝ ^*F*^, we partition it into a positional component and a mass component:

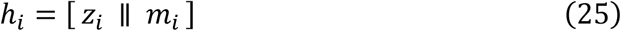

Where *z*_*i*_ ∈ ℝ ^*F*−1^ denotes the positional embedding of cell *𝓋*_*i*_, and *m*_*i*_ ∈ ℝ denotes its mass term. The directed edge probability from cell *𝓋*_*i*_ to cell *𝓋*_*j*_ is defined as

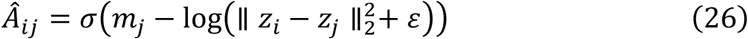

Where *σ*(⋅) denotes the sigmoid function, and *ε* is a small constant for numerical stability.

This decoder encourages cells with nearby positional embeddings and larger mass values to receive incoming transitions with higher probability, thereby providing a geometry-aware reconstruction of the local intra-state transition topology.

#### Inter-state decoder

For each lineage pair (𝒫, 𝒯), we employ an inter-state decoder to reconstruct the bipartite transition graph from progenitor cells in state 𝒫 to terminal cells in state 𝒯. Unlike the intra-state decoder, which focuses on local geometric organization within a cell state, the inter-state decoder is designed to evaluate whether the learned cross-state representations preserve transition compatibility across different developmental stages.

Given the progenitor embeddings 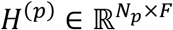 and the terminal embeddings 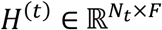, the edge probability from the progenitor cell 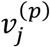 to the terminal cell 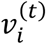 is modeled as

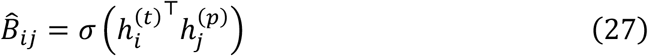

Where 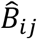 denotes the predicted probability of an inter-state edge, and *σ*(⋅) is the sigmoid function.

This formulation decodes cross-state transitions through latent feature compatibility and serves as a reconstruction objective for the nonlinear progenitor-to-terminal mapping learned by the *LineageModule*.

#### Training strategy

To enable DyMoTree to capture both local intra-state transition topology and cross-state lineage-dependent transformations, we adopt a staged training strategy consisting of two pretraining phases followed by a formal training phase. This progressive optimization scheme stabilizes representation learning by first preserving within-state structure and then introducing cross-state supervision.

##### Stage 1: Intra-state pretraining

In the first stage, each *CellModule* is trained independently on its corresponding intra-state transition graph. The objective of this stage is to learn cell embeddings that preserve the local transition topology within each cell state. For a given cell state, the intra-state reconstruction loss is defined as

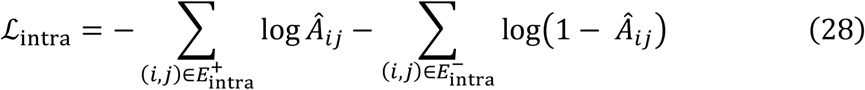

Where 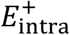 and 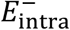 denote the sets of positive and negative edges in the intra-state graph, respectively, and *Â*_*ij*_ is the predicted edge probability.

This stage initializes each *CellModule* with state-specific representations that reflect local developmental geometry before introducing lineage-level interactions.

##### Stage 2: Coupled inter-lineage pretraining

In the second stage, DyMoTree jointly optimizes the *CellModules* associated with each lineage pair (𝒫, 𝒯). For each pair, the model simultaneously reconstructs the intra-state graphs of the progenitor and terminal states, as well as the inter-state bipartite graph connecting them. The inter-state reconstruction loss is defined as

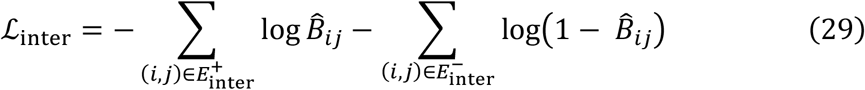

Where 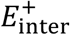 and 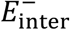 denote the positive and negative edges in the inter-state bipartite graph, respectively.

The total loss in this stage is given by

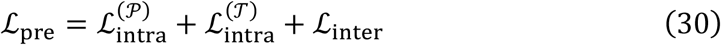

By jointly optimizing intra-and inter-state reconstruction objectives, this stage preserves local transition structure while introducing lineage-aware cross-state supervision.

##### Formal training

After pretraining, DyMoTree is trained on the full lineage tree using the same reconstruction objectives as in Stage 2. During this phase, for each non-root cell state, the embeddings predicted by the LineageModule conditioned on its progenitor state are used together with the corresponding CellModule to reconstruct both the intra-state graph and the inter-state bipartite graph. In this way, the model propagates information progressively along the lineage tree and learns biologically meaningful cross-state transition patterns.

To stabilize optimization, we employ a differential learning rate strategy, assigning a smaller learning rate to the CellModules to preserve the previously learned local transition representations, and a larger learning rate to the *LineageModules* to emphasize the learning of cross-state nonlinear mappings. In addition, all experiments are repeated with 20 independent random seeds, and the final results are reported as averages to mitigate stochastic variation. The hyperparameters used for each dataset are summarized in **Supplementary Tables 2–4**.

#### Edge sampling strategy

To improve computational efficiency and maintain a balanced reconstruction objective, we perform edge sampling separately for intra-state and inter-state graphs during training.

For the intra-state graph, all positive edges are retained. At each iteration, we randomly sample an equal number of negative edges from unobserved cell pairs within the same state, thereby constructing a balanced set of positive and negative examples for intra-state reconstruction.

For the inter-state bipartite graph, mini-batch sampling was performed in a progenitor-cell-centered manner. At each iteration, progenitor cells were independently sampled from the positive and negative edge sets, and all inter-state edges associated with the sampled cells were collected as reconstruction edges for the current optimization step.

The same progenitor-cell batch size was used for both positive and negative edge sets. This strategy preserves coherent progenitor-centered transition neighborhoods within each batch while improving scalability for large bipartite graphs.

#### Defining progenitor cell fate potential toward terminal states

Beyond the latent embeddings learned by the CellModules, the core output of DyMoTree is the sigmoid-transformed normalized attention score matrix 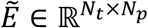 produced by each LineageModule, which encodes the lineage-dependent mapping from progenitor cells to terminal cells for a given lineage pair (𝒫, 𝒯). Each element *e*_*ij*_ represents the contribution of the progenitor cell 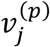 to the terminal cell 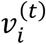, as defined in the LineageModules. For convenience, we define the *state transition matrix* as:

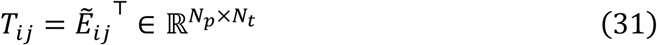

where 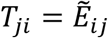 represents the contribution from the progenitor cell 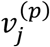 to the terminal cell 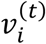.

The fate potential of a progenitor cell 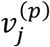 toward a terminal state 𝒯 is defined as the average contribution across all terminal cells in that state:

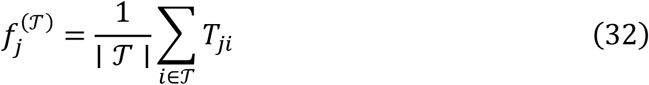

Where 𝒯 denotes the set of terminal cells belonging to the target state.

For a progenitor state with *K* terminal states, we compute *K* fate potential vectors:

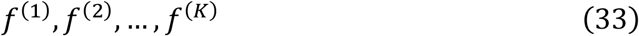

By concatenating these vectors, we obtain the cell fate matrix:

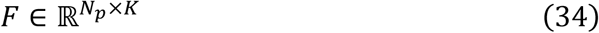

Where *F*_*jk*_ represents the fate potential of the progenitor cell 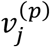 toward the *k*-th terminal state.

The matrix *F* provides a quantitative characterization of lineage bias at the single-cell level. Each row describes the vector of differentiation potential of a progenitor cell across multiple terminal states, enabling downstream analyses such as identification of fate-biased progenitor subpopulations and lineage-specific driver genes.

#### Identification of Fate-specific Progenitor Substates

To identify fate-specific substates within a progenitor cell population, we integrated the fate potential matrix derived from DyMoTree with Archetypal Analysis (AA) **(Extended Data Fig. 5)**. AA identifies a small set of archetypes, extreme representative points that collectively approximate the convex hull of the underlying data manifold. Within the fate manifold, each archetype corresponds to an extremal yet characteristic fate state, providing interpretable vertices that define principal differentiation directions.

To improve the representation of the fate manifold in gene expression space, we first performed feature selection by retaining genes whose expression levels were significantly correlated with the inferred fate potential. This step enriches for genes directly associated with differentiation bias and reduces background transcriptional noise. The expression profiles of the selected genes were then projected into a fate-embedding space using Principal Component Analysis (PCA), capturing the dominant sources of fate-associated variance while mitigating technical noise.

Next, to model the intrinsic topology of progenitor cell states, we constructed an undirected K-nearest neighbor (KNN) graph based on Euclidean distances in the fate-embedding space. This graph preserves local similarity among cells in terms of fate potential and serves as an approximation of the continuous fate manifold. We then applied the Diffusion Map algorithm to learn a smooth, topologically consistent representation of the progenitor cell fate manifold. The resulting diffusion feature matrix is denoted as *X*_*d*_, where *d* is the dimensionality of the diffusion features. In practice, we set both the number of PCA components and diffusion features to 10.

Finally, we applied Archetypal Analysis to the diffusion-based feature matrix to approximate the fate manifold with a limited number of archetypes representing principal fate directions. Formally, AA decomposes the data matrix as:

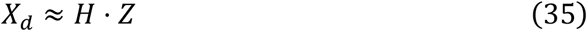

Where *Z* is the archetype matrix, *H* is the assignment matrix, and *s* denotes the number of archetypes. Each archetype in *Z* corresponds to a representative position within the diffusion-based fate manifold, and each cell is expressed as a convex combination of these archetypes, as specified by its row in *H* . The probabilistic assignment encoded in *H* was then used to group cells into fate-related progenitor substates. Cells with similar assignment profiles were clustered together, delineating distinct progenitor substates that reflect the intrinsic organization of the fate manifold.

#### Identification of Lineage-specific Genes

To identify lineage-specific driver genes, DyMoTree integrates fate potentials with sparse modeling and network-based refinement **(Extended Data Fig. 6)**. Since cell fate decisions are often governed by a limited set of key genes, we employed a two-step strategy.

##### Step 1: Sparse Regression

Candidate genes were first ranked based on their correlation with fate potential scores, prioritizing those most likely contributing to lineage-specific transitions. Next, Lasso regression was applied to select genes strongly associated with fate potential:

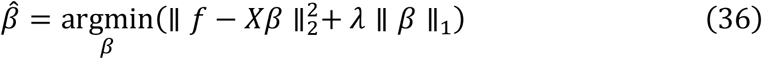

Where *f* denotes the fate potential scores for a specific lineage, *X* is the expression matrix of candidate genes, *β* represents gene-specific contribution weights, and *λ* is the sparsity regularization parameter. This step enforces sparsity, retaining only a small subset of genes with the strongest contributions.

##### Step 2: Network-based Smoothing

To improve robustness against technical noise and sparsity inherent to single-cell data, Lasso-derived coefficients were refined via graph convolutional smoothing over a gene co-expression network constructed among selected genes. Edges were defined based on pairwise gene–gene correlations, with a minimum correlation threshold to remove spurious links. The contribution scores were diffused across the network:

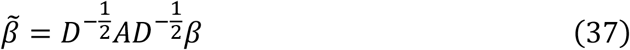

Where *A* is the adjacency matrix of the co-expression network, and *D* is the degree matrix. This process propagates contribution signals across correlated genes, yielding more robust estimates of lineage-specific influence.

Finally, genes were ranked by their smoothed contribution scores 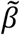, and the top-ranked genes were designated as lineage-specific drivers.

### Performance Evaluation and Benchmark Metrics

To rigorously assess the performance of DyMoTree, fate bias scores were computed for single-cell lineage-tracing and CD8^+^ T-cell differentiation datasets using ground-truth fate potentials:

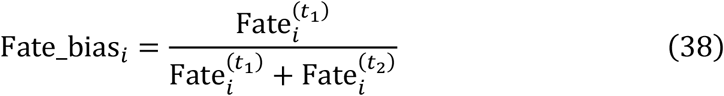

Where Fate_bias_*i*_ ∈ [0,1] quantifies the propensity of progenitor cell *i* to differentiate toward two terminal states. A score greater than 0.5 indicates a bias toward terminal state 1, while a score below 0.5 indicates a bias toward terminal state 2.

To benchmark DyMoTree against state-of-the-art (SOTA) methods, we evaluated both classification performance and biological interpretability using five metrics.

#### 1. Classification Metrics

To measure the accuracy of fate bias predictions, we calculated AUROC, Accuracy, and F1-score:

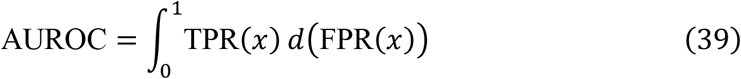

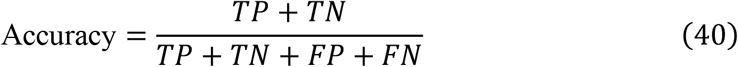

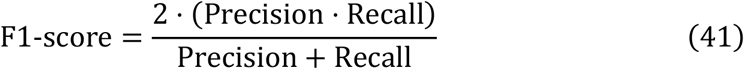

With Precision and Recall defined as:

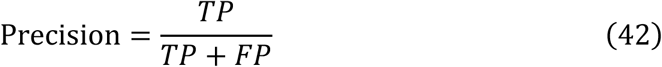

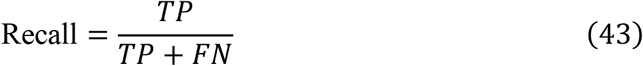

Here, *TP, TN, FP*, and *FN* denote true positives, true negatives, false positives, and false negatives, respectively.

#### 2. Correlation Metrics

To assess the biological relevance of predicted fate biases, we calculated the Pearson correlation coefficient and Spearman’s rank correlation coefficient:

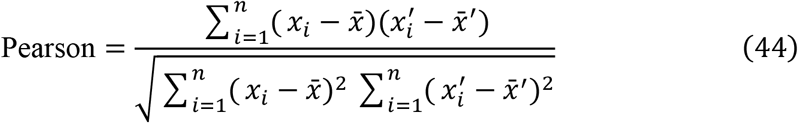

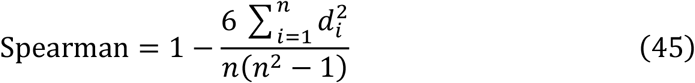

Where *x*_*i*_ and 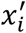 represent the predicted and ground-truth fate bias for progenitor cell *i*, respectively; 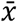 and 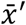 denote their mean values; and *d*_*i*_ is the rank difference between *x*_*i*_ and 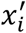. These metrics collectively evaluate both the predictive accuracy and biological interpretability of DyMoTree in inferring early cell fate biases.

#### GO enrichment analysis

Gene Ontology (GO) enrichment analysis was performed using the clusterProfiler package, restricting GO terms to the Biological Process (BP) category. Only enriched terms containing more than five genes from the input gene set were retained for evaluation and downstream analysis. GO terms with false discovery rates (FDR) less than 0.05, calculated using the Benjamini–Hochberg correction, were considered statistically significant.

#### Reproducibility and statistical framework

All analyses were performed using Python (version 3.10) and R (version 4.3) with standard bioinformatics packages, including Scanpy, Seurat, and scikit-learn. In all correlation-based analyses, Pearson correlation coefficients were used unless otherwise specified. For the identification of fate-specific progenitor states, Spearman correlation was employed to capture monotonic relationships between gene expression and inferred fate potential, allowing compact representation of the fate manifold in expression space.

### Datasets and pre-processing

#### Simulated data generation

Synthetic scRNA-seq datasets were generated using the ‘splatSimulatePaths’ function in the Splatter package to evaluate the performance of DyMoTree. The simulation models cell state transitions from a stem state to different child states. Key parameters were adjusted to shape the desired dynamic features: ‘group.prob’ controlled the relative sizes of each cell state, ‘path.from’ specified the lineage connections, ‘de.prob’ and ‘de.facScale’ defined the proportion and magnitude of differentially expressed genes, ‘path.skew’ modulated the distribution bias of cells toward the source or target states, and ‘path.nSteps’ determined the smoothness of state transitions.

#### Single-cell lineage-tracing datasets

All lineage-barcoded cells from the original dataset were retained for analysis. Cells belonging to the monocyte and neutrophil lineages were selected at days 2, 4, and 6 based on lineage barcodes. Following the computational framework established by Weinreb et al., ground-truth fate potentials for hematopoietic stem and progenitor cells (HSPCs) were calculated as the proportion of each clone’s descendant cells committing to the monocyte or neutrophil lineage. We used only day 2 HSPCs carrying ground-truth fate biases for quantitative comparison. In addition, we uniformly sampled comparable numbers of monocyte and neutrophil cells to correct for density imbalance between progenitor and terminal populations, enabling a fair comparison across different computational methods. We also reported the comparison results across terminal cells at different time points, which showed that DyMoTree is robust for cell number (Supplementary Fig. 19).

#### CD8^+^ T cell differentiation datasets

We followed the original study’s annotation, including early effectors (EE), intermediate effectors (IE), terminal effectors (TE), and memory precursors (MP). EE and IE cells were merged to represent the progenitor population giving rise to TE and MP states. Cells with detected T cell receptor (TCR) sequences from single-cell TCR sequencing (scTCR-seq) were retained, and those containing MP-biased or TE-biased TCR clonotypes defined in the original publication were selected. Ground-truth fate potentials for progenitor T cells were then computed, analogous to the lineage-tracing dataset, as the proportion of descendant cells within each clonotype committing to the TE or MP lineage.

#### Mouse early embryogenesis datasets

From the original dataset, all annotated inner cell mass (ICM), primitive endoderm (PrE), and epiblast (Epi) cells collected from mouse embryos at embryonic days E3.5 to E4.5 were extracted for analysis.

#### Lung adenocarcinoma evolution datasets

scRNA-seq data from KT and KPT mouse lung adenocarcinoma (LUAD) models were obtained for analysis. Following the original study’s clustering scheme, cluster 5 (Transitional) cells were extracted and defined as the progenitor population, while cluster 3 (AT1-like) and cluster 11 (epithelial–mesenchymal transition-like) cells were selected as terminal states representing alternative differentiation outcomes of LUAD tumor cells.

#### B-cell acute lymphoblastic leukemia CAR-T cell differentiation datasets

Based on the annotations provided in the original dataset, all pre-infusion CAR-T cells (pre-IP) were defined as the progenitor state. Among post-infusion cells, the three cytotoxic clusters (Cytot-GZMB, Cytot-GZMH, and Cytot-GZMK) were combined into a single cytotoxic (Cyto) state. The Cyto state and memory-like (Mem) states were used as terminal states for modeling CAR-T cell differentiation.

### Data pre-processing

Raw expression matrices were first normalized and log-transformed, followed by the selection of the top 2,000 highly variable genes for downstream analysis. Principal component analysis (PCA) was then applied for dimensionality reduction. For visualization, lineage-tracing datasets were displayed in the two force-directed layouts (SPRING) provided by the original study, the LUAD dataset was visualized using Diffusion embedding, and all other datasets were visualized using Uniform Manifold Approximation and Projection (UMAP).

## Data availability

All the datasets analyzed in this study are publicly available. The single-cell lineage-tracing datasets are available in the ‘single cell lineage tracing database’ (scLTdb, https://scltdb.com/scLT/). The scRNA-seq datasets are available in the Gene Expression Omnibus (GEO) under accession codes GSE241403 for CD8^+^ T cell datasets, GSE152607 for LUAD tumor cells, and GSE162975 for CAR-T cells. The mouse gut endoderm development datasets can be explored and are available at https://endoderm-explorer.com/.

## Code availability

The DyMoTree framework is implemented in Python. The source code of DyMoTree and the codes for reproducing the results are available at https://github.com/xuyungang/DyMoTree. All the data used in this study and the source data for reproducing the results are available at Zenodo (https://zenodo.org/records/20392536).

## Acknowledgments

This work was supported by the National Natural Science Foundation of China (62471378, 82541006, and 62171365), the Young Talent Support Plan of Xi’an Jiaotong University (YX6J021), and Shaanxi Province Key Research and Development Projects (2024SF-GJHX-40 and QCYRCXM-2022-209).

## Extended Data

**Extended Data Fig. 1.**
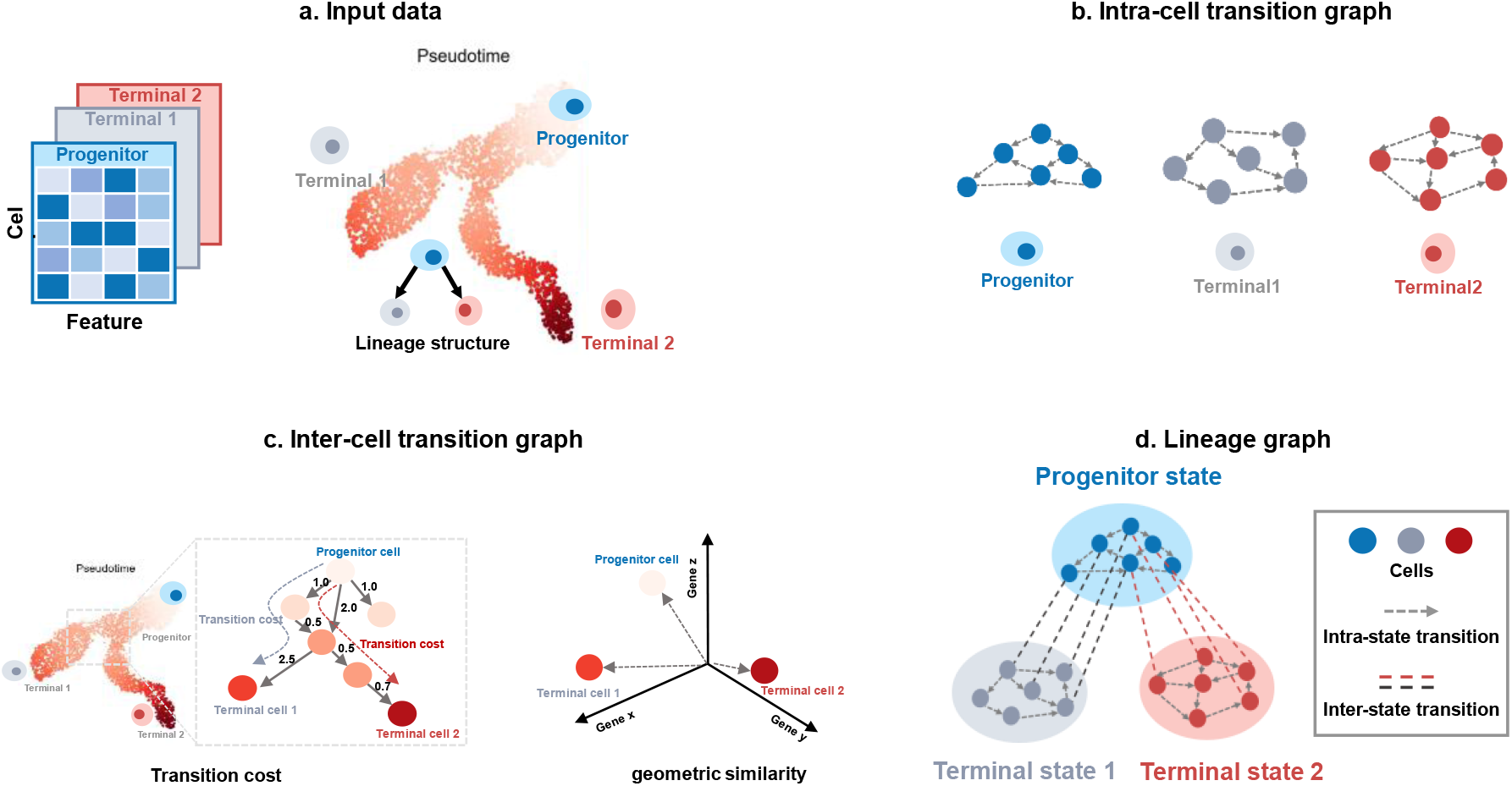
Conceptual workflow for constructing the lineage graph from single-cell data. Gene expression profiles serve as the molecular basis for defining cellular states and revealing lineage relationships among progenitor and progeny populations. Based on these data, potential inter-lineage transitions between cells from adjacent lineage levels are quantified by combining transition cost and geometric similarity in the gene expression space. Within each cell type, directed k-nearest neighbor (KNN) graphs describe bidirectional, local state transitions among cells at the same lineage level. Integrating these intra-type and inter-lineage transition graphs yields a unified lineage graph with single-cell resolution, which captures the hierarchical organization and directional flow of state transitions across the lineage tree.

**Extended Data Fig. 2.**
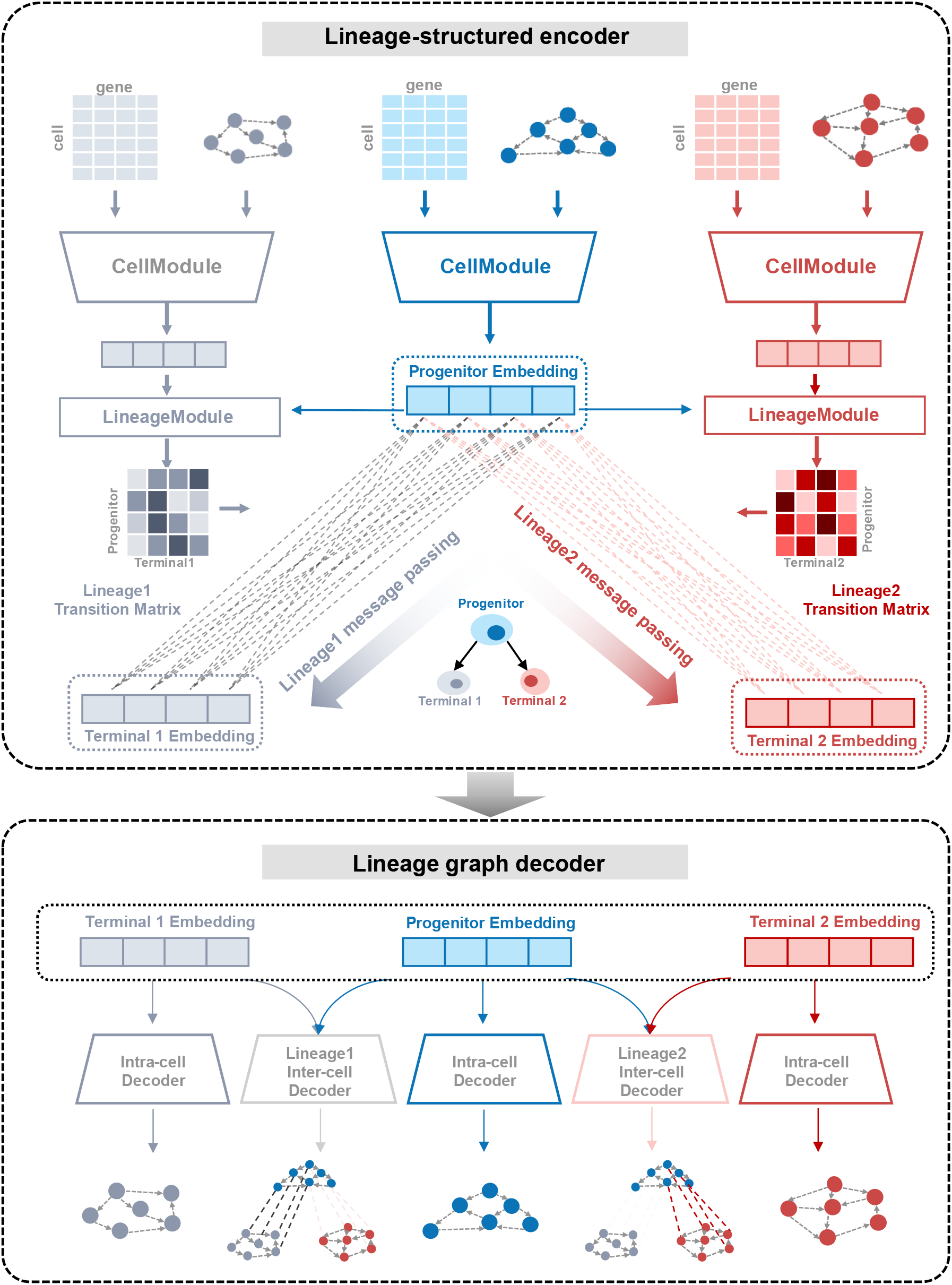
Core neural network architecture of DyMoTree. The lineage-structured encoder receives intra-state transition graphs and gene expression profiles for each cell state as input and employs graph attention network (GAT)-based CellModules to learn cell-state transition representations. Terminal cell embeddings are predicted from progenitor cell embeddings through LineageModules, which integrate terminal cell features from terminal CellModules with progenitor embeddings to model progenitor-to-terminal mappings. The lineage graph decoder contains two types of sub-decoders: an intra-state decoder for reconstructing intra-state graphs and an inter-state decoder for reconstructing inter-state transition graphs.

**Extended Data Fig. 3.**
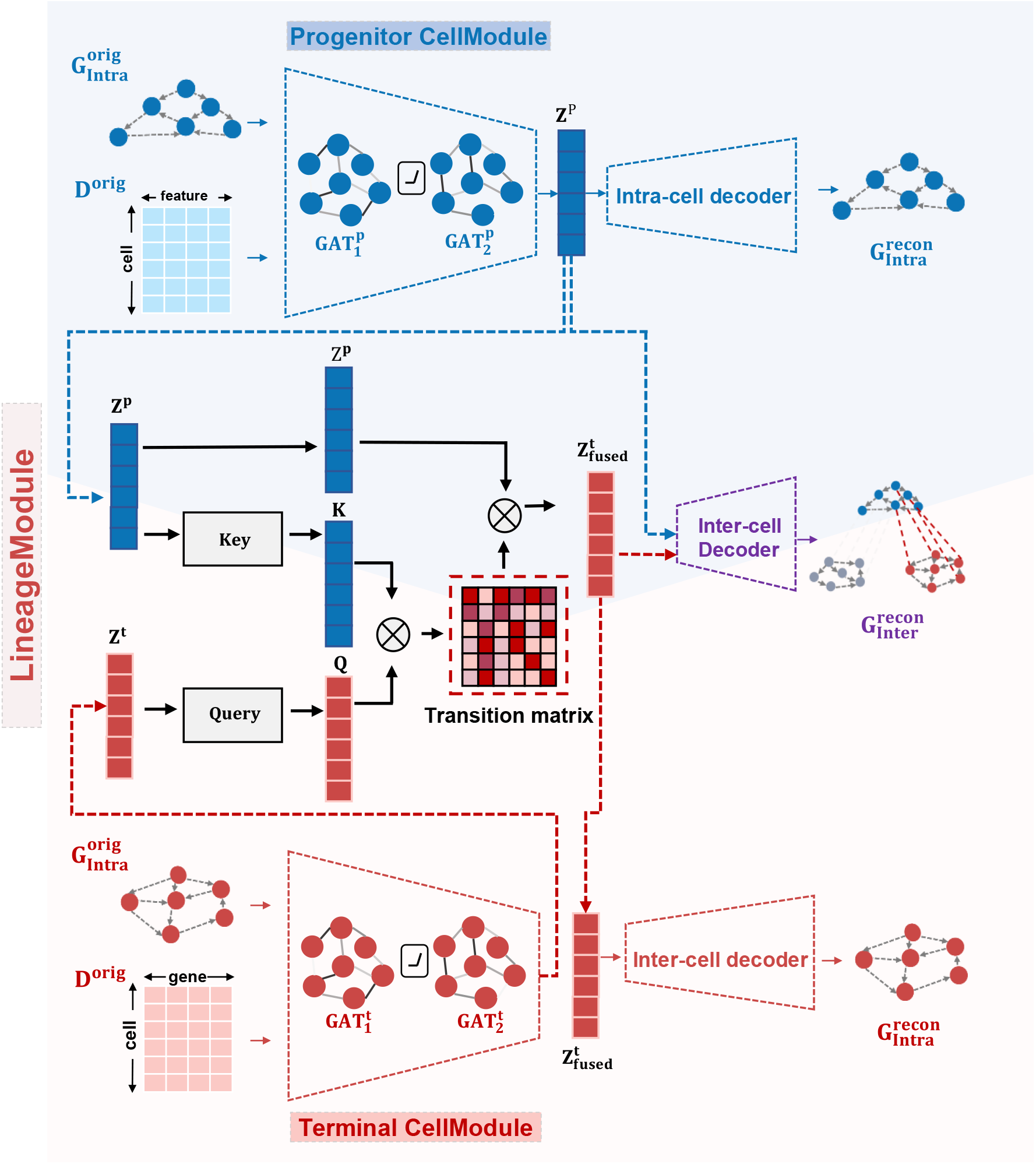
Model components and information flow. DyMoTree consists of three interconnected sub-networks corresponding to different levels of the lineage hierarchy: the Progenitor network, the Lineage network, and the Progeny network. Each Progenitor and Progeny network contains two graph attention modules (GATs) followed by a decoder to reconstruct their respective intra-state transition graphs 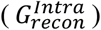, allowing each CellModule to learn local transition patterns within its cell type. The Lineage network connects the two cell-type networks through a cross-attention module that projects parent (progenitor) embeddings into descendant (progeny) representations. The resulting cross-attention matrix (core output) represents the learned lineage-pair–level state transition matrix and is used by the Lineage decoder to reconstruct the inter-lineage bipartite graph 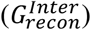. Together, these modules enable DyMoTree to capture both local and hierarchical state-transition relationships across the entire lineage structure.

**Extended Data Fig. 4.**
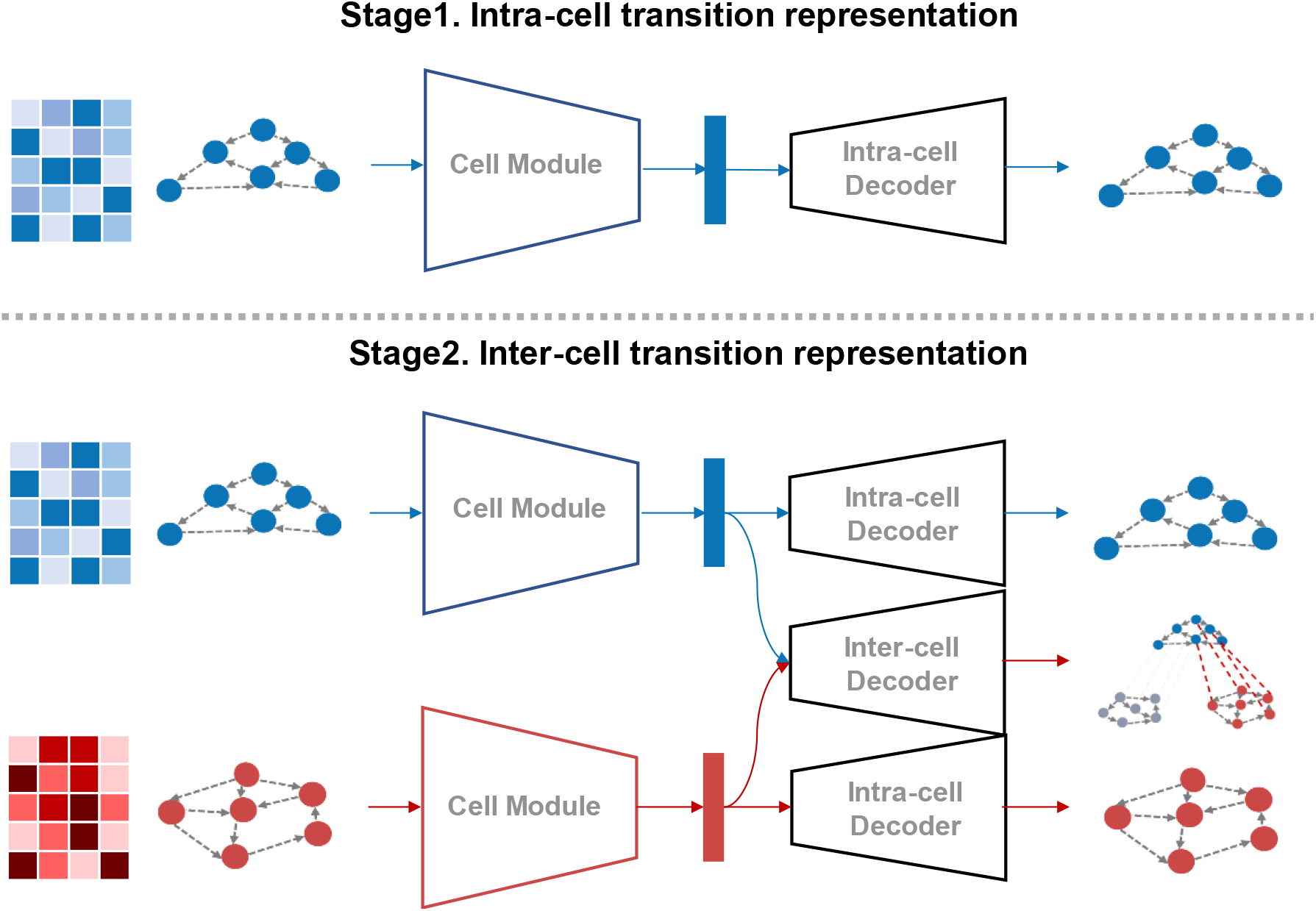
Two-stage pretraining strategy for the CellModule. In Stage 1, each CellModule is trained to reconstruct intra-state transition graphs, capturing local transition patterns within individual cell states. In Stage 2, reconstruction of inter-state bipartite transition graphs between lineage pairs is introduced, enabling coordinated learning of both local and cross-state transition dynamics.

**Extended Data Fig. 5.**
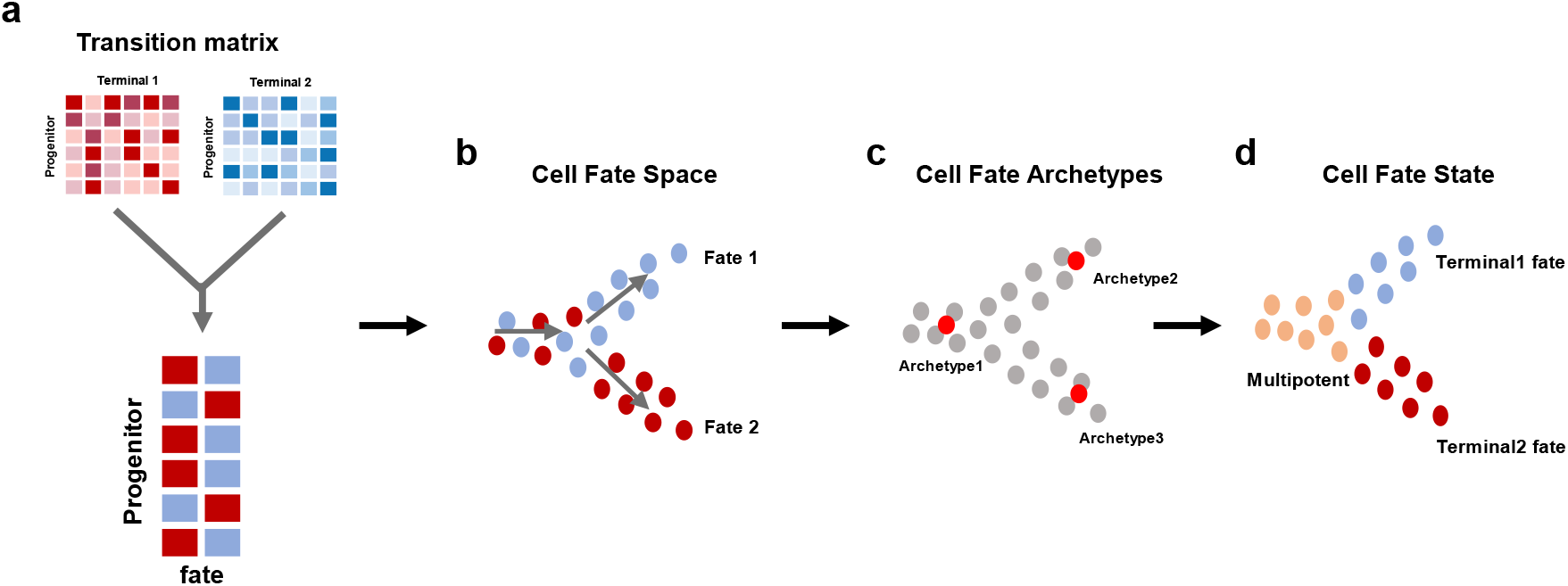
Identification of fate-specific progenitor states based on archetype analysis. The lineage-pair transition matrices learned by DyMoTree are aggregated into a fate potential matrix representing the projection of progenitor cells from gene expression space into a structured fate space. Within this space, progenitor cells are distributed along continuous trajectories corresponding to distinct differentiation directions (fates). To approximate the overall organization of the fate manifold, DyMoTree performs archetype analysis to identify a limited number of representative prototype points (cell fate archetypes) that capture the major fate directions. Each progenitor cell is then expressed as a probabilistic combination of these prototypes, allowing the decomposition of the progenitor population into fate-specific substates. This analysis reveals multipotent and lineage-committed progenitor states corresponding to distinct differentiation outcomes.

**Extended Data Fig. 6.**
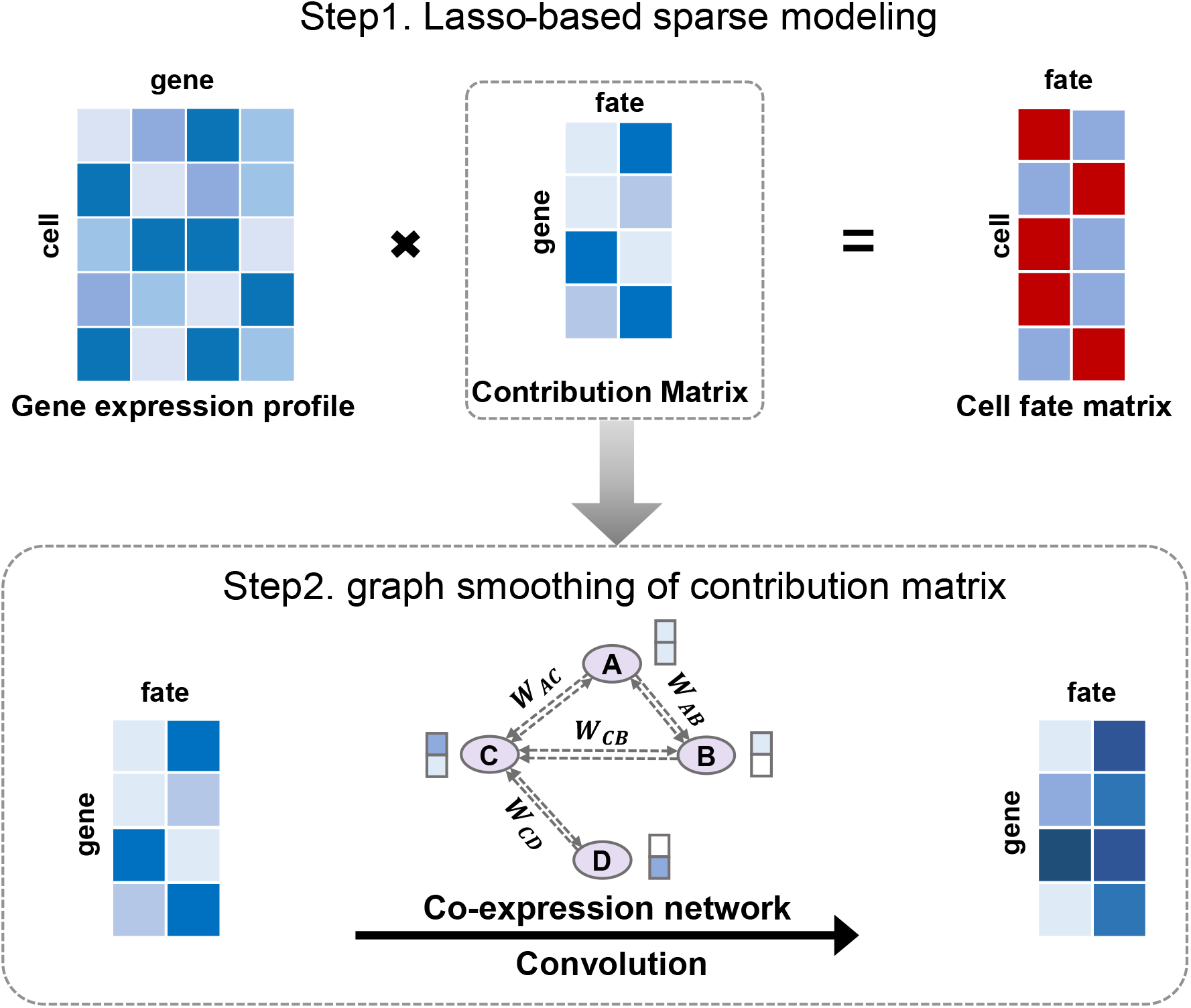
Schematic of candidate driver gene identification. DyMoTree employs a two-step strategy to integrate fate potential with gene expression for identifying lineage-specific genes. **Step 1:** Lasso-based sparse modeling regresses candidate gene expression profiles against cell fate potential scores to obtain a sparse gene contribution matrix. **Step 2:** The Lasso-derived coefficients are refined by graph convolutional smoothing over a gene co-expression network. The contribution score of each gene represents a node feature in the co-expression network, propagating contribution scores among correlated genes to reduce single-cell noise and improve robustness. Genes are finally ranked by their smoothed contribution scores, and top-ranked genes are defined as lineage-specific.

**Extended Data Fig. 7.**
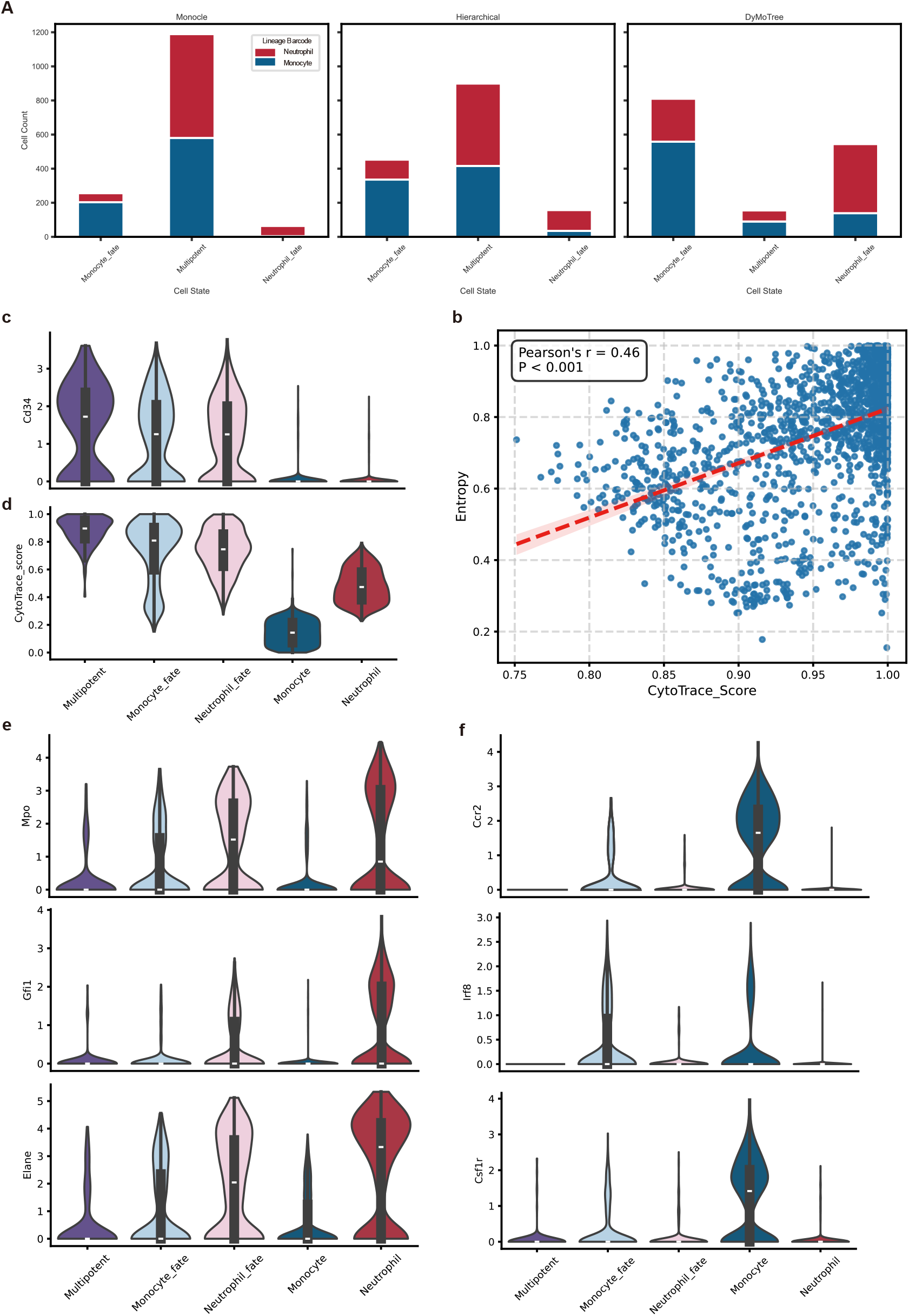
Benchmarking of fate-specific progenitor state identification. **(a)** Comparison of fate-biased HSPCs identified by different approaches. DyMoTree identified a greater number of fate-biased progenitor cells, particularly within the monocyte and neutrophil branches, most of which carried corresponding lineage barcodes. **(b–c)** Violin plots showing the expression of the stemness marker **Cd34 (b)** and CytoTrace-derived differentiation potential **(c)** across identified fate-specific states. Both Cd34 expression and CytoTrace scores were highest in the multipotent population. **(d)** Correlation between DyMoTree-derived entropy of fate potential distributions and CytoTrace differentiation potential (Pearson’s r = 0.46, P < 0.001), indicating that entropy serves as a reliable proxy for differentiation potential. **(e–f)** Expression patterns of representative fate-determining genes in monocyte and neutrophil lineages, including Gfi1, Irf8, Cebpe, Ccr2, and Csf1, showing lineage-specific upregulation in the corresponding fate states.

## References

1. Beumer, J. & Clevers, H. Cell fate specification and differentiation in the adult mammalian intestine. Nat Rev Mol Cell Biol 22, 39–53 (2021).

2. De Belly, H., Paluch, E.K. & Chalut, K.J. Interplay between mechanics and signalling in regulating cell fate. Nat Rev Mol Cell Biol 23, 465–480 (2022).

3. Moris, N., Pina, C. & Arias, A.M. Transition states and cell fate decisions in epigenetic landscapes. Nat Rev Genet 17, 693–703 (2016).

4. Wagner, D.E. & Klein, A.M. Lineage tracing meets single-cell omics: opportunities and challenges. Nat Rev Genet 21, 410–427 (2020).

5. de Visser, K.E. & Joyce, J.A. The evolving tumor microenvironment: From cancer initiation to metastatic outgrowth. Cancer Cell 41, 374–403 (2023).

6. Weinreb, C., Rodriguez-Fraticelli, A., Camargo, F.D. & Klein, A.M. Lineage tracing on transcriptional landscapes links state to fate during differentiation. Science 367 (2020).

7. Griffiths, J.A., Scialdone, A. & Marioni, J.C. Using single-cell genomics to understand developmental processes and cell fate decisions. Mol Syst Biol 14, e8046 (2018).

8. Kester, L. & van Oudenaarden, A. Single-Cell Transcriptomics Meets Lineage Tracing. Cell Stem Cell 23, 166–179 (2018).

9. Heumos, L. et al. Best practices for single-cell analysis across modalities. Nat Rev Genet 24, 550–572 (2023).

10. Herman, J.S. Sagar & Grun, D. FateID infers cell fate bias in multipotent progenitors from single-cell RNA-seq data. Nat Methods 15, 379–386 (2018).

11. Kong, W. et al. Capybara: A computational tool to measure cell identity and fate transitions. Cell Stem Cell 29, 635–649 e611 (2022).

12. Lange, M. et al. CellRank for directed single-cell fate mapping. Nat Methods 19, 159–170 (2022).

13. Pandey, K. & Zafar, H. Inference of cell state transitions and cell fate plasticity from single-cell with MARGARET. Nucleic Acids Res 50, e86 (2022).

14. Setty, M. et al. Characterization of cell fate probabilities in single-cell data with Palantir. Nat Biotechnol 37, 451–460 (2019).

15. Weiler, P., Lange, M., Klein, M., Pe’er, D. & Theis, F. CellRank 2: unified fate mapping in multiview single-cell data. Nat Methods 21, 1196–1205 (2024).

16. Guo, W. et al. scStateDynamics: deciphering the drug-responsive tumor cell state dynamics by modeling single-cell level expression changes. Genome Biol 25, 297 (2024).

17. Klein, D. et al. Mapping cells through time and space with moscot. Nature 638, 1065–1075 (2025).

18. Schiebinger, G. et al. Optimal-Transport Analysis of Single-Cell Gene Expression Identifies Developmental Trajectories in Reprogramming. Cell 176, 928–943 e922 (2019).

19. Wang, S.W., Herriges, M.J., Hurley, K., Kotton, D.N. & Klein, A.M. CoSpar identifies early cell fate biases from single-cell transcriptomic and lineage information. Nat Biotechnol 40, 1066–1074 (2022).

20. Street, K. et al. Slingshot: cell lineage and pseudotime inference for single-cell transcriptomics. BMC Genomics 19, 477 (2018).

21. Trapnell, C. et al. The dynamics and regulators of cell fate decisions are revealed by pseudotemporal ordering of single cells. Nat Biotechnol 32, 381–386 (2014).

22. Guo, W. et al. scTrace+: Enhancing cell fate inference by integrating the lineage-tracing and multi-faceted transcriptomic similarity information. Cell Syst 16, 101398 (2025).

23. Rafelski, S.M. & Theriot, J.A. Establishing a conceptual framework for holistic cell states and state transitions. Cell 187, 2633–2651 (2024).

24. HajiAkhondi-Meybodi, Z., Mohammadi, A., Hou, M., Abouei, J. & Plataniotis, K.N. Vit-Cat: Parallel Vision Transformers with Cross Attention Fusion for Popularity Prediction in Mec Networks. Int Conf Acoust Spee (2023).

25. Battich, N. et al. Sequencing metabolically labeled transcripts in single cells reveals mRNA turnover strategies. Science 367, 1151–1156 (2020).

26. Bergen, V., Lange, M., Peidli, S., Wolf, F.A. & Theis, F.J. Generalizing RNA velocity to transient cell states through dynamical modeling. Nat Biotechnol 38, 1408–1414 (2020).

27. La Manno, G. et al. RNA velocity of single cells. Nature 560, 494–498 (2018).

28. Ren, J. et al. Spatiotemporally resolved transcriptomics reveals the subcellular RNA kinetic landscape. Nat Methods 20, 695–705 (2023).

29. Veličković, P. et al. arXiv:1710.10903 (2017).

30. Lin, H. et al. arXiv:2106.05786 (2021).

31. Persad, S. et al. SEACells infers transcriptional and epigenomic cellular states from single-cell genomics data. Nat Biotechnol 41, 1746–1757 (2023).

32. Wang, T. et al. CellNavi predicts genes directing cellular transitions by learning a gene graph-enhanced cell state manifold. Nat Cell Biol 27, 1863–1874 (2025).

33. Alcacer, A., Epifanio, I., Mair, S. & Mørup, M. arXiv:2504.12392 (2025).

34. Gillariose, J., Joseph, J. & Chesneau, C. Lasso and Ridge regression: a comprehensive review of applications and developments in machine learning. Int J Data Sci Anal 21 (2025).

35. Langfelder, P. & Horvath, S. WGCNA: an R package for weighted correlation network analysis. BMC Bioinformatics 9, 559 (2008).

36. Zappia, L., Phipson, B. & Oshlack, A. Splatter: simulation of single-cell RNA sequencing data. Genome Biol 18, 174 (2017).

37. Jiang, J. et al. scLTdb: a comprehensive single-cell lineage tracing database. Nucleic Acids Res 53, D1173–D1185 (2025).

38. Abdullah, L. et al. The endogenous antigen-specific CD8(+) T cell repertoire is composed of unbiased and biased clonotypes with differential fate commitments. Immunity 58, 601–615 e609 (2025).

39. Irac, S.E., Soon, M.S.F., Borcherding, N. & Tuong, Z.K. Single-cell immune repertoire analysis. Nat Methods 21, 777–792 (2024).

40. Cohen-Addad, V., Kanade, V., Mallmann-Trenn, F. & Mathieu, C. Hierarchical Clustering: Objective Functions and Algorithms. Soda’18: Proceedings of the Twenty-Ninth Annual Acm-Siam Symposium on Discrete Algorithms, 378–397 (2018).

41. Gulati, G.S. et al. Single-cell transcriptional diversity is a hallmark of developmental potential. Science 367, 405–411 (2020).

42. Bessonnard, S. et al. Gata6, Nanog and Erk signaling control cell fate in the inner cell mass through a tristable regulatory network. Development 141, 3637–3648 (2014).

43. Yamanaka, Y., Ralston, A., Stephenson, R.O. & Rossant, J. Cell and molecular regulation of the mouse blastocyst. Dev Dyn 235, 2301–2314 (2006).

44. Nowotschin, S. et al. The emergent landscape of the mouse gut endoderm at single-cell resolution. Nature 569, 361–367 (2019).

45. Plusa, B., Piliszek, A., Frankenberg, S., Artus, J. & Hadjantonakis, A.K. Distinct sequential cell behaviours direct primitive endoderm formation in the mouse blastocyst. Development 135, 3081–3091 (2008).

46. Schrode, N., Saiz, N., Di Talia, S. & Hadjantonakis, A.K. GATA6 levels modulate primitive endoderm cell fate choice and timing in the mouse blastocyst. Dev Cell 29, 454–467 (2014).

47. Xenopoulos, P., Kang, M., Puliafito, A., Di Talia, S. & Hadjantonakis, A.K. Heterogeneities in Nanog Expression Drive Stable Commitment to Pluripotency in the Mouse Blastocyst. Cell Rep 10, 1508–1520 (2015).

48. Kang, M., Garg, V. & Hadjantonakis, A.K. Lineage Establishment and Progression within the Inner Cell Mass of the Mouse Blastocyst Requires FGFR1 and FGFR2. Dev Cell 41, 496–510 e495 (2017).

49. Bravo Gonzalez-Blas, C. et al. SCENIC+: single-cell multiomic inference of enhancers and gene regulatory networks. Nat Methods 20, 1355–1367 (2023).

50. Wang, P. et al. Deciphering driver regulators of cell fate decisions from single-cell transcriptomics data with CEFCON. Nat Commun 14, 8459 (2023).

51. Yuan, Q. & Duren, Z. Inferring gene regulatory networks from single-cell multiome data using atlas-scale external data. Nat Biotechnol 43, 247–257 (2025).

52. Geiselmann, A. et al. PI3K/AKT signaling controls ICM maturation and proper epiblast and primitive endoderm specification in mice. Dev Cell 60, 204–219 e206 (2025).

53. Perez-Gonzalez, A., Bevant, K. & Blanpain, C. Cancer cell plasticity during tumor progression, metastasis and response to therapy. Nat Cancer 4, 1063–1082 (2023).

54. Marjanovic, N.D. et al. Emergence of a High-Plasticity Cell State during Lung Cancer Evolution. Cancer Cell 38, 229–246 e213 (2020).

55. Kaiser, A.M. et al. p53 governs an AT1 differentiation programme in lung cancer suppression. Nature 619, 851–859 (2023).

56. Gene Ontology, C. The Gene Ontology knowledgebase in 2026. Nucleic Acids Res 54, D1779–D1792 (2026).

57. Li, Z. et al. Functional diversification and dynamics of CAR-T cells in patients with B-ALL. Cell Rep 42, 113263 (2023).

58. Weng, N.P., Araki, Y. & Subedi, K. The molecular basis of the memory T cell response: differential gene expression and its epigenetic regulation. Nat Rev Immunol 12, 306–315 (2012).

59. Pais Ferreira, D. et al. Central memory CD8(+) T cells derive from stem-like Tcf7(hi) effector cells in the absence of cytotoxic differentiation. Immunity 53, 985–1000 e1011 (2020).

60. Balin, S.J. et al. Human antimicrobial cytotoxic T lymphocytes, defined by NK receptors and antimicrobial proteins, kill intracellular bacteria. Sci Immunol 3 (2018).

61. Meylan, M. et al. Persistent T cell activation and cytotoxicity against glioblastoma following single oncolytic virus treatment in a clinical trial. Cell 189, 1287–1304 e1218 (2026).

62. Kye, Y.C. et al. STAT1 maintains naive CD8(+) T cell quiescence by suppressing the type I IFN-STAT4-mTORC1 signaling axis. Sci Adv 7, eabg8764 (2021).

63. Chan, J.D. et al. FOXO1 enhances CAR T cell stemness, metabolic fitness and efficacy. Nature 629, 201–210 (2024).

64. Doan, A.E. et al. FOXO1 is a master regulator of memory programming in CAR T cells. Nature 629, 211–218 (2024).

65. Wu, Y. et al. Spatiotemporal Immune Landscape of Colorectal Cancer Liver Metastasis at Single-Cell Level. Cancer Discov 12, 134–153 (2022).

66. Waikhom, L. & Patgiri, R. arXiv:2108.10733 (2021).

