## Supplementary information for "DyMoTree decodes early cell state transitions and drivers from single-cell transcriptomes using a tree-structured neural network"

### Supplementary Figures 1-19 and Tables 1-4

#### DyMoTree decodes early cell state transitions and drivers from single-cell transcriptomes using a tree-structured neural network

Jiayi Wang[1], Rufeng Li[1], Chen Guo[1], Min Qiang[1], Shuai Wang[1],
Genhui Wang[1], Kangsheng Tu[2], Yungang Xu[1,*]

[1] Department of Cell Biology and Genetics, School of Basic Medical Sciences, Xi'an Jiaotong University Health Science Center, Xi'an, Shaanxi 710061, China.

[2] Department of Hepatobiliary Surgery, The First Affiliated Hospital of Xi'an Jiaotong University, Xi'an, Shaanxi, 710061, China

Jiayi Wang,

Rufeng Li,

Chen Guo,

Min Qiang,

Shuai Wang,
Genhui Wang,
Kangsheng Tu,
Yungang Xu,

**Keywords:** single-cell transcriptomics; cell fate inference; lineage-resolved modeling; cell state transitions; tree-structured neural network; driver gene discovery

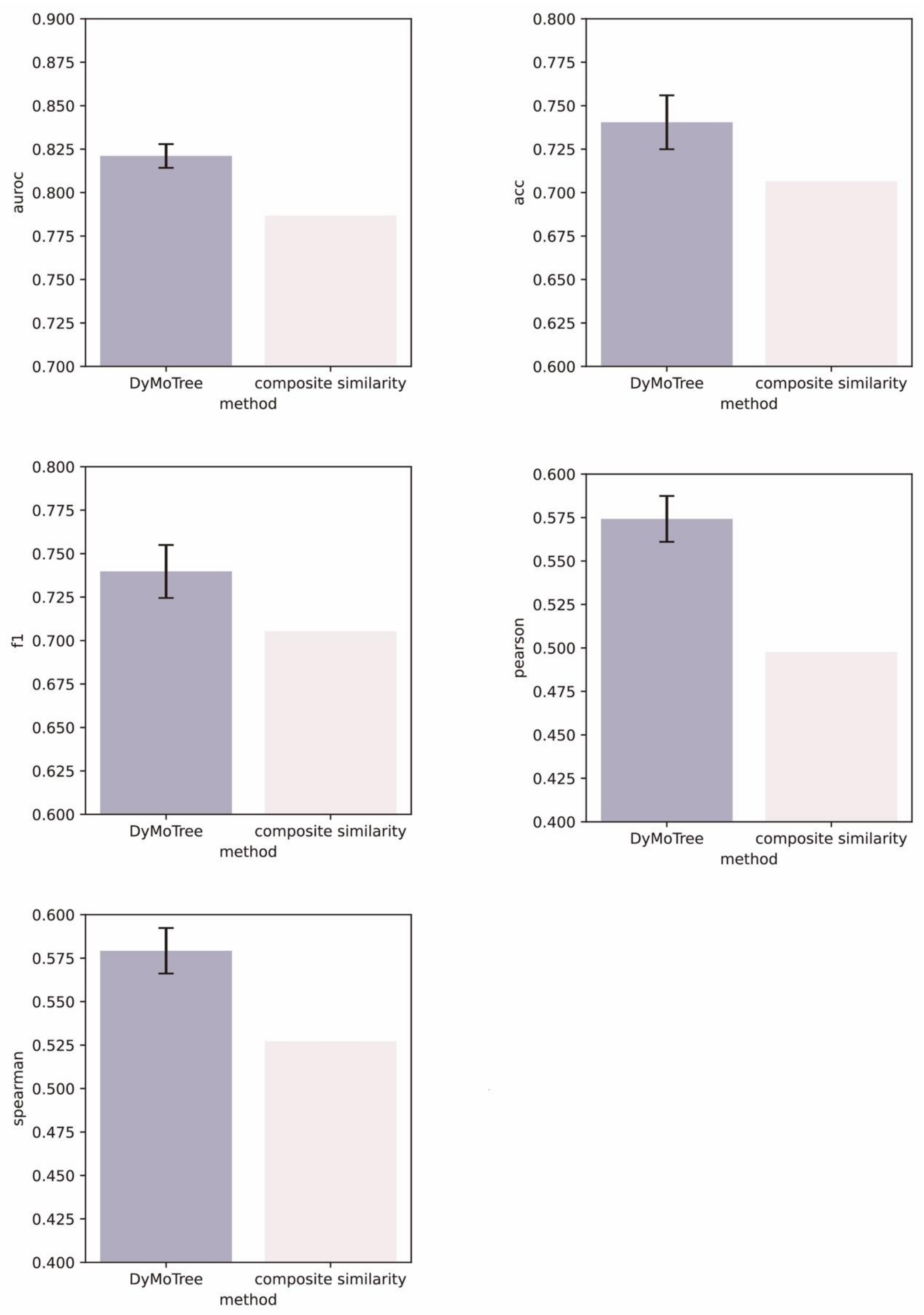


**Supplementary Fig. 1 Comparison between DyMoTree and composite similarity on lineage tracing datasets.**

Performance comparison of DyMoTree and a composite similarity-based method across multiple metrics (AUROC, accuracy, F1-score, Pearson, and Spearman correlations).

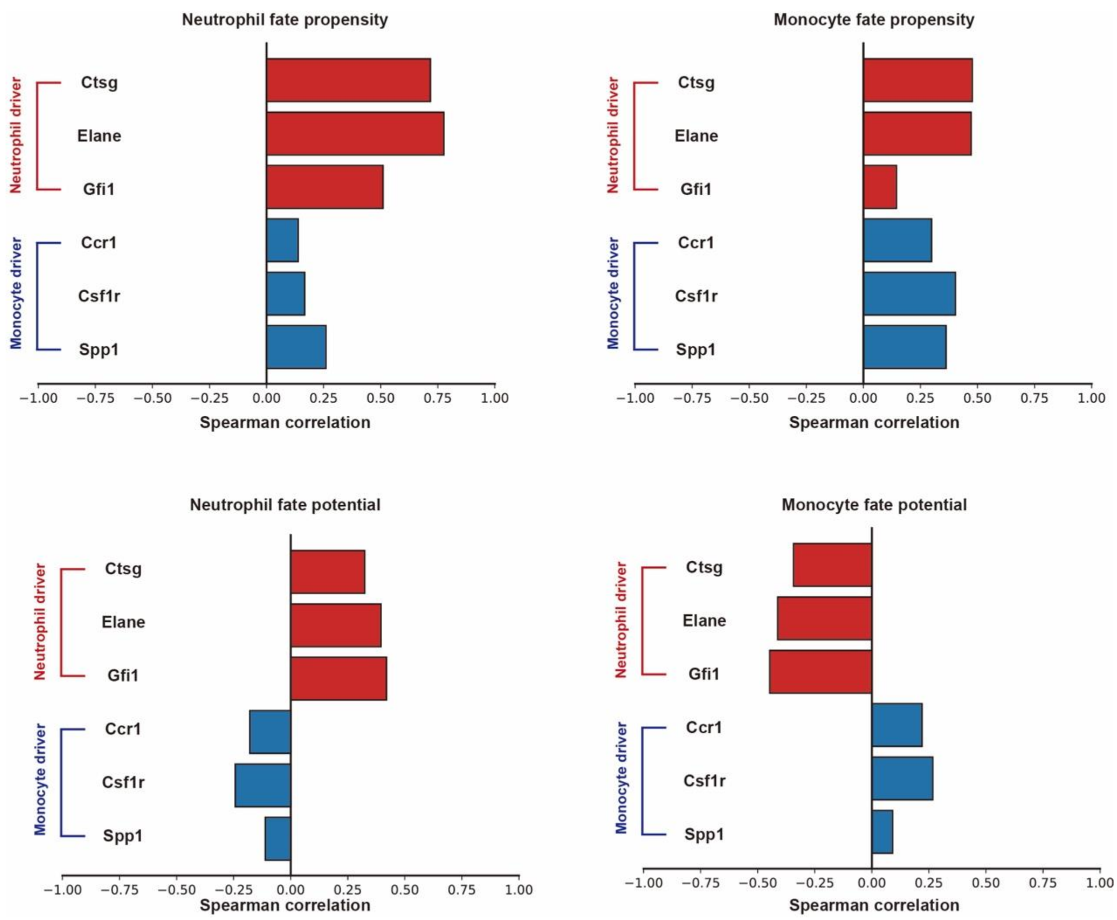


**Supplementary Fig. 2 Correlation of inferred fate scores with lineage driver genes.**
Spearman correlations between gene expression of known driver genes and inferred fate scores for neutrophil and monocyte lineages. The top panels show results based on linear composite similarity (fate propensity), and the bottom panels show results from DyMoTree (fate potential). DyMoTree improves lineage-specific consistency with known driver genes.

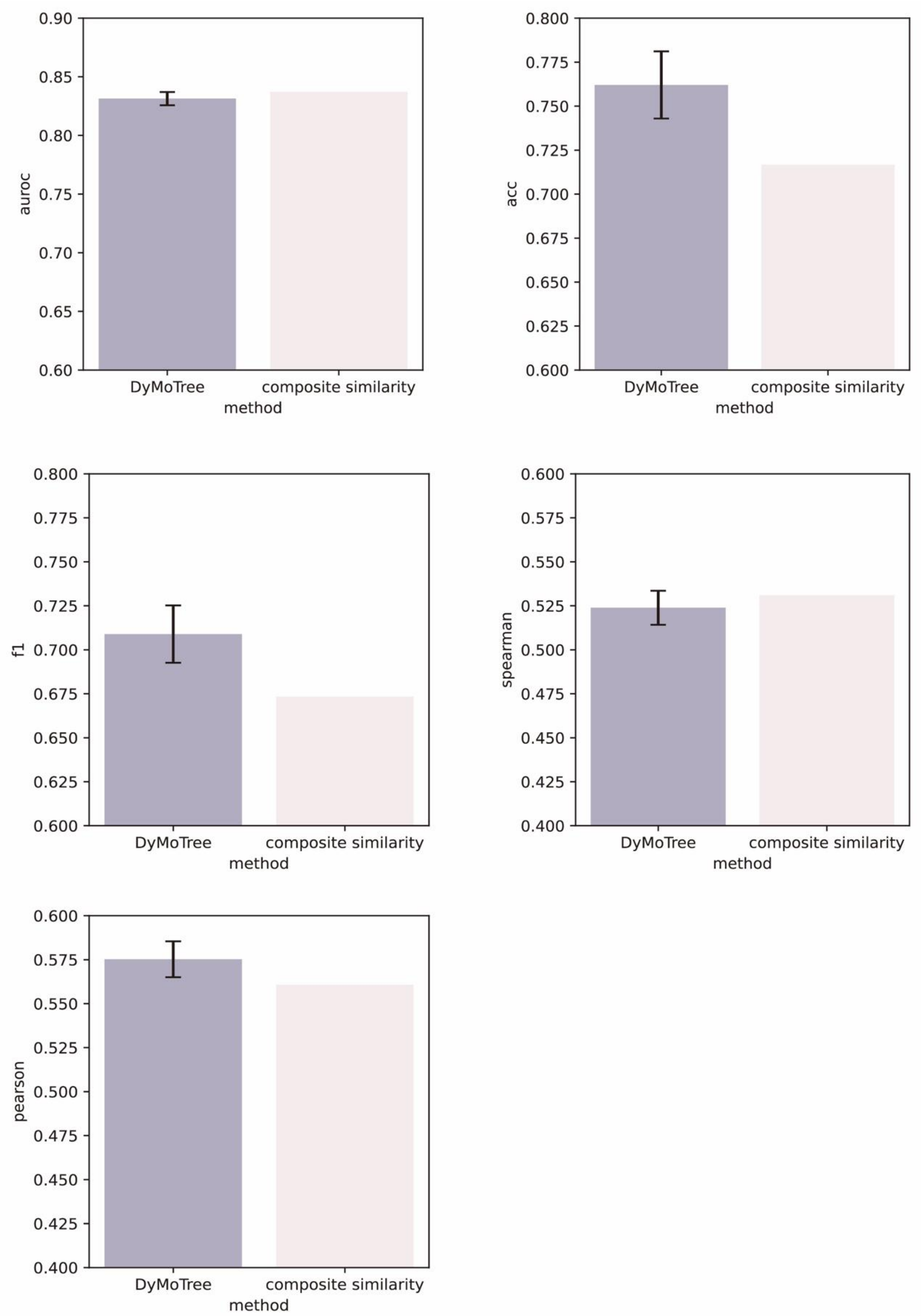


**Supplementary Fig. 3 Comparison between DyMoTree and composite similarity on CD8+ T cell differentiation datasets.**

Performance comparison of DyMoTree and a composite similarity-based method across multiple metrics (AUROC, accuracy, F1-score, Pearson, and Spearman correlations).

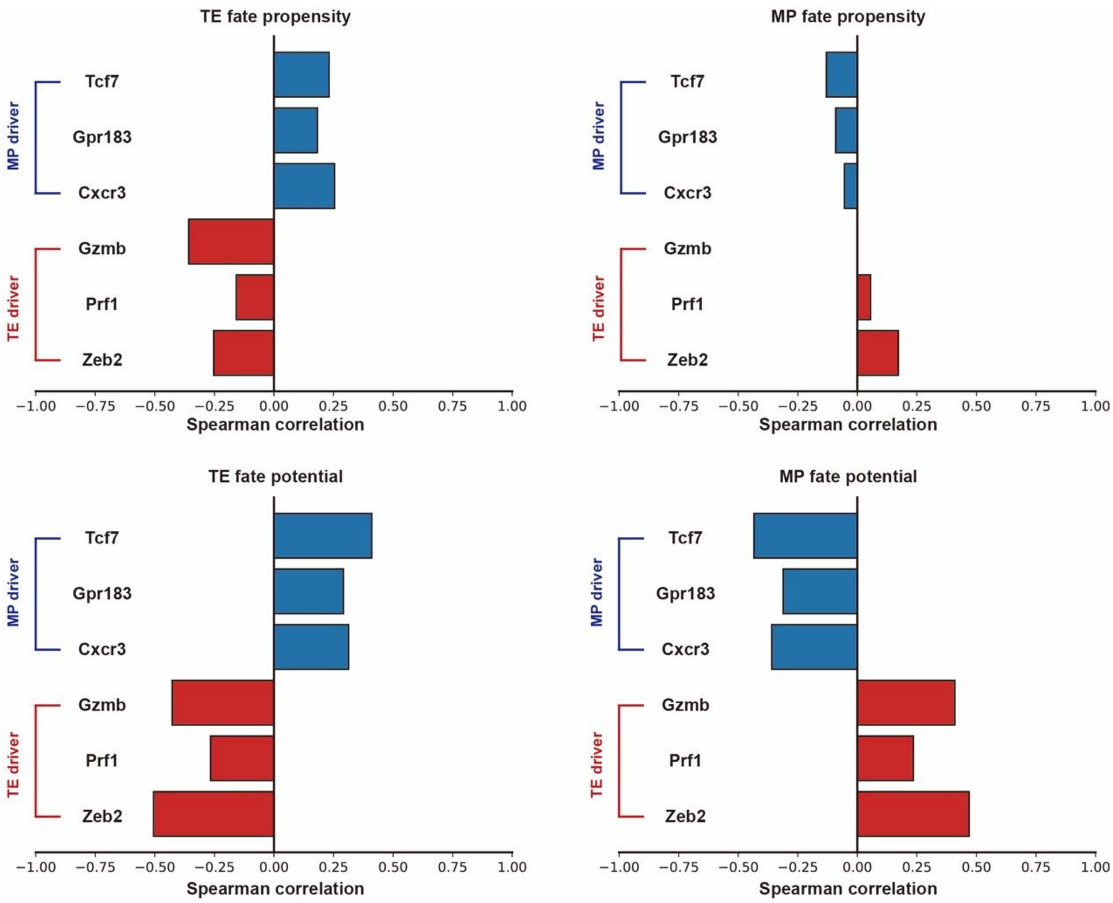


**Supplementary Fig. 4 Correlation of inferred fate scores with lineage driver genes.**

Spearman correlations between gene expression of MP (Tcf7, Gpr183, Cxcr3) and TE (Gzmb, Prf1, Zeb2) driver genes and inferred fate scores. The top panels show fate propensity based on linear similarity, and the bottom panels show DyMoTree-derived fate potential. DyMoTree enhances lineage-specific associations.

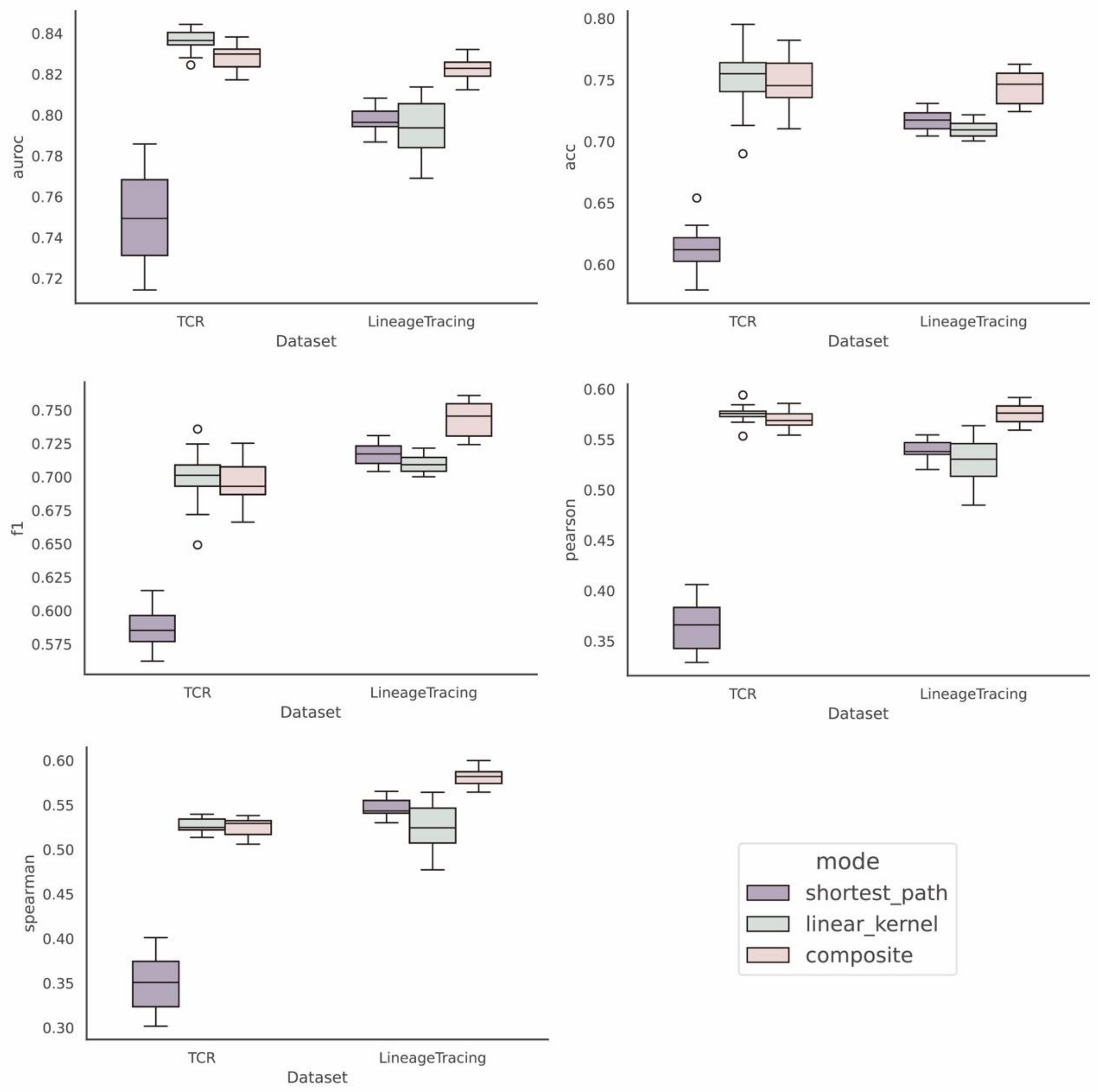


**Supplementary Fig. 5 Effect of different similarity modeling on DyMoTree performance.** Performance comparison of different similarity modes (shortest path, linear kernel, and composite) across TCR and lineage-tracing datasets, evaluated by AUROC, accuracy, F1-score, Pearson, and Spearman correlations.

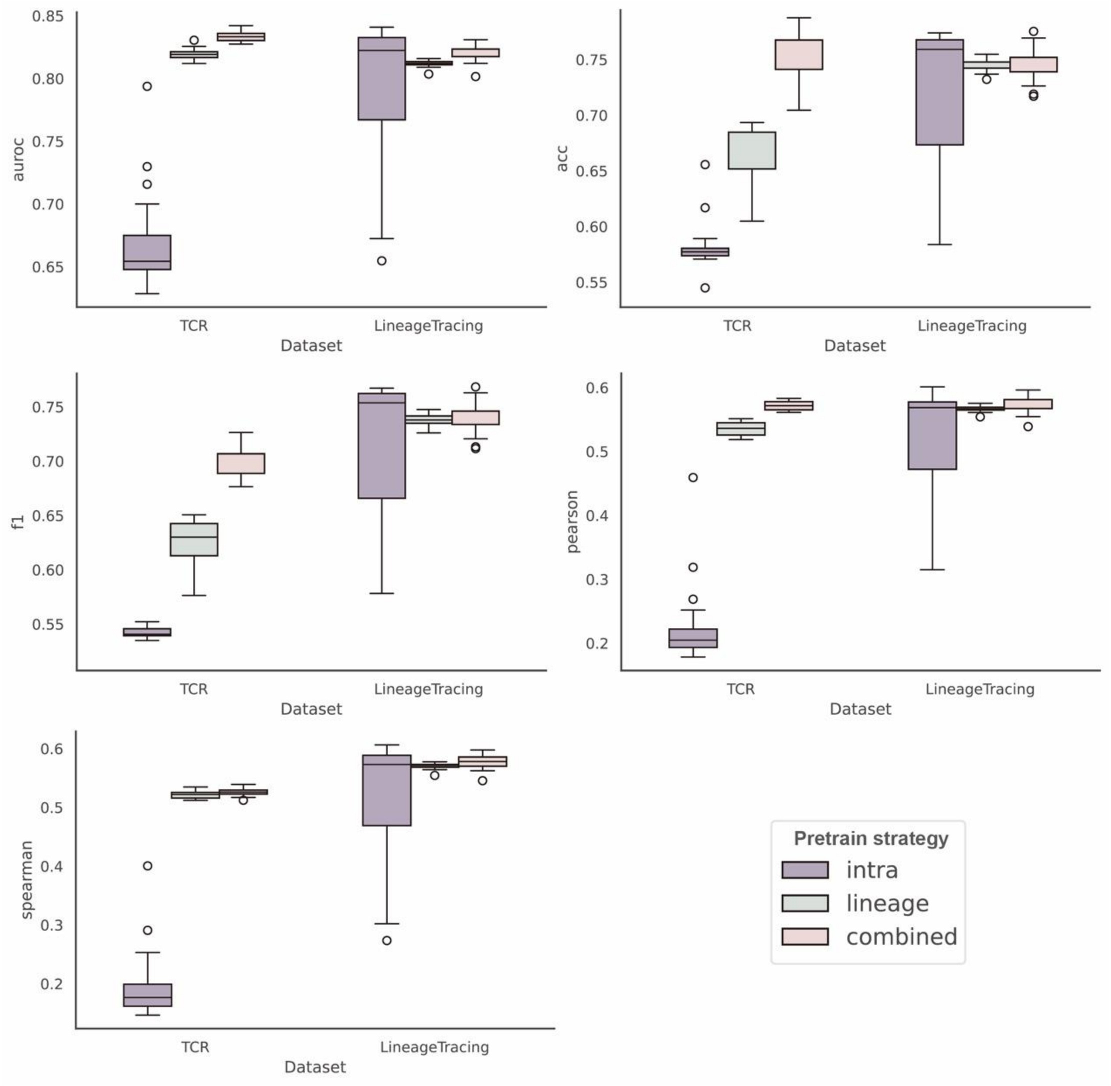


**Supplementary Fig. 6 Impact of pretraining strategy on DyMoTree performance.** Performance comparison of different pretraining strategies (intra-state only, lineage-level, and combined) across TCR and lineage-tracing datasets, evaluated by AUROC, accuracy, F1-score, Pearson, and Spearman correlations.

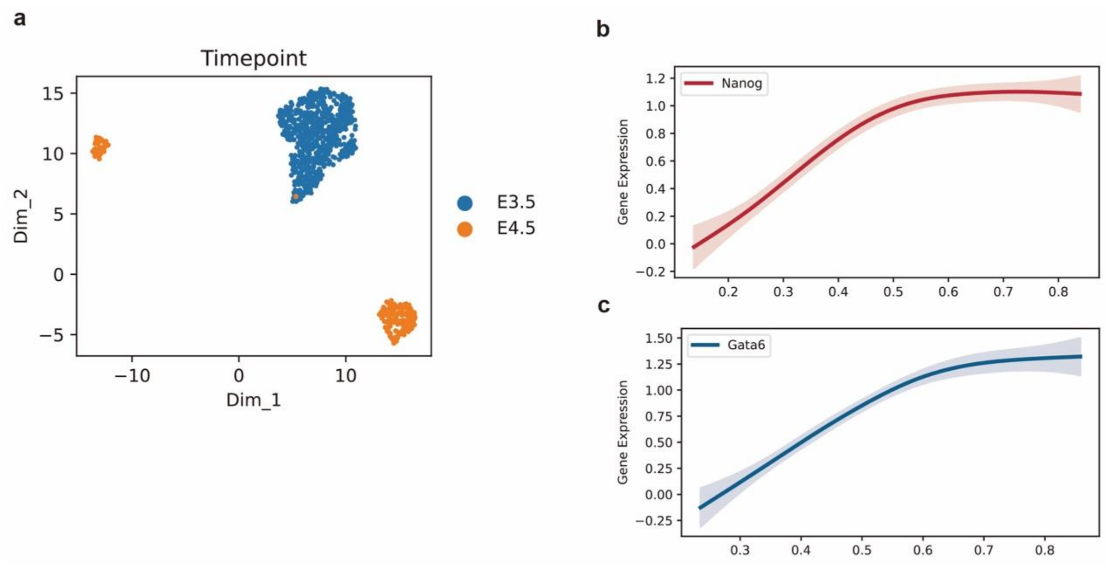


**Supplementary Fig. 7 Detailed transitional dynamics of ICM bifurcation inferred by DyMoTree.**

**(a)** UMAP visualization of mouse early embryogenesis during E3.5-E4.5. **(b)** Gene expression trends of Nanog (a key regulator of epiblast differentiation) along Epi-fate potential, inferred by DyMoTree. **(c)** Gene expression trends of Gata6 (a key regulator of primitive endoderm differentiation) along PrE-fate potential inferred by DyMoTree.

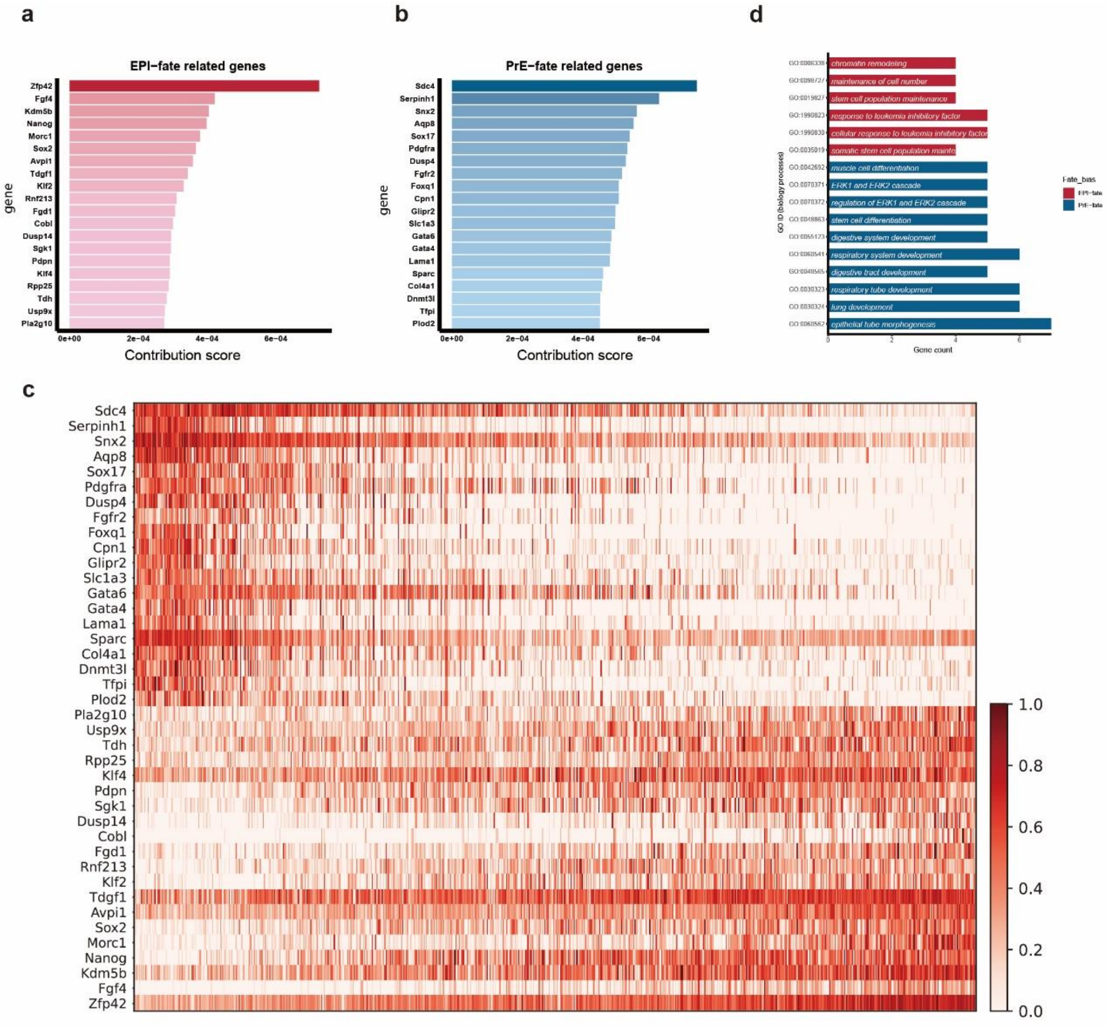


**Supplementary Fig. 8 Identification of lineage-specific genes for ICM differentiating toward PrE or Epi lineages.**

**(a-b)** Bar plots show the contribution scores of the top-20 genes specific to Epi-biased (a) and PrE-biased (b) fate potentials. **(c)** The heatmap shows the expression of the top-20 fate-specific genes toward Epi-biased and PrE-biased fate potentials in ICM cells. Each row denotes a fate-specific gene, the columns denote ICM cells arranged by increasing order of fate-bias calculated by fate potentials. Cells with fate-bias > 0.5 are Epi-biased, and those with bias < 0.5 are PrE-biased. **(d)** The top enriched GO terms of fate-specific gene sets.

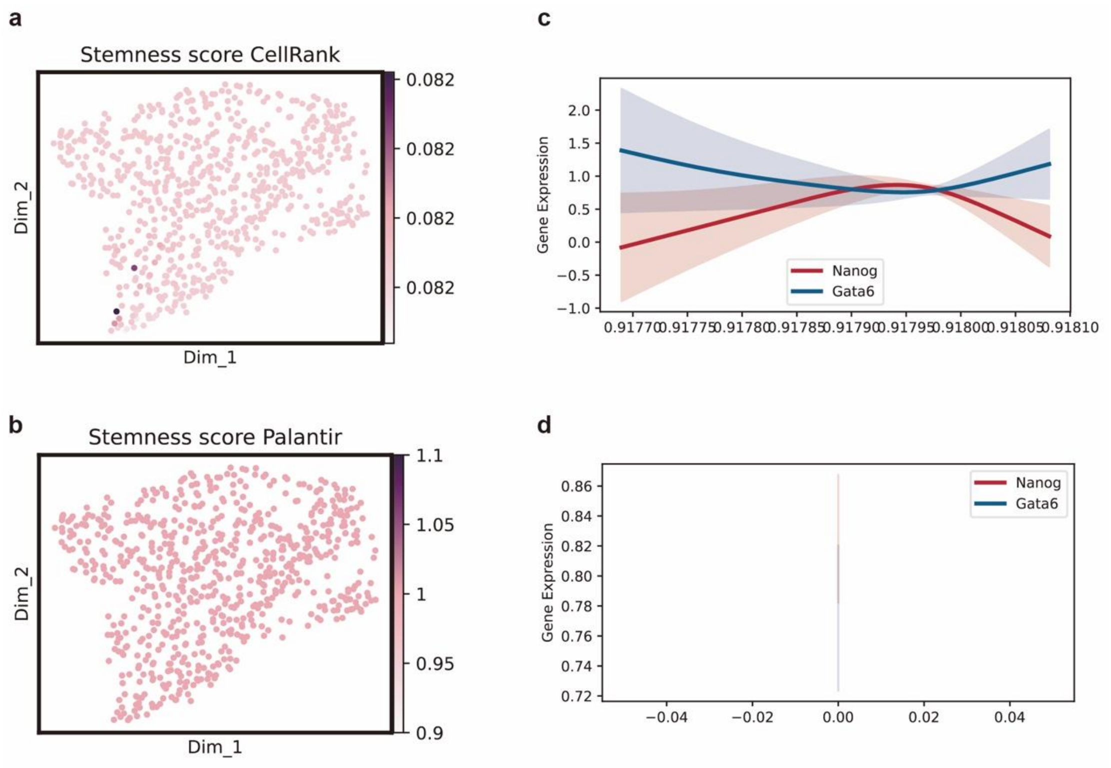


**Supplementary Fig. 9 Stemness score inferred by CellRank and Palantir.**

**(a,b)** Feature plots show stemness score derived from CellRank (a) and Palantir (b). (**c,d**) Expression trends of Gata6 and Nanog in ICM state along differentiation time, approximated by stemness score derived from CellRank (c) and Palantir (d).

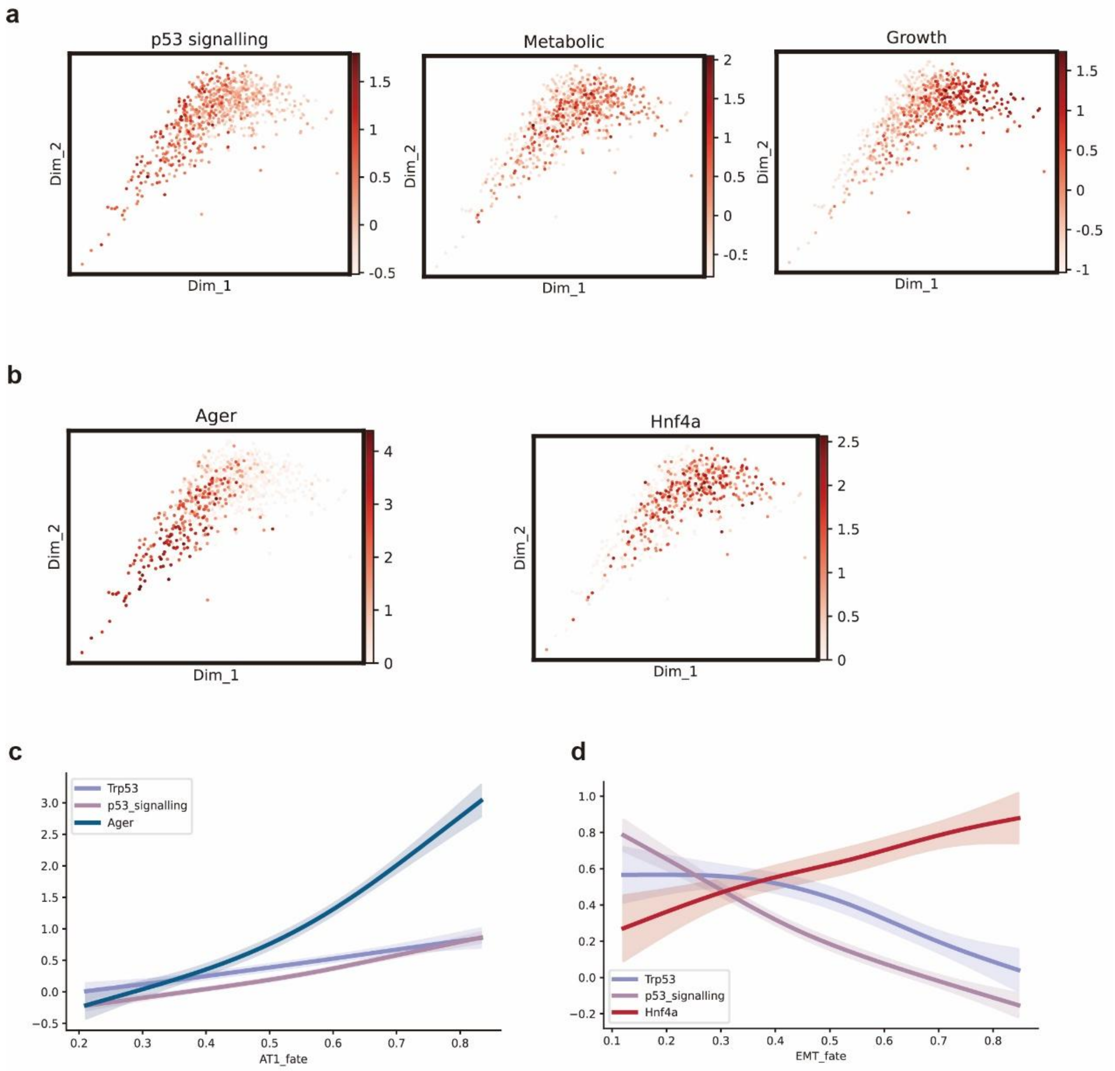


**Supplementary Fig. 10 Expression dynamics of genes and gene sets along DyMoTree-inferred fate potentials.**

**(a)** Aggregate score of three functional gene sets in the transitional state. **(b)** Transcriptomic expression level of Ager and Hnf4a in the transitional state. **(c)** Aggregate score trends of p53 signaling gene set and transcriptomic expression trends of Ager and Trp53 along AT1-biased fate potential in the transitional state. **(d)** Aggregate score trends of p53 signaling gene set and transcriptomic expression trends of Hnf4a and Trp53 along EMT-biased fate potential in the transitional state.

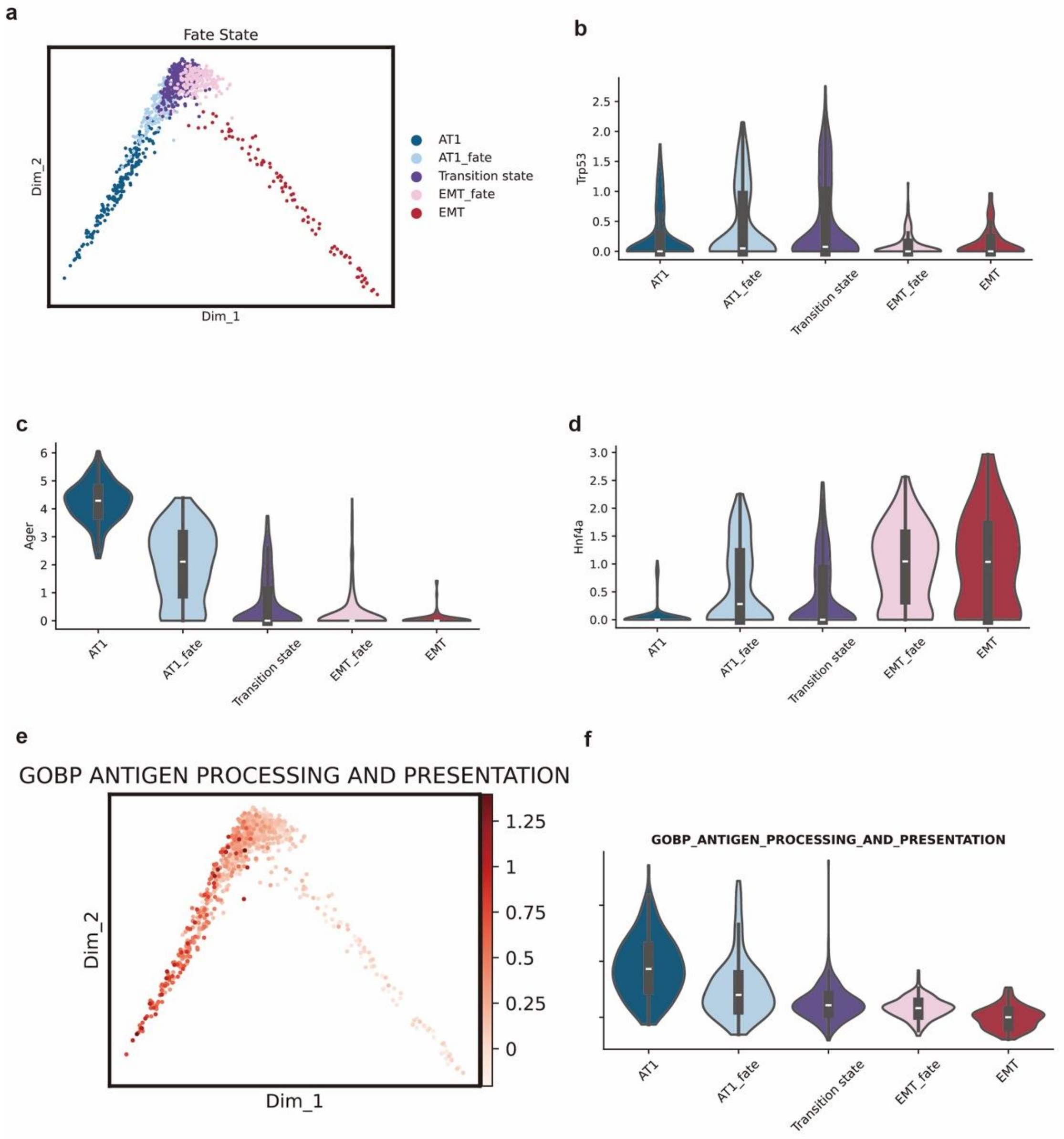


**Supplementary Fig. 11 Validation of fate-specific transitional state identified by DyMoTree.** **(a)** Identification of fate-specific transitional state, including transitional state, AT1-biased state, and EMT-biased state. **(b-d)** Violin plots show the gene expression level of Trp53, Ager, and Hnf4a between LUAD tumor cell substates. **(e)** Aggregate score of antigen processing and presentation gene set in the transitional state. **(f)** Violin plots show the aggregate gene score of the antigen processing and presentation gene set between LUAD tumor cell substates.

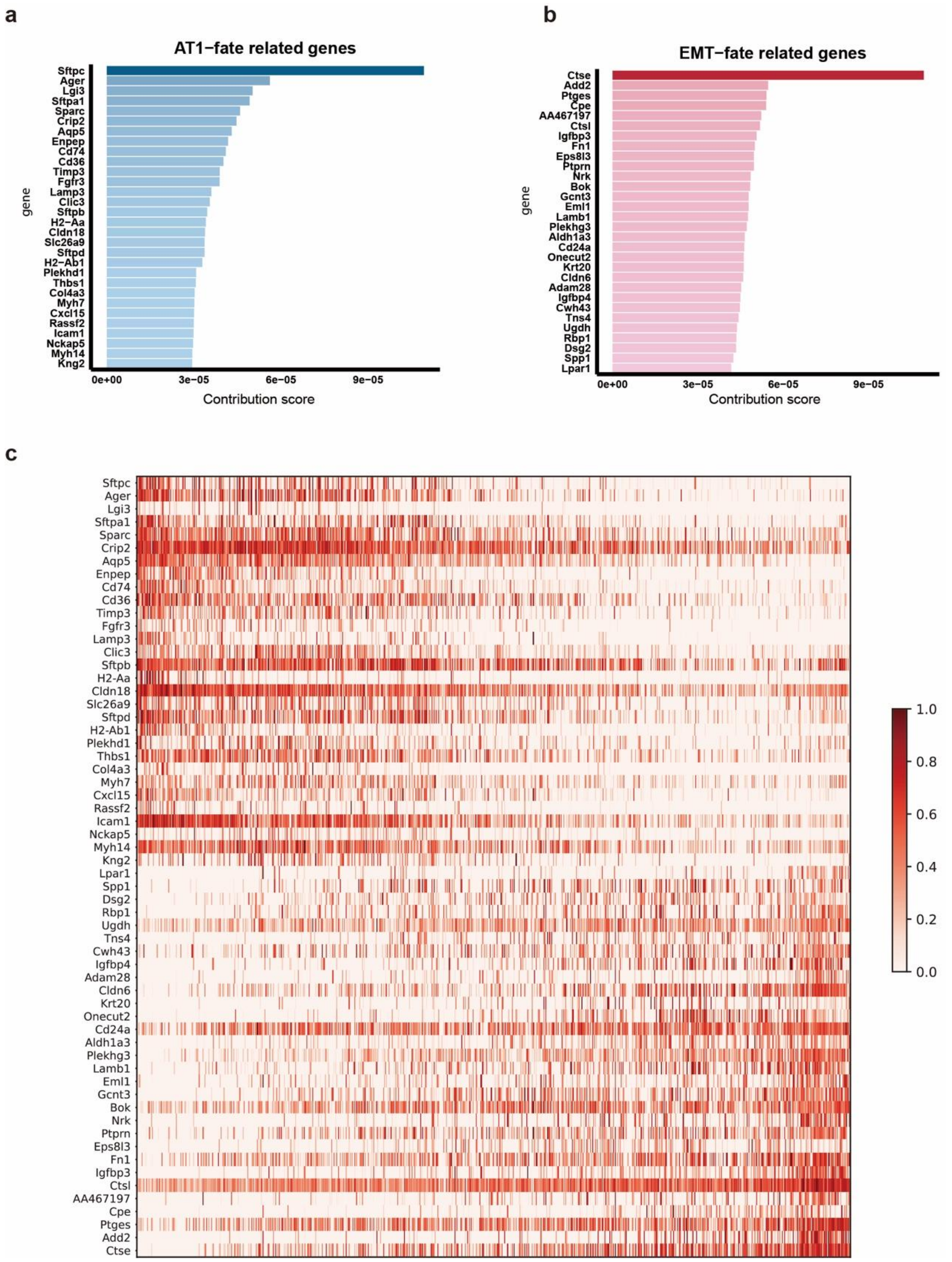


**Supplementary Fig. 12 Identification of lineage-specific genes for LUAD transitional state evolution toward AT1-like or EMT-like lineages.**

**(a-b)** Bar plots show the contribution scores of the top-20 genes specific to AT1-biased (left) and EMT-biased (right) fate potentials. **(c)** The heatmap shows the expression of the top-20 fate-specific genes toward AT1-biased and EMT-biased fate potentials in transitional cells. Each row denotes a fate-specific gene, the columns denote transitional cells arranged by increasing order of fate-bias calculated by fate potentials. Cells with fate-bias > 0.5 are EMT-biased, and those with bias < 0.5 are AT1-biased.

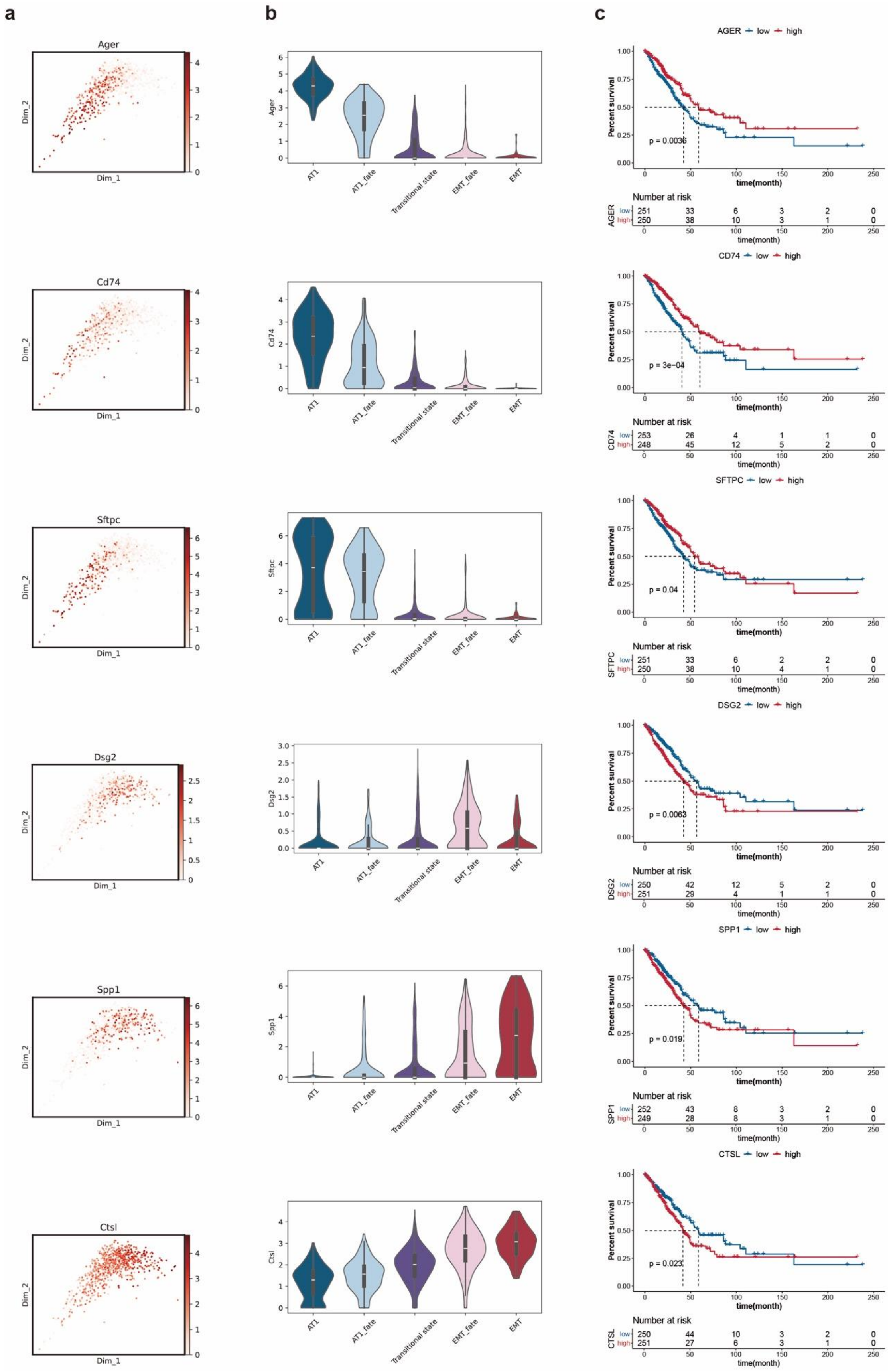


**Supplementary Fig. 13 Representative LUAD fate-specific genes and clinical associations**

**(a)** Transcriptomic expression level of fate-specific genes in the transitional state. **(b)** Violin plots show the gene expression level of fate-specific genes between LUAD tumor cell substates. **(c)** Fate-specific genes for overall survival from a TCGA-LUAD cohort.

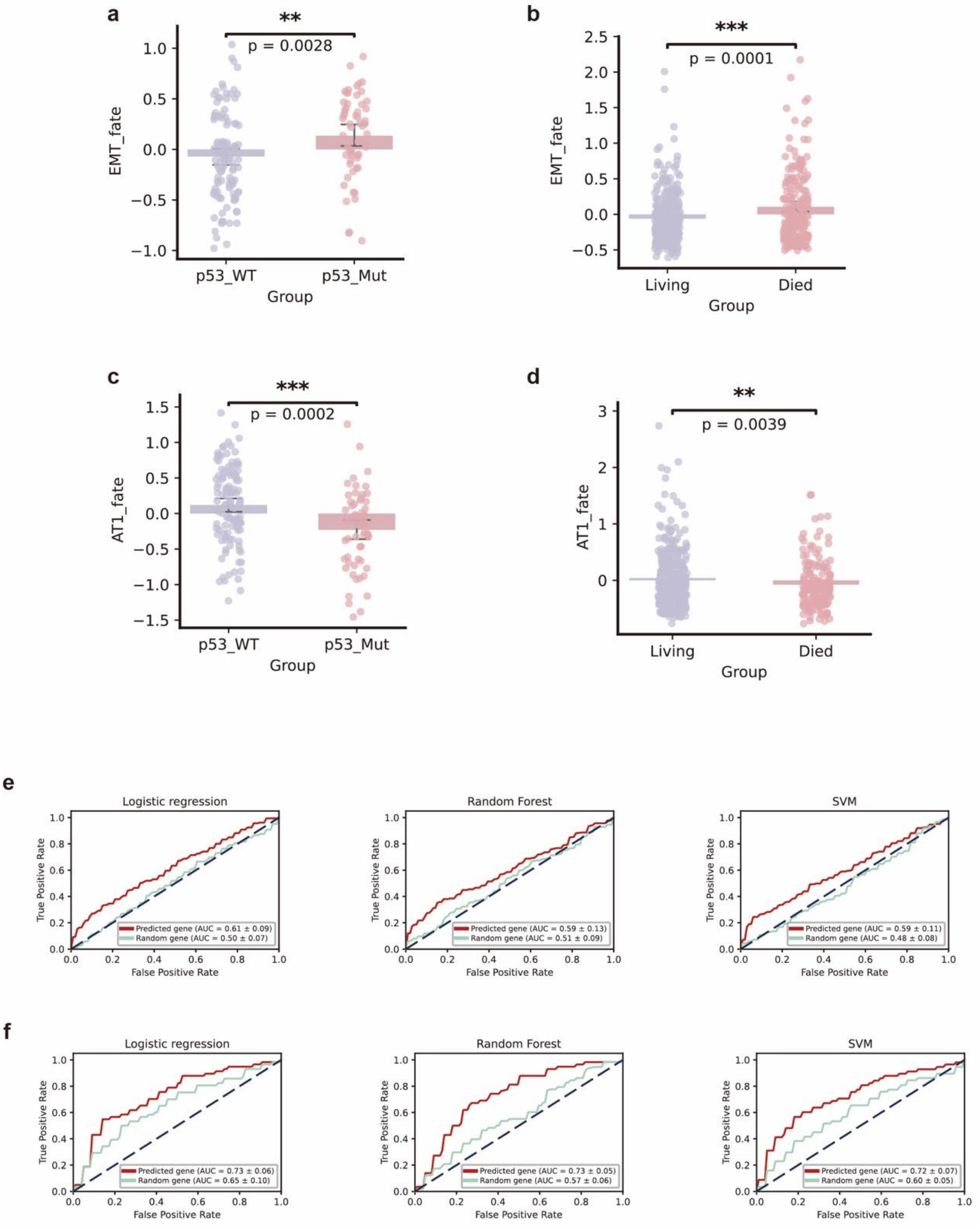


**Supplementary Fig. 14 Evaluation of fate-specific gene set for AT1-biased or EMT-biased fate potentials.**

**(a-b)** Box plots show the mean expression level of EMT-specific genes in a TCGA-LUAD cohort between LUAD patients grouped by p53 mutant status (left) and clinical outcomes (right). **(c-d)** Box plots show the mean expression level of AT1-specific genes in a TCGA-LUAD cohort between LUAD patients grouped by p53 mutant states (left) and survival states (right). **(e-f)** ROC curve analysis showing the predictive performance of fate-specific gene sets on TCGA-LUAD clinical outcomes (top) and TCGA-LUAD p53 mutant status (bottom).

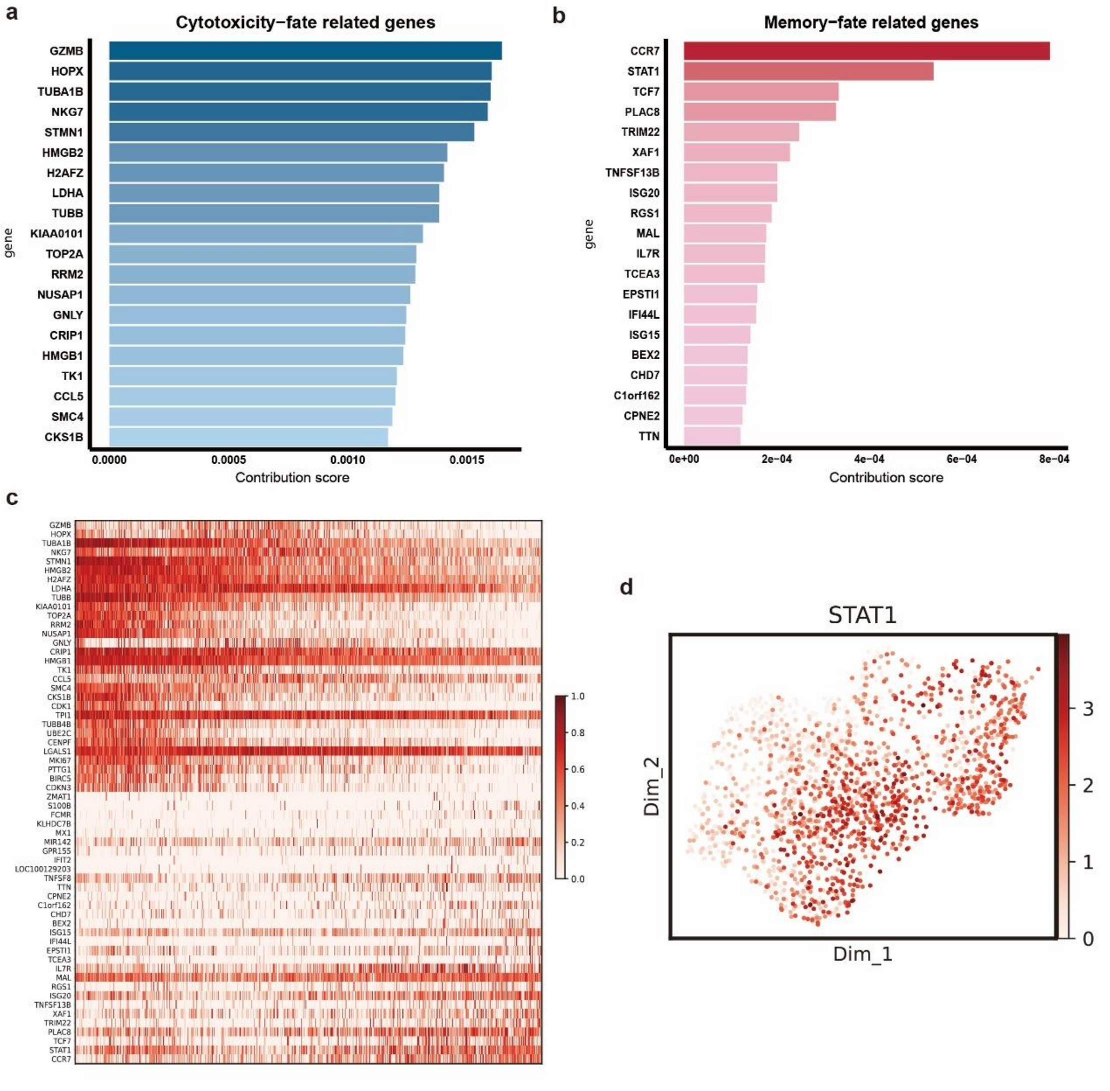


**Supplementary Fig. 15 Identification of lineage-specific genes for pre-infusion CAR-T cells toward cytotoxicity or memory-like lineages.**

**(a-b)** Bar plots show the contribution scores of the top-20 genes specific to cytotoxicity-biased (left) and memory -biased (right) fate potentials. **(c)** The heatmap shows the expression of the top-20 fate-specific genes toward cytotoxicity-biased and memory-biased fate potentials in pre-infusion CAR-T cells. Each row denotes a fate-specific gene, the columns denote pre-infusion CAR-T cells arranged by increasing order of fate-bias calculated by fate potentials. Cells with a fate-bias > 0.5 are memory-biased, and those with a bias < 0.5 are cytotoxicity-biased. **(d)** Gene expression level of STAT1 in Pre-infusion CAR-T cells.

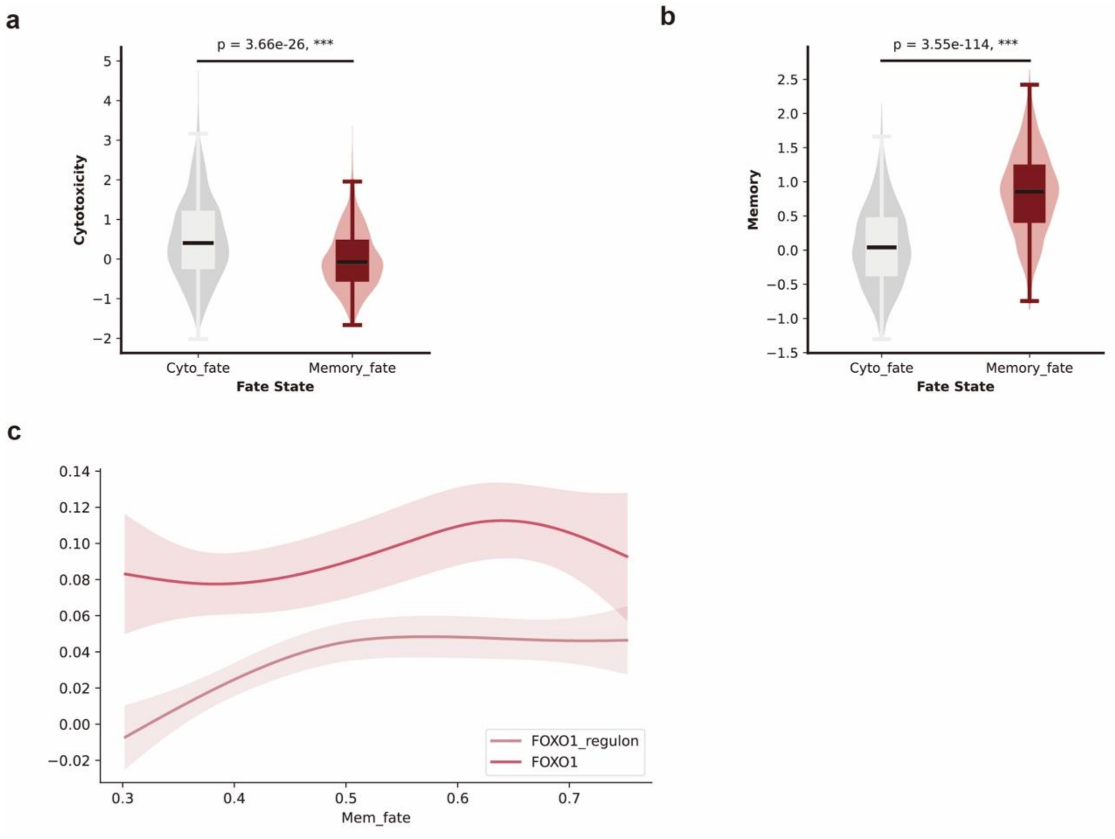


**Supplementary Fig. 16 Validation of fate-specific pre-infusion CAR-T states identified by DyMoTree.**

**(a-b)** Violin plots show the cytotoxicity (left) and memory (right) scores defined from the original study between fate-specific pre-infusion CAR-T states. **(c)** Aggregate score trends of FOXO1 regulon and transcriptomic expression trends of FOXO1 along memory-biased fate potential in pre-infusion CAR-T cells.

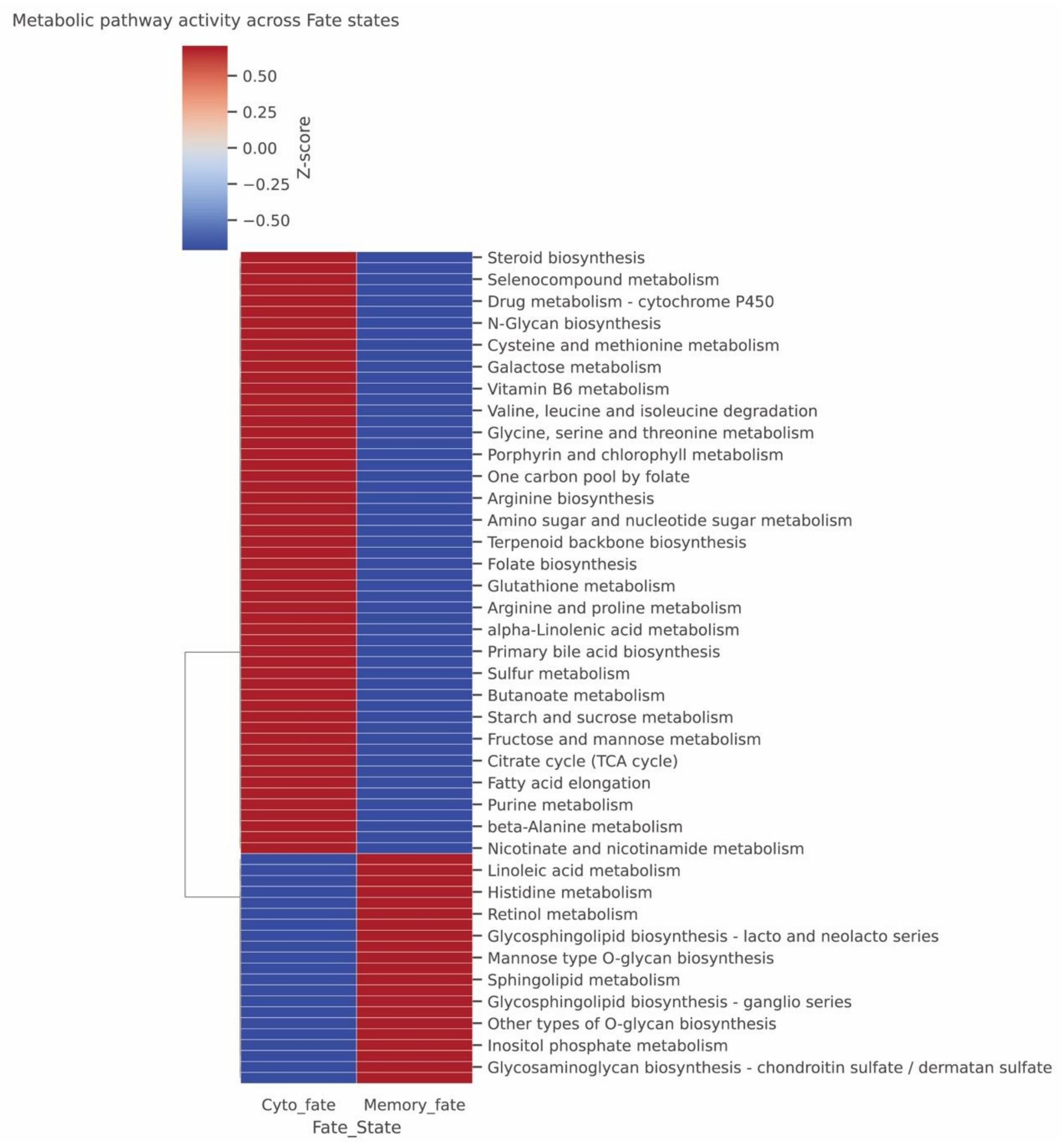


**Supplementary Fig. 17 The metabolic pathway activity heatmap for the pre-infusion CAR-T state.**

Each row denotes a specific metabolic pathway, and each column represents a fate-specific pre-infusion CAR-T state.

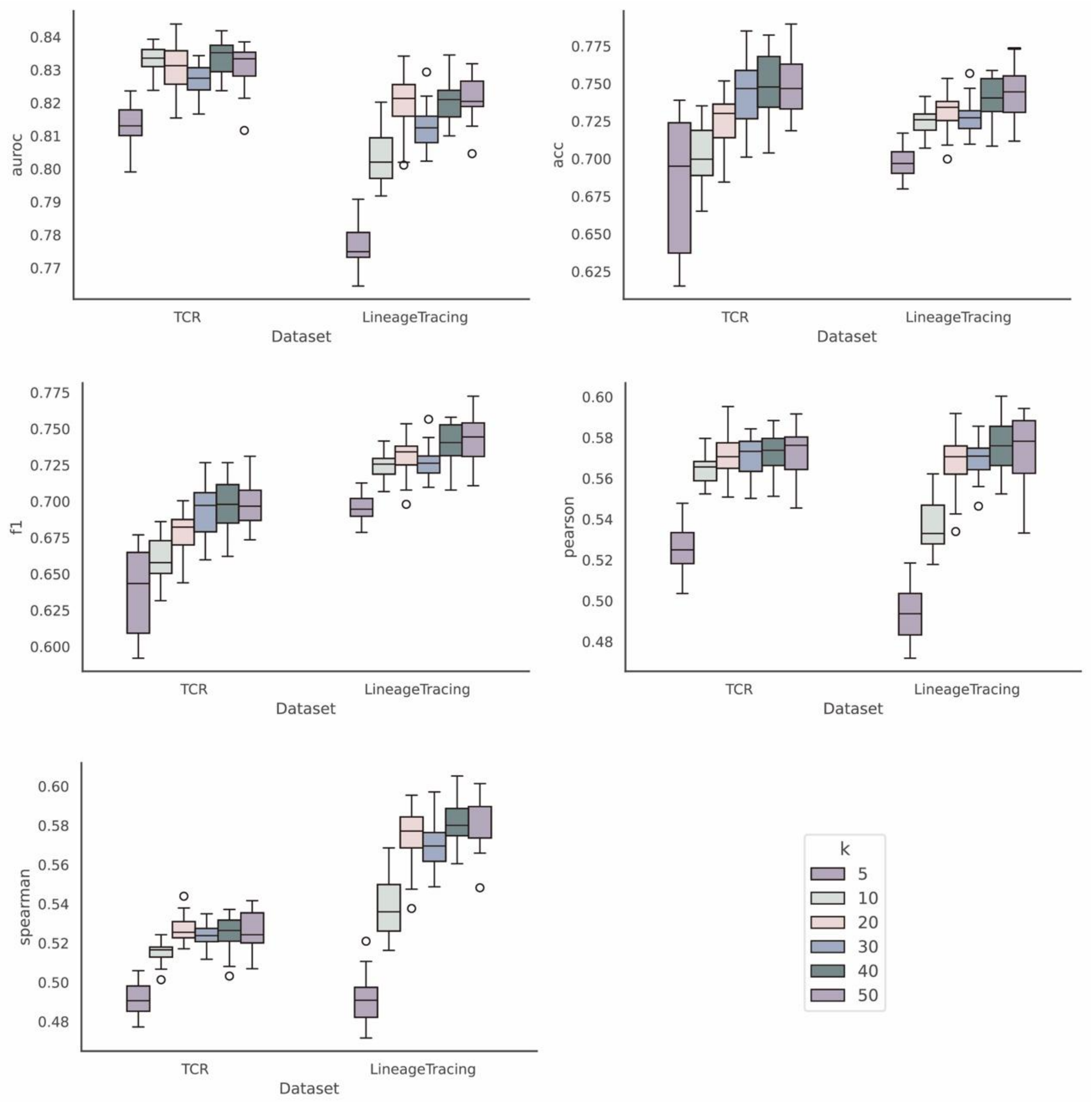


**Supplementary Fig. 18 Sensitivity of DyMoTree performance to the number of neighbors (k) in KNN graph construction.**

Performance comparison across different values of k (5–50) on TCR and lineage-tracing datasets, evaluated by AUROC, accuracy, F1-score, Pearson, and Spearman correlations.

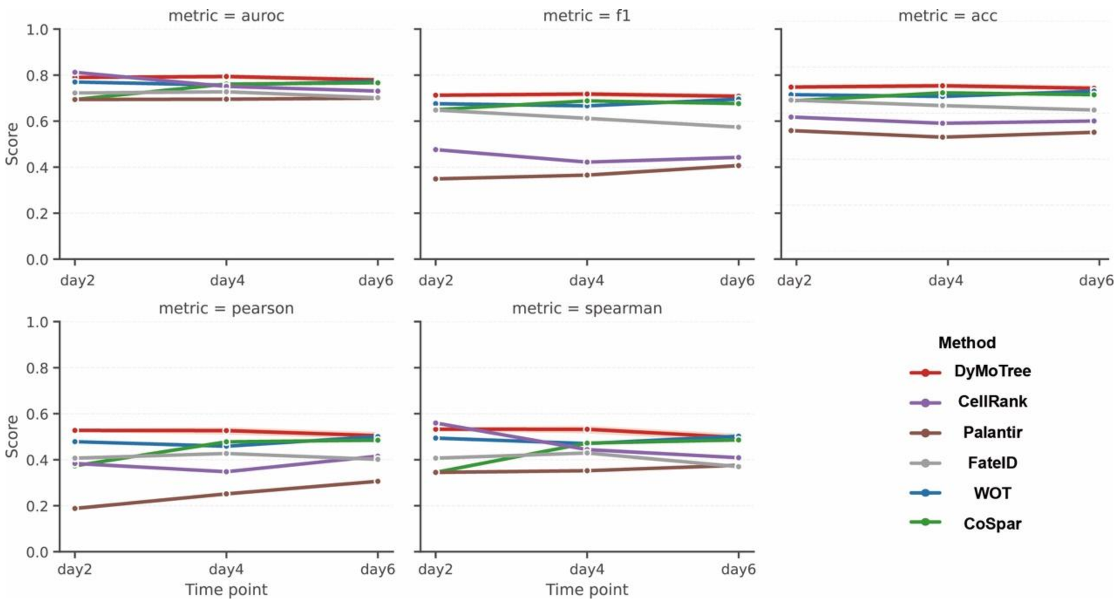


**Supplementary Fig. 19 Performance comparison across terminal cells at different time points.** Quantitative evaluation of DyMoTree and representative methods across day 2, day 4, and day 6 terminal cells in lineage tracing datasets using multiple metrics (AUROC, F1-score, accuracy, Pearson, and Spearman correlations).

**Supplementary Table 1 Gene sets used for functional analysis of the lung adenocarcinoma dataset.**

| Gene set | Gene |
|---|---|
| p53 signaling | Trp53, Phlda3, Prsc1, Ccng1, Zmat3 |
| metabolic processes | Serinc3, Tpi1, Rbp4, Vsig2, Rbp1 |
| cell proliferation | Ctse, Igfbp3, Gpa33, Slc4a8, Hnf4a |

**Supplementary Table 2 Hyperparameters used for DyMoTree model training.**

| Dataset | stage-1 learning rate | stage-1 iteration | stage-2 learning | stage-2 iteration | learning rate | iteration |
|---|---|---|---|---|---|---|
| LT (mixed) | 1.00E-03 | 100 | 1.00E-04 | 300 | 1.00E-04 | 300 |
| TCR | 1.00E-03 | 100 | 3.00E-04 | 500 | 1.00E-04 | 500 |
| ICM | 1.00E-03 | 100 | 1.00E-04 | 350 | 1.00E-04 | 350 |
| LUAD | 1.00E-03 | 100 | 1.00E-04 | 350 | 1.00E-04 | 350 |
| CAR-T | 1.00E-03 | 100 | 1.00E-03 | 200 | 1.00E-04 | 200 |
| LT (day2,4,6) | 1.00E-03 | 100 | 1.00E-03 | 300 | 1.00E-04 | 300 |

**Supplementary Table 3 Hyperparameters used for identifying fate-specific cell states.**

| Datasets | state number | n_pca | n_diff | n_gene |
|---|---|---|---|---|
| LT | 3 | 10 | 10 | 10 |
| ICM | 3 | 5 | 5 | 10 |
| LungCancer | 3 | 5 | 5 | 10 |
| CART | 2 | 10 | 10 | 20 |

**Supplementary Table 4 Hyperparameters used for identifying driver genes.**

| Datasets | top_n | soft_threshold | graph_threshold | lasso_alpha | method |
|---|---|---|---|---|---|
| ICM | 100 | 1 | 0 | 0.05 | pearson |
| LungCancer | 100 | 1 | 0 | 0.1 | spearman |
| CART | 200 | 1 | 0 | 0.01 | pearson |
